# A polyketide-based biosynthetic platform for diols, amino alcohols and hydroxyacids

**DOI:** 10.1101/2024.10.29.620908

**Authors:** Qingyun Dan, Yan Chiu, Namil Lee, Jose Henrique Pereira, Xixi Zhao, Chunjun Zhan, Yiou Rong, Yan Chen, Seokjung Cheong, Chenyi Li, Jennifer W. Gin, Andria Rodrigues, Tyler W. H. Backman, Edward E. K. Baidoo, Christopher J. Petzold, Paul D. Adams, Jay D. Keasling

## Abstract

Medium- and branched-chain diols and amino alcohols are important industrial solvents, polymer building blocks, cosmetics and pharmaceutical ingredients, yet biosynthetically challenging to produce. Here, we present a novel approach utilising a modular polyketide synthase (PKS) platform for the efficient production of these compounds. This platform takes advantage of a versatile loading module from the rimocidin PKS and NADPH-dependent terminal thioreductases (TRs), previously untapped in engineered PKSs. Reduction of the terminal aldehyde with specific alcohol dehydrogenases enables production of diols, oxidation enables production of hydroxy acids, and transamination with specific transaminases enables production of various amino alcohols. Furthermore, replacement of the malonyl-coenzyme A (CoA)–specific acyltransferase (AT) in the extension module with methyl- or ethylmalonyl-CoA–specific ATs enables production of branched-chain diols and amino alcohols. In total, we demonstrated production of nine 1,3-diols (including the difficult-to-produce insect repellent and cosmetic ingredient 2-ethyl-1,3-hexanediol), six amino alcohols, and two carboxylic acids using our PKS platform in *Streptomyces albus*. Finally, tuning production of the PKS acyl-CoA substrates enabled production of high titers of specific diols and amino alcohols (1 g/L diol titer in shake flasks), demonstrating high tunability and efficiency of the platform.

## Main

Diols are industrially important commodity and specialty chemicals widely used today^1^. Over the last decades, a few natural and artificial pathways have been established in *Escherichia coli* and other industrial microbes targeting different short-chain diols^2–5^. The development and industrialization of microorganisms to convert renewable feedstocks into 1,3-propanediol, 1,4- butanediol, and 1,3-butanediol (1,3-BDO or butylene glycol, used as a humectant and cosmetic solvent) has been a major advance for metabolic engineering^6,7^. Besides short-chain diols, medium-chain and branched-chain diols are also important solvents, polymer building blocks, fragrances and cosmetics ingredients^1,8^. A prime example is 2-ethyl-1,3-hexanediol (2-E-1,3- HDO or etohexadiol), an insect repellent, boron extractant, ink solvent and ingredient in cosmetics used for over half a century^9^. Besides its own market use, its derivative 2-ethylhexanol is a major building block for polyvinyl chloride plasticisers, is produced in millions of metric tons annually, and had a six billion USD market size in 2022^10^. Despite their importance, metabolic pathways to produce medium- and branched-chain diols are rare and have not been commercialised. Also, each molecule generally requires a distinct biosynthetic pathway that will not produce other analogs, making it impossible to extend learnings from synthesis of one molecule to others. In fact, this biosynthetic challenge is widely present in almost all medium- and branched-chain chemical bioproduction (alcohols, amines, amino alcohols, to list a few), presenting a major knowledge gap in metabolic engineering. To tackle this problem, we aimed to design a biosynthetic platform suitable for production of 1,3-BDO, 2-E-1,3-HDO, or nearly any medium- or branched-chain alcohol and derivative, with a key feature of programmable access to precisely altered chemistry.

Modular type I polyketide synthases (PKSs) are megasynthases that utilise coenzyme A (CoA) substrates to produce complex natural products, many of which are used as pharmaceuticals^11^. The chemical structure of PKS products are strictly determined by the order of each enzymatic domain and module. Because the structure of the molecule it produces is encoded in the DNA sequence, scientists have long dreamed of designing PKSs to produce nearly any organic molecule^12,13^. And recently, PKSs have been redesigned to incorporate unnatural functionalities, such as fluorination^14,15^ or to produce new-to-nature molecules, such as short-chain ketones^16^ and triketide lactones^17,18^. Thus, we hoped to investigate if PKSs can be used as a platform to biosynthesize many different diols and related molecules that have not been biosynthesized or molecules that require distinct biosynthetic pathways. For PKS production of 1,3-BDO and 2-E-1,3-HDO, two prerequisites are an initiation module that selects for acetyl- and butyryl- starter units and an extension module with ketosynthase-acyltransferase- ketoreductase-acyl carrier protein (KS-AT-KR-ACP) architecture to install an OH on the third carbon (**Fig. 1a**). Finally, we must terminate polyketide biosynthesis with an alcohol if we want to produce a diol with an alcohol on the first carbon (or with an amine if we want to produce an amino alcohol with an amine on the first carbon); unfortunately, most natural PKSs terminate polyketide biosynthesis with thioesterases (TEs), which either directly hydrolyze the ACP intermediates to produce carboxylic acids or catalyse macrocyclization to give lactones or lactams^19^. Indeed, the monotony of TE-mediated termination is a major hurdle for expanding current PKS design space.

**Figure 1.**
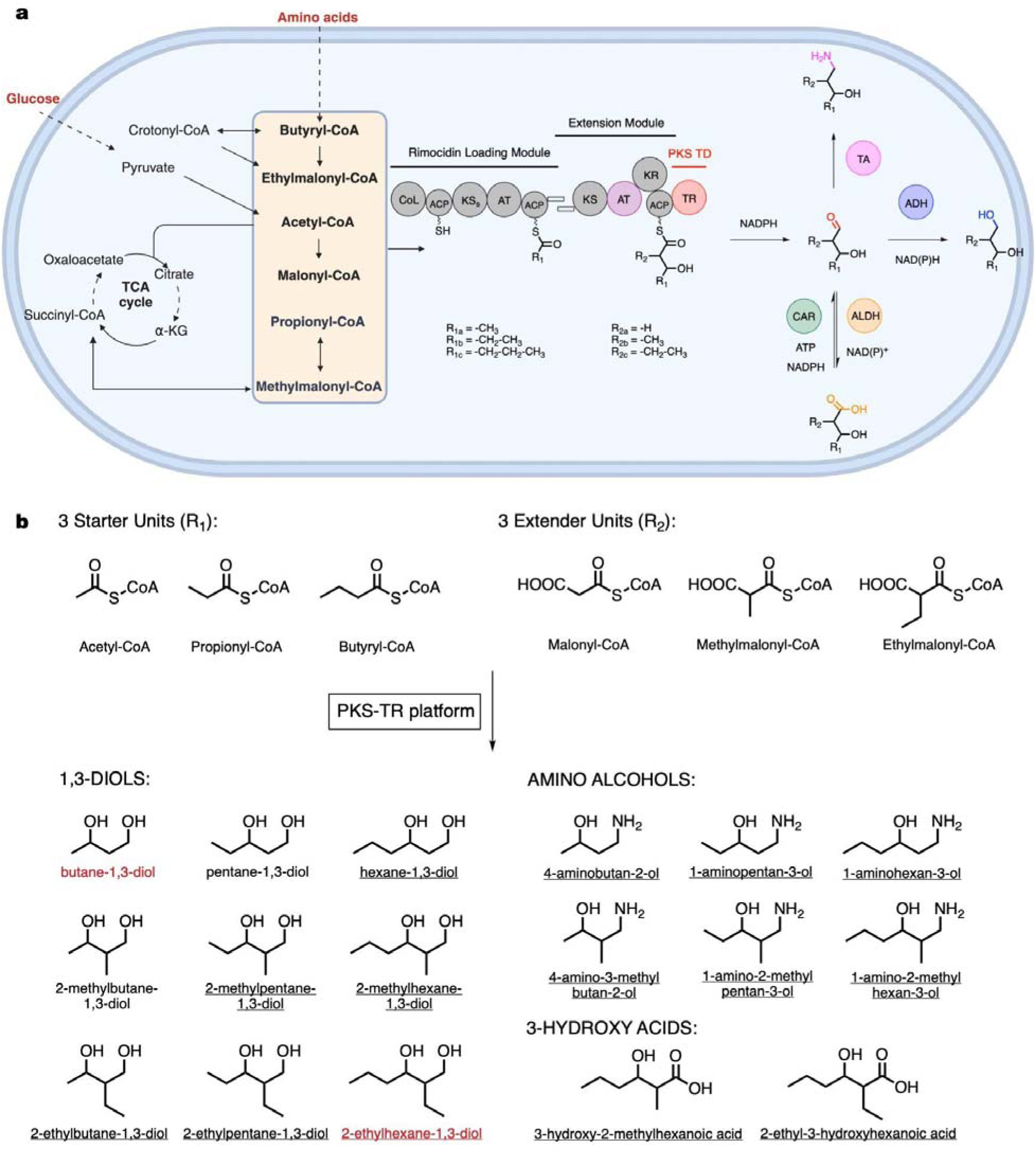
Schematic representation of the rimocidin PKS-TR platform in *Streptomyces*. a, Using glucose (and L-valine) as the carbon source, engineered rimocidin PKS can load three CoA starter units and three extender units. Incorporation of a PKS TR led to production of diverse products via programmed post-PKS modification by alcohol dehydrogenases (ADHs), aldehyde dehydrogenases (ALDHs) or transaminases (TAs). Dashed lines indicate multiple steps in the precursor pathways. TD: terminal domain; CoL: CoA ligase; ACP: acyl carrier protein; KS: ketosynthase; AT: acyltransferase; KR: ketoreductase; TR: thioreductase; ADH: alcohol dehydrogenase; ALDH: aldehyde dehydrogenase; CAR: carboxylic acid reductase; TA: transaminase. b, Summary of all products in this study. Molecules highlighted in red are industrially valuable, and underlined products were not microbially synthesised before.

One appealing alternative to overcome this challenge is the use of terminal thioreductases (TRs)^13^. PKS chain release catalysed by TRs was revealed in the recent decade, with only a handful of examples^20–26^, including the coelimycin and cyclizidine PKSs. They are often coupled with cognate transaminases (TAs) in the biosynthetic gene clusters (BGCs): the TR catalyses reductive cleavage of the acyl substrate to produce an aldehyde, which is subsequently converted to amine by the TA^20^. Apart from the few studies to understand natural PKS TRs, they have not been used in engineered PKSs. Here, we biochemically and structurally characterised PKS TRs and confirmed that TRs catalyse two electron reductive cleavage and produce aldehydes. Based on the fact that aldehydes are highly reactive intracellularly, we designed an engineered PKS- TR-based biosynthetic platform for bioproduction of a wide range of aldehyde derivatives, illustrated by production of nine 1,3-diols including 1,3-BDO and 2-E-1,3-HDO, six amino alcohols, and at least two 3-hydroxy acids, fourteen of which have not been previously biosynthesized (**Fig. 1b**). This platform utilises PKS substrate promiscuity and assembly-line architecture to provide designable carbon skeletons, and TR catalysis and programmed post-PKS modification to diversify C1 chemistry, with potential to produce a wide range of bioproducts including diols, alcohols, amines, and hydrocarbon biofuels.

## Results

### PKS TRs terminate polyketides with an aldehyde group

Comprehensive PKS TR engineering was not previously reported, thus we first built a PKS TR library based on phylogeny analysis (**Fig. 2a and Supplementary** Figure 1) and selected nine TRs to study (TR1-9). The selected TRs uniformly covered the phylogenetic tree, and seven of them had validated pathway products. Among them, the TR from the coelimycin PKS BGC (TR1 or CpkC TR) was reported to be an NADH-dependent reductase^20^, capable of reducing octanoyl-CoA, a natural substrate mimic (**Supplementary** Figure 2), to octanol in two steps (**Fig. 2b**). Our phylogenetic analysis suggested a close evolutionary relationship between PKS TRs and NADPH-dependent PKS ketoreductases (KRs)^27^ and a distant relationship with NADH-dependent alcohol dehydrogenases^28^ (ADHs), disagreeing with this NADH-dependent literature report (**Fig. 2a**). To gain further insight into TR catalysis, we overexpressed TR1, TR2, TR7, and TR9 in *E. coli* BL21(DE3) and purified these TRs to > 95% homogeneity (**Supplementary** Figure 3a-3d). We also purified the TR1-cognate ACP1 with a maltose binding protein (MBP) tag fused at the N-terminus of ACP1 to improve protein solubility (**Supplementary** Figure 3e). We then loaded octanoyl-CoA onto MBP-ACP1 *in vitro* with Sfp, a promiscuous phosphopantetheinyl transferase^29^. After removal of excess octanoyl-CoA via dialysis, octanoyl- ACP1 was tested as a TR1 substrate together with octanoyl-CoA and octanal. TR1 did not reduce octanoyl-CoA or octanal when NADH was used as cofactor (**Supplementary** Figure 4b) but it did reduce octanoyl-ACP1. With NADPH, TR1 reduced octanoyl-ACP1 and octanoyl-CoA at a much faster rate than octanal (**Supplementary** Figure 4a). These results indicate that TR1 is an NADPH-preferred reductase. To conclusively address substrate scope and cofactor preference of PKS TRs, we next purified ACP9 without the MBP tag and tested reactivity with TR9 (**Supplementary** Figure 3f). TR9 is from the venediol BGC, with 63% sequence identity to TR1^25^. TR9 only used NADPH as cofactor, and readily reacted with the natural substrate analog octanoyl-ACP9 or octanoyl-CoA but would not react with NADH or octanal (**Fig. 2c-2d and Supplementary** Figure 4c). From these data, we concluded that TRs are NADPH-dependent termination enzymes that produce aldehydes and are incapable of further reducing aldehydes to alcohols.

**Figure 2.**
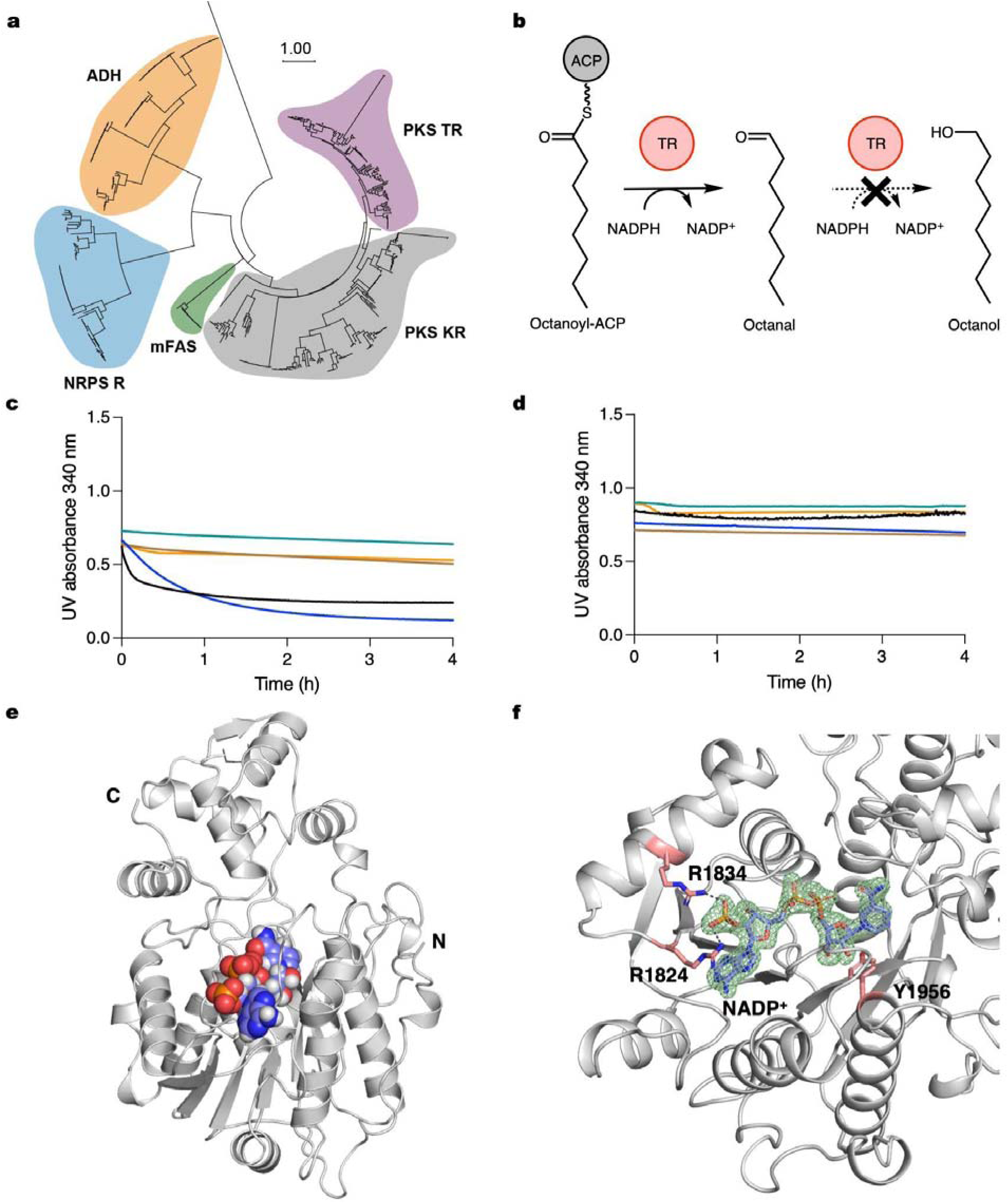
PKS TR phylogeny, catalysis, and crystal structures. a, Phylogenetic analysis of PKS TRs in purple, PKS KRs in grey, mammalian fatty acid synthases (mFASs) in green, NRPS R domains in cyan, and alcohol dehydrogenases (ADHs) in primary metabolism in orange. b, TR9-catalysed reaction scheme of a natural substrate mimic, octanoyl-ACP9. c-d, Characterization of TR9 substrate scope and cofactor preference, with NADPH in c and NADH in d. NAD(P)H consumption was assessed by monitoring UV absorbance at 340 nm. Black: TR9 + NAD(P)H + octanoyl-ACP9; Blue: TR9 + NAD(P)H + octanoyl-CoA; Brown: TR9 + NAD(P)H + octanal; Green: NAD(P)H + octanoyl-CoA; Orange: NAD(P)H + octanal. For TR catalysis, the preferred substrates are ACPs or CoAs, and the preferred cofactor is NADPH. e, CpkC TR (TR1) crystal structure bound with NADP^+^, with protein coloured in white and cofactor coloured in blue. f, Active site view of TR1, R1824 and R1834 coordinate NADP^+^ ribose 2’-phosphate, Y1956 is the catalytic proton donor. The NADP^+^ omit map (mFo-DFc, 3.0 σ level) is coloured in green.

To understand the structural basis for TR cofactor recognition, we determined the crystal structure of CpkC TR (TR1) bound with NADP^+^ to 1.8 Å resolution (**Fig. 2e**). The TR1 structure consists of the N-terminal nucleotide binding domain and the C-terminal substrate binding domain, featuring an invariant “TGX_2_GX_2_G” motif, and two conserved arginines at 1824 and 1834 coordinating the ribose 2’-phosphate in NADP^+^, providing structural explanation for NADP^+^ preference^28,30^ (**Fig. 2f and Supplementary** Figure 5). Overall, these structural features align well with published tyrosine-dependent “extended” short-chain reductase/dehydrogenase (SDR) structures^31,32^. We performed structural homology search with Dali server^33^ and found the closest structural homologs to be nonribosomal peptide synthetase MxaA terminal reductase (NRPS R) domain^31^ with root mean squared deviation (r.m.s.d.) of 1.14 Å, and carboxylic acid reductase (SrCAR) R domain^34^ with r.m.s.d. of 1.97 Å. The major structural differences between PKS TR, NRPS R and CAR R lie in the post-β5 loop and α10-α11 “helix-turn-helix” (HTH) motif (**Supplementary** Figure 6), which are the putative ACP or peptidyl carrier protein (PCP) binding sites. The HTH motif is absent in other common types of SDRs, supporting the importance of HTH for ACP/PCP recognition^35^. Furthermore, the limited structural similarity between PKS TR, NRPS R and CAR R HTH regions suggest distinct substrate binding patterns for these three types of reductases in secondary metabolism.

### Engineering rimocidin PKS and TRs for 1,3-diol production

With a functional TR in hand and an understanding of its catalytic potential, we next sought to use it to produce aldehyde-derived alcohols, including 2-E-1,3-HDO. We used retrobiosynthesis software, ClusterCAD RetroTide, which we developed previously to design PKSs for specific products^36^. We designed PKS domain architectures for 2-E-1,3-HDO production, which contains a butyryl-CoA-specific loading module, an ethylmalonyl-CoA-specific extension module with KS-AT-KR-ACP architecture, and a TR termination domain. Based on that information, we searched for suitable PKS candidates and narrowed our candidates to the rimocidin (Rim) PKS. Rimocidin-like natural products have been isolated from several *Streptomyces* species^37,38^. Depending on which substrates are loaded onto the PKS, the products are CE-108 and rimocidin in *Streptomyces diastaticus* var. 108 and BU16 and rimocidin in *Streptomyces mauvecolor* strain BU16. Only partial BGC information was available, thus we analysed the Rim BGCs and validated their PKS architecture with antiSMASH^39^ **(Supplementary** Figure 7). Since the Rim PKS loading module RimM0 has a novel CoA ligase loading domain architecture, (CoL)-ACP1-KS-AT-ACP2, we investigated its loading mechanism as the first step. However, we failed to obtain soluble RimM0 when we expressed it in *E. coli*. Instead, we turned our attention to a RimM0 homolog protein in the natamycin/pimaricin (Pim) PKS pathway, PimS0, which has the same domain architecture^40^ (**Supplementary** Figure 8a). We expressed the gene encoding PimS0 in *E. coli*, purified it, and confirmed this novel PKS initiation mode with an acetyl- starter unit (**Supplementary** Figures 8b-8c and 9). Despite the same initiation module architecture as Rim PKS, Pim PKS is unable to load the butyryl- start unit required for 2-E-1,3-HDO production. Thus, we tested whether the Rim PKS can be fused with TRs and produce 1,3-diols in a microbial host (**Fig. 3a**).

**Figure 3.**
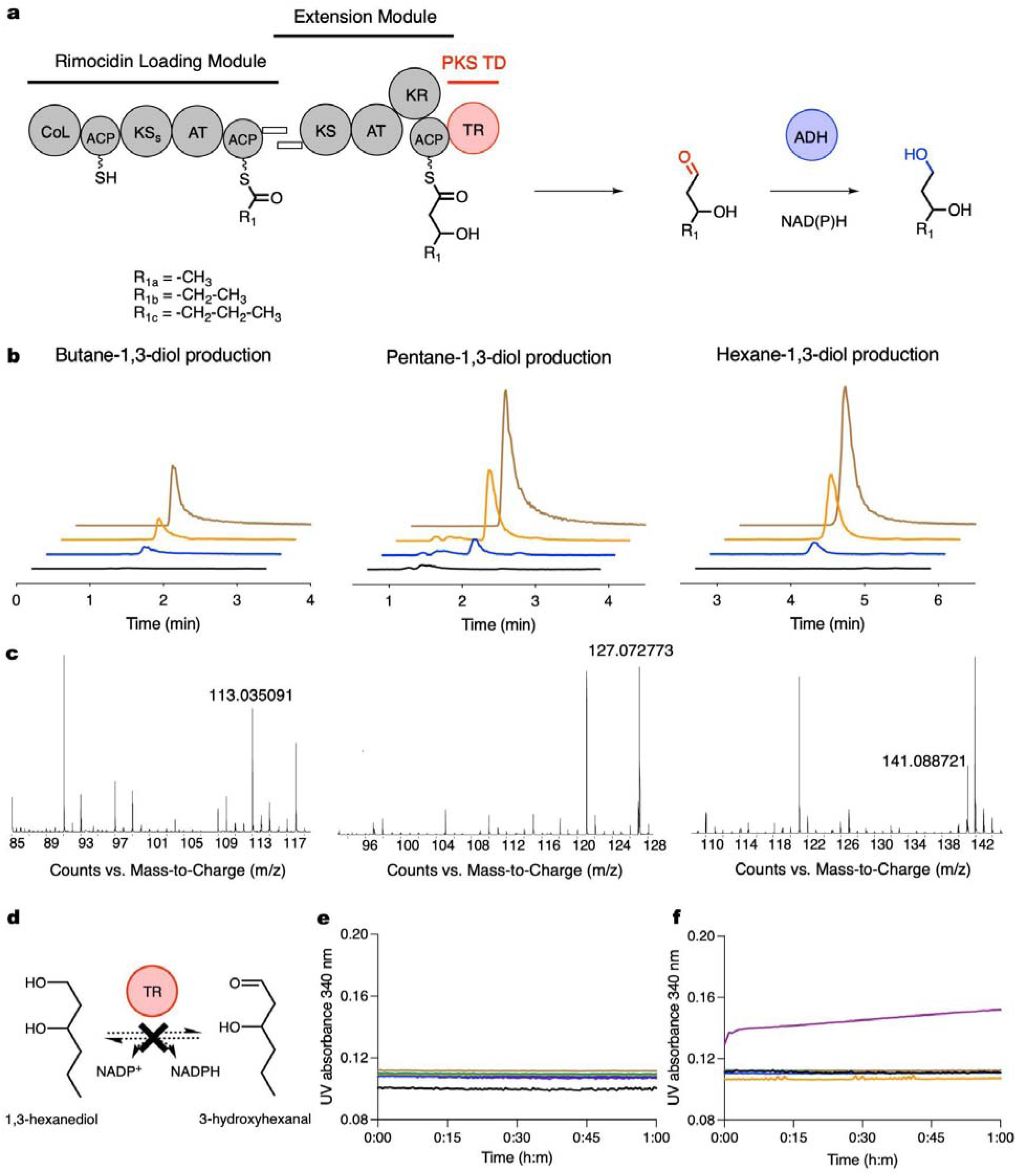
Rimocidin PKS- and TR-based 1,3-diol bioproduction. a, Scheme of RimPKS-TR for production of 1,3-BDO, 1,3-PDO, and 1,3-HDO. b, LC-MS extracted ion chromatograms (EICs) of 1,3-diols produced by *S. albus* RimM0M1-TR1 (QD27). Black: *S. albus* RimM0 (QD1) control; Blue: QD27; Orange: QD27 spiked with standards; Brown: 1,3-diol standards. c, 1,3- diol [M + Na]^+^ *m/z* detection, see Supplementary Figure 12 for detailed information. d-f, Characterization of 1,3-HDO enzymatic oxidation. NADPH (e) or NADH (f) accumulation was assessed by monitoring UV absorbance at 340 nm. Black: 1,3-HDO + NAD(P)^+^; Blue: TR1 + 1,3-HDO + NAD(P)^+^; Brown: TR2 + 1,3-HDO + NAD(P)^+^; Green: TR7 + 1,3-HDO + NAD(P)^+^; Orange: TR9 + 1,3-HDO + NAD(P)^+^; Purple: TADH2 + 1,3-HDO + NAD(P)^+^. No TR catalysed such oxidation reactions. TADH2 serves as a positive control.

Using bacteriophage integrases^41^, we integrated the native *rimM0* gene driven by a constitutive *PrpsL(RO)* promoter into *Streptomyces albus* J1074 and *Streptomyces coelicolor* M1152, two common heterologous hosts for PKS engineering. RimM0 protein abundance in these *Streptomyces* hosts was assessed by proteomics analysis, which showed that RimM0 was present at 1.2 ± 0.3% relative protein abundance in *S. albus* and absent in *S. coelicolor* (**Supplementary** Figure 10a). We also screened *rimM0* expression under two other constitutive promoters, *Pgapdh(EL)* and *kasOP*\*^42,43^. *Pgapdh(EL)*-driven *rimM0* expression peaked the screening test, with 8.3 ± 0.9% protein abundance, while *kasOP*\*-driven *rimM0* expression was not detected. With *S. albus* RimM0 (QD1) in hand, we examined potential target molecule consumption by our *Streptomyces* hosts, a phenomenon that we have observed when engineering other hosts to produce new-to-nature molecules^44,45^. Neither of the two target molecules, 1,3- pentanediol (1,3-PDO) and 1,3-hexanediol (1,3-HDO), were catabolized by our *Streptomyces* hosts when added to cultures (**Supplementary** Figure 11). This drove us to build *S. albus* RimM0M1-TR1 (QD27) and look for 1,3-diol production. For the chimeric RimM1-TR1 design, we identified the N-terminal boundary of TR1 based on the crystal structure, fused the N-terminus of TR1 with the C-terminus of RimM1 ACP, and integrated the chimeric *Pgapdh(EL)*- *rimM1-TR1* into QD1, making QD27. LC-MS analysis was used for detection of 1,3-diol products. We successfully detected production of 6.7 mg/L 1,3-BDO, 12.3 mg/L 1,3-PDO, and 0.76 mg/L 1,3-HDO in TSB medium after 72 hours, demonstrating the feasibility of PKS-TR engineering for diol bioproduction for the first time (**Fig. 3b-3c, Supplementary** Figure 12). In comparison, we also constructed *S. albus* RimM0M1-DEBS TE (QD18) and looked for 3- hydroxy acid production, but with no success (**Supplementary** Figure 13a). 3- Hydroxypentanoic acid and 3-hydroxyhexanoic acid, two putative QD18 products, were quickly consumed by *S. albus* J1074 after direct feeding, providing a possible explanation (**Supplementary** Figure 13b-13c). This presents a vivid example of PKS-TR advantage over traditional PKS-TE engineering, as polyketide-based carboxylic acids may be prone to β- oxidation, resulting in loss of desired product and ultimately design failure. Apart from providing alternative C1 chemistry as more reactive aldehyde, PKS-TR may also protect the designed carbon skeleton from host consumption and degradation.

Because TR is incapable of aldehyde reduction, the aldehyde to 1,3-diol conversion that occurred in our engineered *S. albus* QD27 was likely catalysed by unknown ADHs in the host. To confirm this hypothesis, we tested whether four purified TRs can oxidise the 1,3-diols that QD27 produced to 3-hydroxy aldehydes, as ADH-catalysed reactions between alcohols and aldehydes are usually reversible (**Fig. 3d**). For example, promiscuous tomato alcohol dehydrogenase 2 (TADH2) readily catalysed NAD^+^-dependent oxidation on 1,3-BDO, 1,3-PDO, or 1,3-HDO^46^ (**Fig. 3e-3f and Supplementary** Figures 14-15). In comparison, none of the four TRs accepted these 1,3-diols as the substrate, suggesting that native *S. albus* ADHs catalyse the final reduction step in 1,3-diol biosynthesis without the need to incorporate exogenous ADHs (**Fig. 3e-3f**).

### Exploring engineering strategies of a tunable PKS-TR platform

Our next goal was to increase the titers of all 1,3-diols and also ratio of 1,3-HDO to other diols. We first screened for a series of common *Streptomyces* growth media including TSB, R5, M042, ISP2, and ISP4 (**Supplementary** Figure 16). When grown in R5 medium, QD27 produced the highest titers of 1,3-diols in 72 hours (38.6 mg/L). Based on this result, for the following experiments, we chose R5 as the standard medium, which includes 1% glucose. Next, we tested eight RimM1-TR2/3/4/6/7/8/9 chimaeras and confirmed 1,3-diol production in all designs except for RimM1-TR4 and RimM1-TR7. From the six successful designs, RimM0M1-TR2 (QD28) was the best 1,3-diol producer (**Fig. 4a**). Finally, prolonged 10-day cultivation of QD28 led to an increase of 1,3-diol titers to 264 mg/L, with 135.4 mg/L 1,3-BDO, 99.7 mg/L 1,3-PDO, and 29.1 mg/L 1,3-HDO produced in shake flasks (**Fig. 4b**). Interestingly, although *Pgapdh(EL)*-driven *rimM0* expression was higher than *PrpsL(RO)*-*rimM0* expression in engineered *S. albus*, 1,3-diol production in the latter strain was optimal (**Supplementary** Figure 10b).

**Figure 4.**
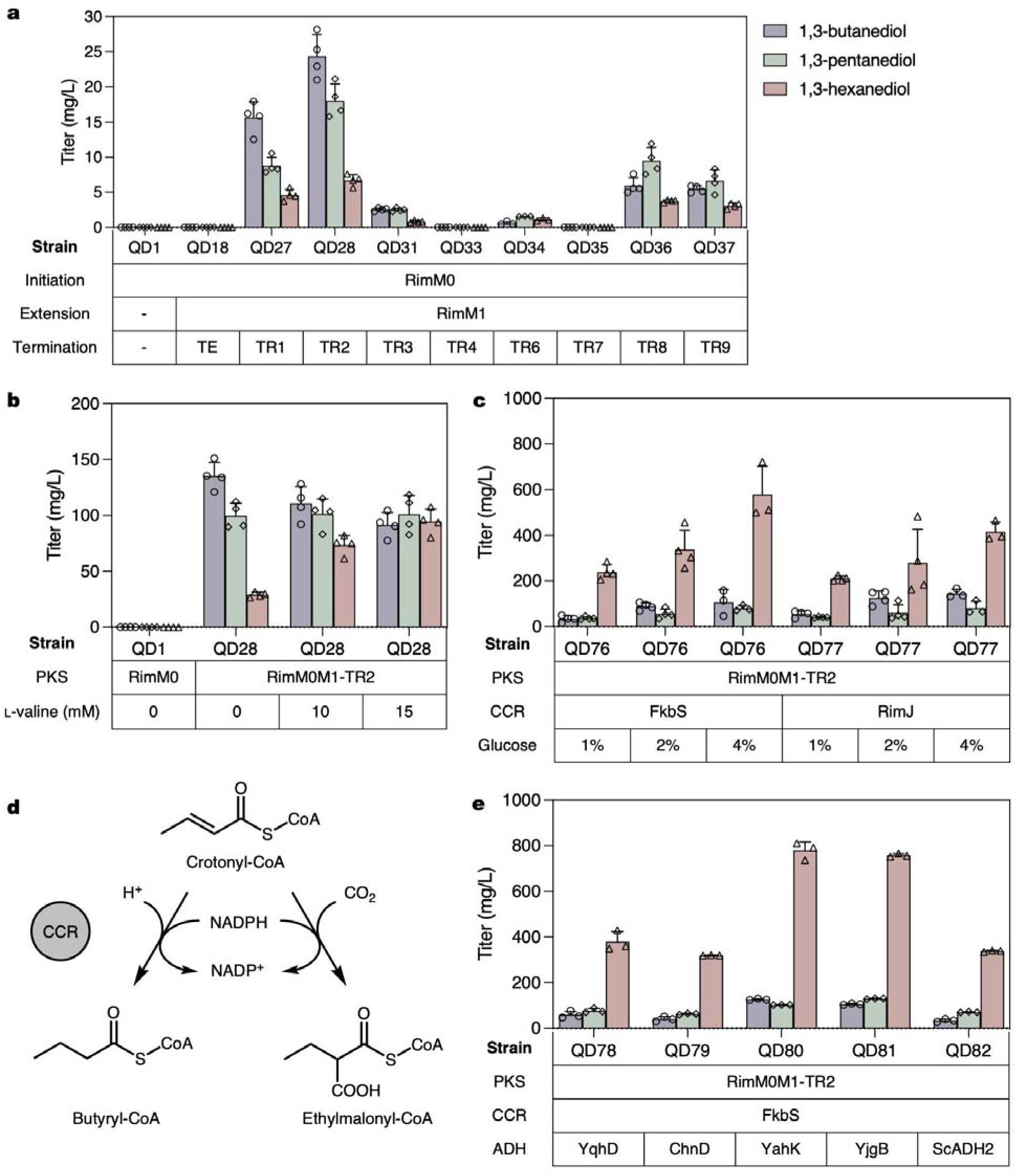
**Improving 1,3-diol production in *Streptomyces albus* J1074 RimPKS-TR**, with 1,3- BDO coloured in purple, 1,3-PDO in green, and 1,3-HDO in red. a, RimM0M1-TR 3 d cultivation results for screening TRs in R5 medium. b, Addition of butyryl-CoA precursor, L- valine, increased 1,3-HDO production titer and ratio in RimM0M1-TR2 (QD28). c, Overexpression of FkbS or RimJ crotonyl-CoA reductase (CCR) in QD28 and increasing glucose concentration in R5 led to 765 mg/L total 1,3-diol production in shake flasks. d, CCR-catalysed reactions to convert crotonyl-CoA to butyryl-/ethylmalonyl-CoA. e, Overexpression of an exogenous alcohol dehydrogenase (ADH) YahK in engineered *S. albus* RimM0M1-TR2 + FkbS (QD76) led to QD80 with increased 1,3-diol total titers of 1008 mg/L after 7 d cultivation in R5 + 2% glucose + 15 mM L-valine. All cultivation data are represented as mean value and error bars indicate standard deviation (s.d.) of at least three biological replicates.

Because 1,3-HDO was still a minor product in QD28, we hoped to further improve its titer and product ratio. Substrate-promiscuous PKSs have been proposed as biosynthetic platforms elegantly tuned by their CoA substrate regulation, with a few successful engineering reports^16^. In *S. albus*, addition of L-valine increased intracellular butyryl-CoA and ethylmalonyl- CoA levels^47^, both of which are putative building blocks for 1,3-HDO and 2-E-1,3-HDO biosynthesis. To validate the hypothesis that L-valine was converted to 1,3-HDO involving engineered RimPKS-TR catalysis, we fed in [^13^C]labelled L-valine and observed [^13^C]labelled 1,3-HDO as the final product (**Supplementary** Figure 17). We then added 15 mM L-valine in R5 medium and detected a slight increase of overall 1,3-diol titers from 264.2 mg/L to 287.2 mg/L, while featuring significant improvement of both 1,3-HDO titer (from 29.1 mg/L to 94.5 mg/L) and ratio (from 11% to 33%, **Fig. 4b**).

Next, we tried to further increase butyryl-CoA and ethylmalonyl-CoA substrate pools by enzymatically modifying their biosynthetic pathways. Ethylmalonyl-CoA is the third most used extender unit in PKS biosynthesis^48^, and several BGCs that extend with ethylmalonyl-CoA have annotated crotonyl-CoA carboxylase/reductase (CCR) genes in them, presumably for enhancing the CoA substrate supply^49,50^ (**Fig. 4d**). We selected two candidates, *rimJ* CCR in the rimocidin BGC^37^ and *fkbS* CCR in the FK520 BGC^51^. Both CCR genes were integrated into *S. albus* QD28, and the resulting strains showed improved 1,3-HDO titer of 238.9 mg/L (QD76 with *fkbS* integration) and 210.4 mg/L (QD77 with *rimJ* integration) after 7-day cultivation. Remarkably, QD76 completely reversed the product ratio and produced 309.2 mg/L 1,3-diols with 77% being 1,3-HDO, demonstrating robustness of tuning PKS product profiles by adjusting CoA substrate pools. Moreover, increasing glucose concentration in R5 from 1% to 2% led to total production of 517.3 mg/L 1,3-diols, and R5 + 4% glucose produced 765.1 mg/L 1,3-diols in shake flasks (**Fig. 4c**). Finally, we integrated five exogenous ADH genes (*yqhD*, *chnD*, *yahK*, *yjgB* and *ScADH2*)^4,52,53^ into QD76, creating strains QD78-82, and tested whether overexpression of these ADHs can improve 1,3-diol production. Among them, the *yahK*-integrated strain QD80 and the *yjgB*-integrated strain QD81 improved production, with QD80 being the best producer and achieving 1008.5 mg/L 1,3-diol titer after 7-day cultivation in R5 + 2% glucose + 15 mM L-valine, of which 77% was 1,3-HDO (**Fig. 4e**). These results convincingly demonstrate the efficiency and tunability of our PKS-TR platform.

### Bioproduction of medium-chain and branched-chain diols

Branched-chain diol biosynthesis remains a major challenge to this date due to the relatively narrow scope of available pathways for biosynthesis. Amino acids with branched side-chains such as L-valine and L-isoleucine can serve as precursors for branched-chain diol biosynthesis through CoA-independent biosynthesis routes^4^, but the lack of available enzymes to modify amino acid side-chain length and chemical diversity hinders their further application. We aimed to use our PKS-TR platform via AT domain exchange to produce a series of branched-chain diols (**Fig. 5a**). Among our targets, 2-methyl-1,3-butanediol (2-M-1,3-BDO) has been produced in *E. coli* at low titer (12.1 mg/L), whereas 2-methyl-1,3-pentanediol (2-M-1,3-PDO), 2-methyl- 1,3-hexanediol (2-M-1,3-HDO), and three 2-ethyl-1,3-diols were previously biosynthetically inaccessible^4^. For methyl-branched diol production, we replaced the malonyl-CoA–specific AT in the PKS extension module with a methylmalonyl-CoA–specific AT from module 7 of the rimocidin BGC (RimM7 AT) constructing *S. albus* RimM0M1(RimM7 AT)-TR2 (QD66) and *S. albus* RimM0M1(RimM7 AT)-TR2 + FkbS (QD83). We selected 7 days in R5 + 2% glucose + 15 mM L-valine as our standard cultivation method, under which QD66 produced 425.3 mg/L 2- M-1,3-BDO, 56.2 mg/L 2-M-1,3-PDO, and 40.0 mg/L 2-M-1,3-HDO (**Fig. 5b**). Furthermore, no unbranched-chain diol was detected, highlighting a precise PKS engineering strategy (**Supplementary** Figures 18-20). Next, TR screens showed that TR7 was the best TR accepting methyl-branched substrates, resulting in production of 743.6 mg/L 2-methyl-1,3-diols (**Fig. 5b**). We also tested a RimM7 AT homolog in Pim PKS, PimM7 AT, and achieved a similar 2-methyl- 1,3-diol titer of 572.8 mg/L in RimM0M1(PimM7 AT)-TR7 (QD69). In QD69, FkbS CCR overexpression created QD85 strain with increased both 2-M-1,3-HDO titer (from 18.6 mg/L to 79.2 mg/L) and ratio (from 3% to 31%). The relatively low ratio of 2-M-1,3-HDO in the product profile is likely due to accumulation of its carboxylic acid derivative, 2-methyl-3- hydroxyhexanoic acid, as FkbS overexpression also led to 8.6-fold increase of 2-methyl-3- hydroxyhexanoic acid production (**Supplementary** Figure 21).

**Figure 5.**
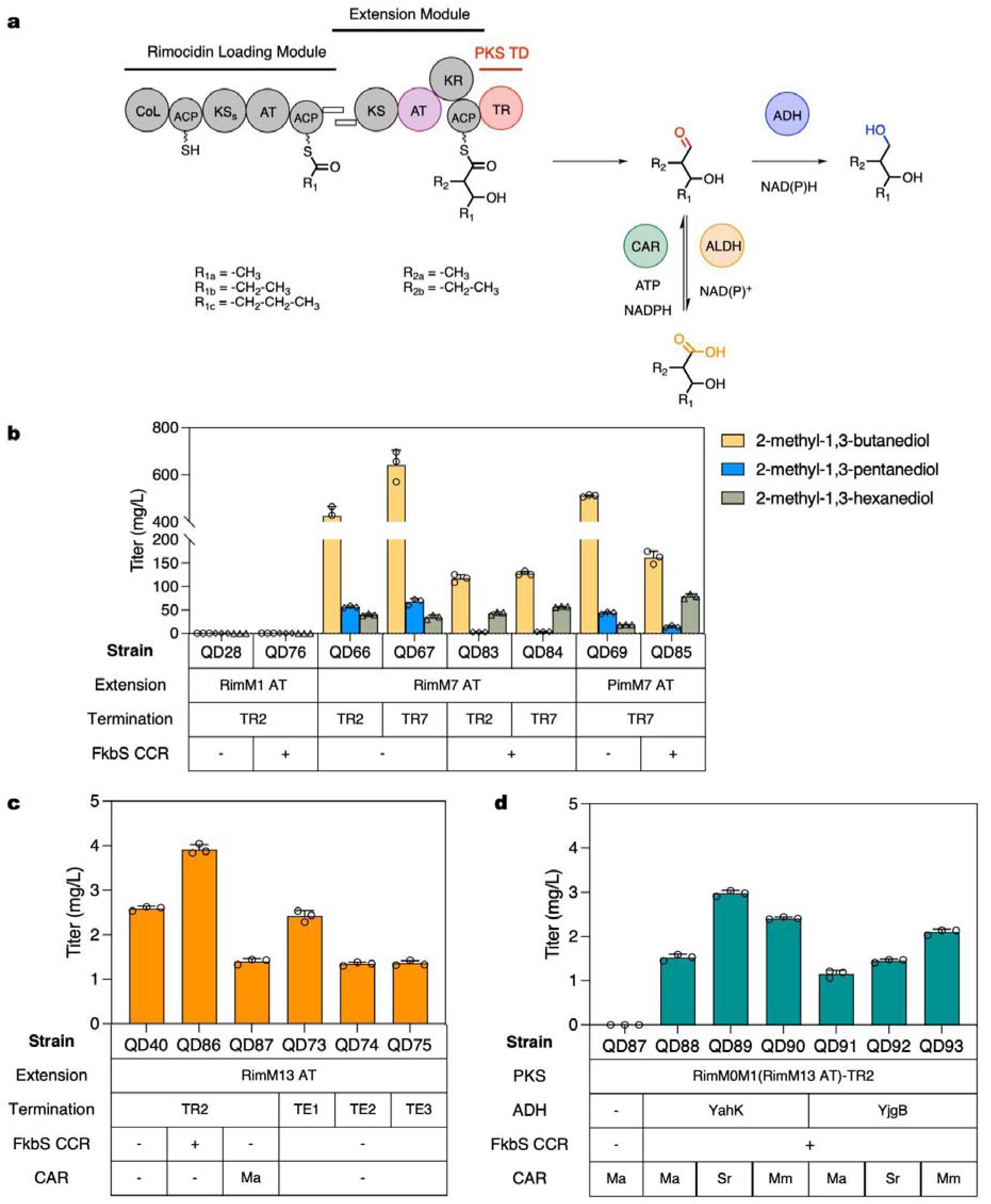
Expanding RimPKS-TR biosynthetic platform for medium- and branched-chain diol and carboxylic acid production. a, Scheme of PKS-TR engineering via AT exchange and post-PKS modification. b, Bioproduction of 2-methyl-1,3-diols in *S. albus* RimM0M1(Rim/PimM7 AT)-TRs, with 2M-1,3-BDO coloured in yellow, 2M-1,3-PDO coloured in blue, and 2M-1,3-HDO coloured in green. c, Bioproduction of 2-ethyl-3-hydroxyhexanoic acid (orange) in engineered *S. albus* RimM0M1(RimM13 AT)-TR2/TEs. TE 1-3: DEBS TE, Pikromycin TE, and Rimocidin TE. d, Bioproduction of insect repellent 2E-1,3-HDO (cyan) in *S. albus* RimM0M1(RimM13 AT)-TR2 + FkbS + ADH + CAR. All data were collected after 7 d cultivation in R5 + 2% glucose + 15 mM L-valine, as mean value and error bars indicate standard deviation (s.d.) of at least three biological replicates.

We next moved on to construct a microbe for 2-E-1,3-HDO production by replacing the malonyl-CoA–specific AT with an ethylmalonyl-CoA–specific AT. RimM13, the last module of Rim PKS, has the same domain composition as RimM1, except RimM13 AT loads ethylmalonyl-CoA based on pathway analysis. This drove us to test a series of RimPKS-TR chimeric designs, including exchanging only RimM1 AT with RimM13 AT (QD40), RimM1 AT-KR-ACP with RimM13 AT-KR-ACP (QD51), or the entire RimM1 with RimM13 (QD42) (**Supplementary** Figure 22). Among these designs, none produced 2-E-1,3-HDO; only *S. albus* RimM0M1(RimM13 AT)-TR2 (QD40) produced 2.6 mg/L 2-ethyl-3-hydroxyhexanoic acid (**Fig. 5c**). To validate whether 2-ethyl-3-hydroxyhexanoic acid was produced via enzymatic condensation of the butyryl starter unit and ethylmalonyl extender unit, we performed [^13^C]labelled L-valine feeding, and observed both half- and fully-[^13^C]labelled 2-ethyl-3- hydroxyhexanoic acid, confirming its biosynthetic route (**Supplementary** Figure 23). We also tested three other TEs as the termination domains; compared to DEBS/Pik/Rim TE, TR2- containing QD40 remained the best 3-hydroxy acid producer. FkbS CCR overexpression in QD40 led to QD86 with an increased 2-ethyl-3-hydroxyhexanoic acid titer at 3.9 mg/L, albeit still with no 2-E-1,3-HDO production.

We reasoned that endogenous ADHs in *S. albus* were not capable of reducing ethyl- branched aldehydes to diols, in agreement with our observation that overexpression of MaCAR carboxylic acid reductase in QD40 did not produce 2-E-1,3-HDO. To enzymatically convert 2- ethyl-3-hydroxyhexanoic acid to 2-E-1,3-HDO, we then tested three CARs (MaCAR/SrCAR/MmCAR) and two ADHs (YahK/YjgB) in QD86, and all the resulting QD88- QD93 microbes produced 2-E-1,3-HDO, with SrCAR + YahK combination (QD89) producing the highest 2-E-1,3-HDO titer (3.0 mg/L, **Fig. 5d and Supplementary** Figure 24). Moreover, LC-MS peaks matching 2-E-1,3-BDO and 2-E-1,3-PDO theoretical [M + Na]^+^ *m/z* were observed (**Supplementary** Figure 25), together with 2-E-1,3-HDO production demonstrating successful engineering of 2-ethyl- extender units into our diol biosynthetic pipelines.

### Bioproduction of amino alcohols via post-PKS transamination

Amino alcohols are also important specialty chemicals in the polymer and pharmaceutical industries, and they can be synthesised using the PKS platform we constructed to synthesise diols. Moreover, most PKS pathways in nature have dedicated post-PKS decoration enzymes with extraordinary diversity and peculiarity, including TR-cognate transaminases, yet their application in PKS engineering has not been reported^20,54^. To explore this possibility (**Fig. 6a**), we synthesised *TA1* and *TA2* genes from Cpk and B24891 BGCs, and integrated them into *S. albus* RimM0M1-TR1 (QD27) or RimM0M1-TR2 (QD28). The four resulting strains (QD94- QD97) successfully produced 4-aminobutan-2-ol, 1-aminopentan-3-ol, and 1-aminohexan-3-ol, confirming the activity of TAs (**Fig. 6b and Supplementary** Figures 26-27). Despite the fact that post-PKS TAs are often TR-cognate in natural PKSs, our choice of TAs was not strictly restricted by the TRs selected for PKS termination, because QD96 with TR2 + TA1 combination was the best amino alcohol producer in R5 + 2% glucose (425.1 mg/L 4-aminobutan-2-ol, 112.1 mg/L 1-aminopentan-3-ol, and 17.1 mg/L 1-aminohexan-3-ol, 554.2 mg/L in total). Moreover, we successfully tuned the amino alcohol product profile by the same CoA substrate regulation strategies: 15 mM L-valine supplemented to QD98 (FkbS overexpression in QD96) cultures led to dramatically increased 1-aminohexan-3-ol titer (from 17.1 mg/L to 364.4 mg/L) and product ratio (from 3% to 70%). The AT exchange was also compatible with PKS-TR-TA-based branched-chain amino alcohol production, as *S. albus* RimM0M1(RimM7 AT)-TR2 + TA1 + FkbS (QD99) produced 21.6 mg/L 4-amino-3-methyl-2-butanol, as well as 1-amino-2-methyl-3- pentanol and 1-amino-2-methyl-3-hexanol (**Fig. 6b and Supplementary** Figures 28-29).

**Figure 6.**
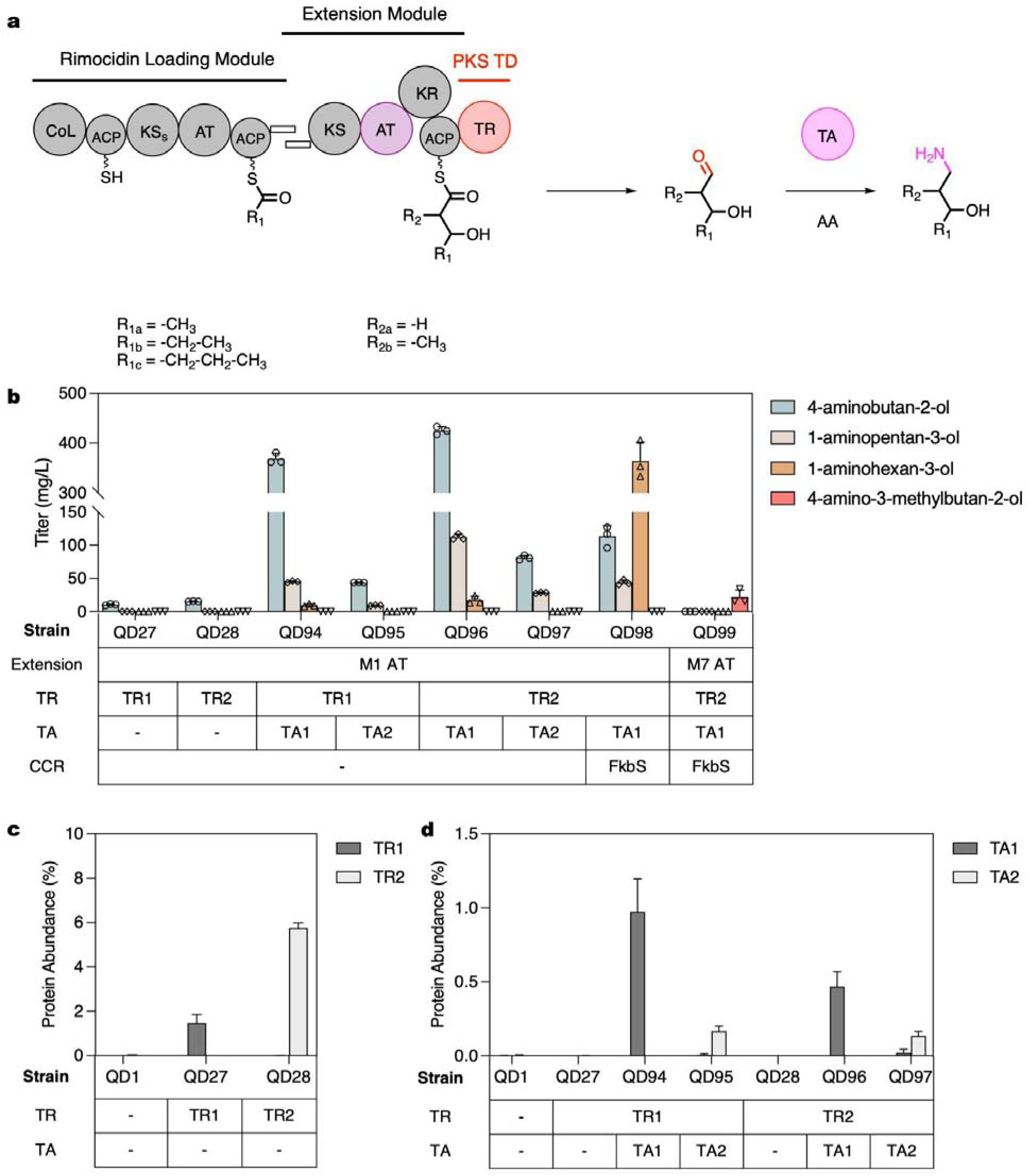
Utilisation of PKS TR-cognate transaminases (TAs) for amino alcohol production. a, Scheme of PKS-TR engineering via PKS TR-cognate transamination. AA: amino acid. b, Bioproduction of 1-amino-3-alcohols in *S. albus* RimM0M1-TRs + TAs after 7 d cultivation in R5 + 2% glucose. 15 mM L-valine was also added in the QD98 and QD99 groups. In QD99, RimM7 AT exchange was applied for methyl-amino-alcohol production. c, Protein abundance of TR1 (black) and TR2 (white). d, Protein abundance of TA1 (black) and TA2 (white). All cultivation data are represented as mean value and error bars indicate standard deviation (s.d.) of at least three biological replicates.

Next, we evaluated the impact of protein abundance on amino alcohol titers. Proteomics analysis confirmed expression of all incorporated genes in *S. albus* and showed higher RimM1- TR2 level than RimM1-TR1 chimaera (**Fig. 6c**), explaining higher product titers when using RimM1-TR2 to produce either 1,3-diols or amino alcohols. Furthermore, the TA1 protein level was 5 times higher than that of TA2 (**Fig. 6d**), also in agreement with the amino alcohol production data. All together, these data demonstrate that optimal protein expression level is key to high product titers by these engineered PKS systems in *Streptomyces* hosts.

Another possible key factor that may play a critical role in PKS-TR-based bioproduction was intracellular NAD(P) levels, as both PKS KRs and TRs are NADPH-dependent, and many ADHs are NADH-dependent. We measured the NAD(P) levels in several of our engineered *S. albus* strains, and concluded that they were similar in all tested strains (**Supplementary** Figure 30), ruling out the possibility of cofactor influence on observed product titers.

## Discussion

Nature designs and produces a wide range of structurally complex molecules with modular type I PKSs, which provide parts to engineer megasynthases for a variety of unnatural molecules that would be difficult or impossible to biosynthesize in any other way. In this study, we developed PKSs with terminal reductases (TRs) to enable biosynthesis of medium- and branched-chain aldehydes. Compared to previously established pathways that can be used to form C-C bonds, such as reverse β-oxidation^55^, PKSs have three major advantages. First, because PKSs liberate carbon dioxide during Claisen condensation, they have a thermodynamic advantage over thiolase-catalysed non-decarboxylative condensation reactions. Taking the 2-ethyl-3- hydroxyhexanoic skeleton as an example, the PKS biosynthesis route has a 3 kcal/mol advantage in Gibbs standard free energy over the thiolase pathway (**Supplementary Table 5**). Second, the PKS biosynthesis route is a platform to many different molecules because of the great structural diversity in the starter choice as well as the extender choice due to AT-gated substrate loading (e.g., desmethyl-, methyl-, ethyl-, allyl-, hydroxy-, methoxy-, and amino-, to name a few^48^). This is particularly advantageous over other routes for adding branches at designated even positions. Furthermore, because of the stepwise catalytic feature and ACP protection of the intermediates, PKS product profiles are more amenable to fine tuning. Rimocidin PKS naturally initiates with acetyl-, propionyl-, and butyryl- starter units, based on which we successfully engineered RimPKS-TRs for bioproduction of a series of unbranched 1,3-diols, including the humectant 1,3- BDO. We also validated the plausibility of the canonical PKS AT exchange engineering strategy when terminating with TRs, presenting a platform suitable for medium-chain methyl diol and ethyl diol biosynthesis, featuring the insect repellent 2-E-1,3-HDO. Finally, natural PKS pathways provide lavish enzymatic toolkits including many novel post-PKS decoration enzymes. In this study we showcased the TR-cognate TAs, incorporation of which led to bioproducts with terminal -NH_2_ in our engineered RimPKS-TR + TA hosts.

Moreover, we achieved tunable product profiles of RimPKS-TRs through substrate CoA pool engineering. PKS pathways are often tightly regulated in nature, as shown by wide presence of PKS pathway specific transcription regulators within the BGCs^56^, and more recent discovery of pyrroloquinoline quinone (PQQ) gene clusters that co-evolved with PKS BGCs and enhanced natural product production^57^. Another common PKS-related regulation approach prioritises regulating CoA substrate biosynthesis, investigation of which benefited identification of key steps in engineering such PKSs. For example, 2(*S*)-ethylmalonyl CoA biosynthesis in *Streptomyces* has been heavily investigated^58^: the major route is a CCR-catalysed reaction to convert crotonyl-CoA to ethylmalonyl-CoA, and a minor pathway is propionyl-CoA carboxylase-catalysed butyryl-CoA carboxylation. Butyryl-CoA is a major product of L-valine catabolism, and multi omics analysis revealed a strong suppression of CCR gene expression upon L-valine supplementation^47^. This observation is indicative of a CCR shortage in our engineered system and may dampen butyryl-based 1,3-HDO production, thus we overexpressed a secondary FkbS or RimJ CCR and reversed the diol production ratio with a single step of engineering, leading to 1,3-HDO product ratio increase from 33% to 77%.

Lastly, terminal TR domains in PKS BGCs were reported in multiple biosynthetic studies, yet their application in PKS engineering was largely uninvestigated; our work presents the first PKS TR engineering report to the best of our knowledge. A common theme of these natural TR-containing pathways are amine production through TR-catalysed reductive cleavage of the thioester bond and subsequent TA-catalysed transamination. TRs were reported to be NADH-dependent, which was unexpected given the fact that all PKSs are secondary metabolic enzymes, and similar PKS KRs are NADPH-dependent. In this study, we concluded that TRs are evolutionarily related to PKS KRs and also NADPH-dependent. Moreover, our observation that TRs do not catalyse aldehyde reduction agrees with the fact that PKS TRs always have cognate TAs in the BGCs reacting on TR-produced aldehydes. The incapability of aldehyde reduction by PKS TRs separates them from NRPS Rs, many of which can catalyse four electron reduction to alcohols^31^. The CpkC TR structure has an open and solvent-exposed substrate binding pocket, which may facilitate the aldehyde escaping the active site after reductive cleavage from the ACP, preventing further reduction to alcohols. Besides modular type-I PKS and NRPS pathways, TRs are also widely present in other secondary metabolism, including putative iterative type I PKSs in *Dictyostelium discoideum*^59,60^, but none of these *Dictyostelium* TRs (DD TRs) cooperated with Rim PKS, and the chimeric Rim PKS-DD TR proteins had no detected function when expressed in *S. albus* or *E. coli* (unpublished data).

Taken together, our study presents the first comprehensive PKS-TR engineering effort targeting diols, amino alcohols and carboxylic acids, which provides valuable toolkits for PKS retrobiosynthesis and lays the foundation for transforming PKS design schemes to produce previously inaccessible alcohols and amines. Our engineering efforts may expand the narrow scope of available biosynthetic pathways for medium- and branched-chain products; further chemical diversification on the C1 aldehyde group can also generate polyketide-based alkenes and alkanes, many of which are interesting biofuels and pharmaceutical intermediates.

## Methods

### Materials

*Streptomyces albus* J1074, *Streptomyces coelicolor* M1152, *E. coli* ET12567/pUZ8002, *E. coli* ET12567/pUB307, *E. coli* BAP1 were collected as previously described^61,62^. *Streptomyces rimosus subsp. rimosus* ATCC 10970 and *Streptomyces natalensis* ATCC 27448 were purchased from ATCC. *Streptomyces tsukubaensis* NRRL 18488 was purchased from NRRL. *E. coli* BL21(DE3), DH5α, and DH10β were purchased from New England Biolabs.

The pSC and p41 *Streptomyces* integration vectors were collected as previously described^63^, pOSV807 (Addgene plasmid 126600) and pOSV809 (Addgene plasmid 126602) *Streptomyces* integration vectors were purchased from Addgene. pHIS (Addgene plasmid 29653) and pMBP (Addgene plasmid 29656) *E. coli* expression vectors were purchased from Addgene as well. pG- KJE8 chaperone plasmid was purchased from Takara Bio.

1,3-butanediol (309433), 2-methyl-butane-1,3-diol (S309524), 2-ethyl-hexane-1,3-diol (E29125), sodium acetate (S2889), sodium malonate (M4795), coenzyme A trilithium salt (C3019), acetyl coenzyme A lithium salt (A2181), malonyl coenzyme A lithium salt (M4263), [^13^C]labelled L-valine, ATP, NAD^+^, NADH, NADP^+^, and NADPH were purchased from Millipore Sigma. 1,3- pentanediol (BD01000039) and 2-ethyl-3-hydroxyhexanoic acid (BD01079153) were purchased from BLDpharm. 1,3-hexanediol (EN300-142438), 2-methylpentane-1,3-diol (EN300-298644), 2-methylhexane-1,3-diol (EN300-7632942), 4-aminobutan-2-ol (EN300-76843), 1-aminopentan- 3-ol (EN300-62209), 1-aminohexan-3-ol (EN300-237616), and 4-amino-3-methylbutan-2-ol (EN300-683176) were purchased from Enamine. 3-hydroxypentanoic acid (A282505) was purchased from AmBeed. 3-hydroxyhexanoic acid (H825320) was purchased from Toronto Research Chemicals.

### Cloning of integration plasmids and *Streptomyces* integration

All integration plasmids were constructed using the same Gibson assembly protocol, with triparental conjugation into *S. albus* J1074, and biparental conjugation into *S. coelicolor* M1152. For *S. albus* J1074 triparental conjugation, using p41_rimB27 (p41 plasmid carrying *Pgapdh(EL)-rimM1-TR1*) as an example, we PCR amplified the VWB integrase gene-containing p41 backbone, *rimM1* piece and *TR1* piece using PrimeSTAR GXL enzyme premix. The plasmid backbone and inserts were ligated through Gibson assembly, and the recombinant plasmid was then transformed into chemically competent *E. coli* DH5α. Plasmid-containing DH5α was inoculated into 5 mL Luria-Bertani (LB) broth overnight at 37 °C with apramycin (50 µg/mL) for plasmid mini-prep. The plasmid sequence was verified by whole-plasmid sequencing. The validated plasmid was subsequently transformed into chemically competent *E. coli* DH10β, which was selected for on LB agar containing apramycin (50 µg/mL). Helper cells ET12567/pUB307 were also plated and selected for on LB agar containing kanamycin (50 μg/mL) and chloramphenicol (25 µg/mL). Integration plasmid-containing DH10β and helper cells ET12567/pUB307 were then inoculated in 10 mL LB broth at 37 °C with proper antibiotics until they reached an O.D. 600 nm value of 0.4. They were centrifuged at room temperature to remove the remaining media, washed with sterile water twice, and finally resuspended in 250 µL - 1 mL LB. 500 µL *S. albus* spores that were kept at -80 °C were heat-activated at 50 °C for 10 min and mixed with 250 µL plasmid-containing DH10β and 250 µL ET12567/pUB307. They were then concentrated to 100 µL final volume, plated on mannitol soy agar (2% mannitol, 2% soy flour, and 10 mM MgCl_2_), and incubated at 30 °C for 16 h. 1 mL nalidixic acid (50 µg/mL) and apramycin (50 µg/mL) were overlaid the next day to eliminate those *S. albus* that failed to integrate the recombinant plasmid.

For *S. coelicolor* M1152 biparental conjugation, using pSC_rimA0 (pSC plasmid carrying *PrpsL(RO)-rimM0*) as an example, the validated pSC_rimA0 plasmid obtained from *E. coli* DH5α was transformed into *E. coli* ET12567/pUZ8002 through electroporation. *E. coli* ET12567/pUZ8002, serving as the conjugal donor, was then grown and selected for in 10 mL LB broth containing spectinomycin (100 µg/mL), kanamycin (50 μg/mL) and chloramphenicol (25 µg/mL) at 37 °C until it reached an O.D. 600 nm value of 0.4. The following conjugation procedure with heat-activated *S. coelicolor* M1152 spores and antibiotics overlay (1 mL 50 µg/mL nalidixic acid and 400 µg/mL spectinomycin) was identical to the triparental conjugation experiments.

Conjugated *Streptomyces* colonies were inoculated into 3 mL Tryptic Soy Broth (TSB) medium with 2-3 plating beads in a 24-well block at 30 °C for 2 d, and 100uL culture aliquots were taken and boiled at 98 °C for 20 min to verify successful integration via genome PCR amplification.

### *Streptomyces* collection and cultivation

100 uL of *Streptomyces* culture with successful integration was spread evenly on a mannitol soy agar containing nalidixic acid (50 µg/mL) and grown at 30 °C for 7 d to allow sporulation. Spores were then harvested from the plate with 5 mL of 2×YT medium and filtered by a sterilised cotton syringe. The spore mixture was subsequently mixed with 2 - 3 ml of 60% glycerol stock and stored at -80 °C. For *Streptomyces* cultivation, spores were 1:100 inoculated in 3 mL TSB medium with 2-3 plating beads and cultured at 30 °C for 2 d. 1 mL of *Streptomyces* seed culture was then added to 30 mL broth in a 250 mL baffled shake flask for cultivation at 30 °C and 200 r.p.m.. TSB medium: 17 g tryptone, 3 g soytone, 5 g NaCl, 2.5 g K_2_HPO_4_, 2.5 g glucose, in 1 L dH_2_O, pH = 7.3. Standard R5 medium: 103 g sucrose, 0.25 g K_2_SO_4_, 10.12 g MgCl_2_·6H_2_O, 10 g glucose, 0.1 g Difco casamino acids, 5 g Difco yeast extract, 5.73 g TES buffer, and 2 ml trace element solution were added in dH_2_O (1 L final). 1 mL trace element solution was prepared by 40 mg ZnCl_2_, 200 mg FeCl_3_·6H_2_O, 10 mg CuCl_2_·2H_2_O, 10mg MnCl_2_·4H_2_O, 10mg Na_2_B_4_O_7_·10H_2_O, and 10 mg (NH_4_)_6_Mo_7_O_24_·4H_2_O. Before use, 100 mL R5 was mixed with 1 mL KH_2_PO_4_ (0.5%), 0.4 mL CaCl_2_·2H_2_O (5M), 1.5 mL L-proline (20%), and 0.7 mL NaOH (1N). ISP2: 4 g yeast extract, 10 g malt extract, 4 g dextrose, in 1 L dH_2_O. ISP4: 10 g soluble starch, 1 g MgSO_4_·7H_2_O, 1 g NaCl, 2 g (NH_4_)_2_SO_4_, 2 g CaCO_3_, 1 mL trace salts solution (0.1 g FeSO_4_·7H_2_O, 0.1 g MnCl_2_·4H_2_O, 0.1 g ZnSO_4_·7H_2_O in 100 mL dH_2_O). M042 was prepared as previously reported^16^. 1 mL of *Streptomyces* culture was collected at different time points (maximum 10 d) for LC-MS analysis.

### LC-MS sample preparation and analysis

1 mL of *Strept*omyces culture was centrifuged at 4,000*g* for 5 min to collect the soluble fraction, which was subsequently mixed with the same volume of LC-MS grade MeOH, vortexed for 10 s, and centrifuged in Millipore Amicon Ultra-0.5 centrifugal filters (3 kDa) at 14,000*g* for 15 min. The follow-through solution was diluted in 50% LC-MS grade MeOH by 25 folds if necessary, and analysed by LC-MS using the Agilent LC/MSD iQ single quadrupole mass spectrometer. Accurate mass measurements were performed via an Agilent Technologies 6520 Q-TOF LC/MS system.

For 1,3-diol or 3-hydroxy acid detection and quantification, 5 µL samples were injected onto Phenomenex Kinetex XB-C18 (2.6 µm, 100 × 3 mm, 100 Å) LC Column, and analysed with the following HPLC protocol: buffer A: water with 0.1% v/v formic acid; buffer B: MeOH with 0.1% v/v formic acid; flow rate: 0.42 mL/min; 20 – 72.1% buffer B gradient for 4.5 min, 72.1 – 95% buffer B gradient for 1.3 min, 95% buffer B for 3 min, 95 – 20% buffer B for 0.2 min, 20% buffer B for 2.2 min; mass detection rage: m/z = 50 – 400.

For amino alcohol detection and quantification, 5 µL samples were injected onto Agilent Technologies ZORBAX Eclipse Plus C18 (3.5 μm, 4.6 mm × 150 mm) LC column, and analysed with the following HPLC protocol: buffer A: water with 0.1% v/v formic acid; buffer B: acetonitrile with 0.1% v/v formic acid; flow rate: 0.40 mL/min, 2% buffer B for 0.5 min, 2 – 13% buffer B gradient for 4.5 min, 13 – 80% buffer B gradient for 0.1 min; flow rate changed to 1.0 mL/min, 80% buffer B for 2 min, 80 – 2% buffer B for 3.1 min, 2% buffer B for 1.1 min; flow rate changed to 0.4 mL/min, 2% buffer B for 1.1 min; mass detection rage: m/z = 70 – 300.

### 1,3-diol/3-hydroxy acid/[13C]labelled L-valine feeding

For 1,3-diol/3-hydroxy acid feeding experiments, 50 mg/L 1,3-diol or 3-hydroxy acid was added to 30 mL R5 + 1% glucose, and 1 mL of *Streptomyces* seed culture was then inoculated into the broth in a baffled shake flask for cultivation at 30 °C and 200 r.p.m.. Culture samples were collected every 24 h for LC-MS quantification as previously described. For [^13^C]labelled L-valine feeding, 5 mM or 10 mM [^13^C]labelled L-valine was added to 30 mL R5 + 1% glucose. After *Streptomyces* seed culture inoculation, LC-MS samples were collected after 3 d cultivation.

### Cloning of expression plasmids

*RimM0*, *pimS0*, *TADH2*, *ACP9*, *TR1*, *TR2*, *TR7*, and *TR9* genes were cloned into the pHIS vector via Ligation-Independent Cloning (LIC). *ACP1* was cloned into the pMBP vector. The resulting plasmids (pHIS_rimA, pHIS_pimS0, pHIS_TADH2, pMBP_ACP1, pHIS_ACP9, pHIS_TR1, pHIS_TR2, pHIS_TR7, pHIS_TR9) were transformed into DH5α and miniprepped using QIAprep Spin Miniprep Kit. All the plasmid sequences were validated by whole-plasmid sequencing from Primordium Labs.

### Protein expression and purification

For protein expression, pHIS or pMBP expression plasmid was co-transformed with pG-JKE8 plasmid into *E. coli* BL21(DE3). For pHIS_rimA or pHIS_pimS0, *E. coli* BAP1 was used for *holo*-protein expression. A single colony was inoculated in 10 mL LB broth with 50 μg/mL kanamycin and 25 μg/mL chloramphenicol, and grown overnight at 37 °C, 200 r.p.m.. The overnight culture was inoculated into 500 mL Terrific Broth + 4% v/v glycerol with 50 μg/mL kanamycin and 25 μg/mL chloramphenicol in a 2 L baffled flask, and incubated at 37 °C, 200 r.p.m. until O.D. 600 nm reached 0.6. The flask was then transferred to 20 °C, incubated for 1 h, 200 r.p.m., and induced with 1 mg/mL L-arabinose, 10 ng/mL tetracycline, and 0.2 mM IPTG. After 20 h induction, the cells were centrifuged at 4000*g* for 30 min and the cell pellet was harvested and stored at -20 °C.

For protein purification, the cell pellet was resuspended in 50 mL lysis buffer (10% v/v glycerol, 300 mM NaCl, 20 mM imidazole pH 7.5, 25 mM HEPES buffer pH 7.5, 0.1 mg/mL lysozyme, 0.05 mg/mL DNase, and 1 mM MgCl_2_) and vortexed for 30 min. To completely lyse the cells, sonication (amplitude 40%, 30 s on, 1 min off, repeat for 3 cycles) was applied at 4 °C. The cells were centrifuged at 20000*g* for 30 min, and the lysate supernatant was collected in a 50 mL falcon tube after 0.22 μm filtration. The supernatant was subsequently subjected to Ni-NTA affinity chromatography in 4 mL/min flow rate, and washed with 10 column volumes of Ni-NTA buffer (10% v/v glycerol, 300 mM NaCl, 20 mM imidazole pH 7.5, 25 mM HEPES buffer pH 7.5). The His-tagged protein was eluted with an imidazole gradient from 20 mM to 500 mM in 15 min. Fractions containing the target His-tagged protein were analysed by SDS-PAGE, pooled and incubated with 5 mM ATP pH 7.0 for 1 h to completely dissociate attached chaperones. After ATP incubation, the protein solution was concentrated using Amicon Ultra-15 centrifugal filters to a final volume of 4 – 5 mL. To achieve higher purity and better understanding of the protein oligomeric distribution, the protein solution was further subjected to size-exclusion chromatography (SEC) with a GE Hiload 16/60 Superdex 200 prep grade column equilibrated with 10% v/v glycerol, 50 mM NaCl, 25 mM HEPES pH 7.5 in 1 mL/min flow rate. SEC fractions were assessed by SDS-PAGE, and ideal fractions were concentrated to 5 – 40 mg/mL, aliquoted and flash cooled in liquid N_2_, and stored at -80 °C. All proteins have > 95% homogeneity, except for PimS0 with ∼80% purity.

### Crystallisation, X-Ray data collection and structure determination of CpkC TR

The CpkC TR sample was concentrated at 10 mg/ml. The cofactor NADP^+^ was added prior to crystallisation trials to a final concentration of 5 mM. The CpkC TR in complex with NADP^+^ was screened against the crystallisation set of solutions: Berkeley Screen^64^, MCSG-1 (Anatrace), ShotGun (Molecular Dimensions), PEG/Ion, Index, Crystal Screen, and PEGRx (Hampton Research). Crystals of CpkC TR were found in Berkeley Screen condition B3 composed of 0.4 M sodium chloride, 0.1 M BIS-Tris pH 6.5 and 30 % PEG 3,350. The crystal of CpkC TR was placed in a reservoir solution containing 20% (v/v) glycerol, then flash-cooled in liquid nitrogen. The X-ray data set for CpkC TR was collected at the Berkeley Center for Structural Biology beamline 5.0.1 at the Advanced Light Source at Lawrence Berkeley National Laboratory. The diffraction data were processed using the program Xia2^65^. The crystal structure of CpkC TR – NADP^+^ was solved by molecular replacement with the program PHASER^66^ using initial coordinates of the CpkC TR model generated by ALPHAFOLD^67^. The atomic positions obtained from the molecular replacement were used to initiate refinement within the Phenix suite^68^. Structure refinement was performed using the phenix.refine program. Manual rebuilding was done using COOT^69^. Root-mean-square deviations from ideal geometries for bond lengths, bond angles and dihedral angles were calculated with Phenix.refine^70^. The stereochemical quality of the final model of CpkC TR was assessed by the program MOLPROBITY^71^. Summary of crystal parameters, data collection, and refinement statistics can be found in Supplementary Table 4.

### Proteomics analysis

*Streptomyces* cultures were collected after 3 d. Cells were harvested and stored at -80 °C until further processing. Protein was extracted from cell pellets and tryptic peptides were prepared by following established proteomic sample preparation protocol^72^. Briefly, cell pellets were resuspended in Qiagen P2 Lysis Buffer (Qiagen, Germany) to promote cell lysis. Proteins were precipitated with addition of 1 mM NaCl and 4x vol acetone, followed by two additional washes with 80% acetone in water. The recovered protein pellet was homogenised by pipetting mixing with 100 mM ammonium bicarbonate in 20% methanol. Protein concentration was determined by the DC protein assay (BioRad, USA). Protein reduction was accomplished using 5 mM tris 2- (carboxyethyl)phosphine (TCEP) for 30 min at room temperature, and alkylation was performed with 10 mM iodoacetamide (IAM; final concentration) for 30 min at room temperature in the dark. Overnight digestion with trypsin was accomplished with a 1:50 trypsin:total protein ratio. The resulting peptide samples were analysed on an Agilent 1290 UHPLC system coupled to a Thermo Scientific Orbitrap Exploris 480 mass spectrometer for discovery proteomics^73^. Briefly, peptide samples were loaded onto an Ascentis® ES-C18 Column (Sigma–Aldrich, USA) and were eluted from the column by using a 10 minute gradient from 98% solvent A (0.1 % FA in H_2_O) and 2% solvent B (0.1% FA in ACN) to 65% solvent A and 35% solvent B. Eluting peptides were introduced to the mass spectrometer operating in positive-ion mode and were measured in data-independent acquisition (DIA) mode with a duty cycle of 3 survey scans from m/z 380 to m/z 985 and 45 MS2 scans with precursor isolation width of 13.5 m/z to cover the mass range. DIA raw data files were analysed by an integrated software suite DIA-NN^74^. The databases used in the DIA-NN search (library-free mode) are *S. albus* and *S. coelicolor* latest Uniprot proteome FASTA sequences plus the protein sequences of the heterologous proteins and common proteomic contaminants. DIA-NN determines mass tolerances automatically based on first pass analysis of the samples with automated determination of optimal mass accuracies. The retention time extraction window was determined individually for all MS runs analysed via the automated optimization procedure implemented in DIA-NN. Protein inference was enabled, and the quantification strategy was set to Robust LC = High Accuracy. Output main DIA-NN reports were filtered with a global FDR = 0.01 on both the precursor level and protein group level. The Top3 method, which is the average MS signal response of the three most intense tryptic peptides of each identified protein, was used to plot the quantity of the targeted proteins in the samples^75,76^. The generated mass spectrometry proteomics data have been deposited to the ProteomeXchange Consortium via the PRIDE partner repository with the dataset identifier PXD046595^77^. DIA-NN is freely available for download from https://github.com/vdemichev/DiaNN.

### Enzymatic assays: PimS0 substrate loading

PimS0 was added in 100 μL reaction mix (10% v/v glycerol, 50 mM NaCl, 25 mM HEPES pH 7.5, 5 mM ATP, 1 mM acetate + CoA/malonate + CoA/acetyl-CoA/malonyl-CoA) in 7.5 mg/mL final concentration to initiate the substrate loading reaction. At 1 h, 4 h, and 24 h, 20 uL reaction solution was quenched with 1% formic acid, flash cooled in liquid N_2_, and stored at -80 °C for subsequent treatment. PimS0 with no tested substrate added in the reaction mix was taken as a negative control. Proteins in the reaction samples were precipitated by addition of 1 mM NaCl and 4 x vol acetone, followed by two additional washes with 80% acetone in water. Proteins were resuspended with 100 mM ammonium bicarbonate, reduced in 5 mM tris 2- (carboxyethyl)phosphine (TCEP) and alkylated in 10 mM iodoacetamide (IAM) before subjecting to trypsin digestion. The resulting peptides were analyzed using an Agilent 1290 Infinity liquid chromatography system coupled to an Agilent 6460 QQQ mass spectrometer (Agilent Technologies, Santa Clara, CA). Peptides (∼10 μg) were separated on an Ascentis Express Peptide ES-C18 column (2.7 μm particle size, 160 Å pore size, 50 x 2.1mm) fitted with a guard column (5 mm x 2.1 mm, Sigma Aldrich). The column was heated to 60°C. The mobile phase consisted of 0.1% formic acid in H2O (A) and 0.1% formic acid in acetonitrile (B). Peptides were eluted from the column by using a 3.5 minute linear gradient from 95% solvent A and 2% solvent B to 60% solvent A and 40% solvent B. Peptides were ionized using an Agilent Jet Stream ESI source operating in positive-ion mode with the following source parameters: Gas Temperature = 250°C, Gas Flow = 13 L/min, Nebulizer Pressure = 35 psi, Sheath Gas Temperature = 250°C, Sheath Gas Flow = 11 L/min, and Capillary Voltage = 3,500 V. Dwell times were set to 18ms. Data was acquired using Agilent MassHunter Data Acquisition (Version B.08.02). The MRM method for quantifying phosphopantetheine bearing peptides was built using the Skyline (version 21.2). LC-MS raw data were imported and analysed in Skyline. The MRM transitions and their integrated peak areas are available on the LC-MS data sharing platform Panorama Public^78^.

### Enzymatic assays: TR-catalysed reduction

Purified Sfp was purchased from NEB. To perform Sfp treatment for the formation of *holo*- ACPs, purified *apo*-MBP-ACP1 and *apo*-ACP9 were added separately in 500 uL reaction mixture (10% v/v glycerol, 50 mM NaCl, 50 mM HEPES pH 7.5, 2.5 mM octanoyl-CoA, 4 μM Sfp, and 20mM MgCl_2_) and incubated at 30 °C for 3 h. The reaction mixture was then placed in the dialysis buffer (10% v/v glycerol, 50 mM NaCl, 50 mM HEPES pH 7.5) at 4 °C for 3 h to remove any remaining free octanoyl-CoA. To study the substrate scope and co-factor preference of TR-catalysed reduction, TR1 and TR9 storing in buffer (10% v/v glycerol, 50 mM NaCl, 25 mM HEPES pH 7.5) were tested respectively with 20 μM final concentration in a 200 μL reaction mixture containing any of the three substrates (0.5 mM octanoyl-ACP, 1 mM octanoyl-CoA, or 1 mM octanal), along with either of the two cofactors (200 μM NADH and 200 μM NADPH). These experiments were conducted in a Corning 96-well clear bottom black microplate, and 340 nm absorbance was monitored for the detection of NAD(P)H elimination for a maximum of 16 h. All assays were repeated in duplicate.

### Enzymatic assays: 1,3-diol oxidation

300 uL reaction mixture (10% v/v glycerol, 50 mM NaCl, 25 mM HEPES pH 7.5, 0.5 mM NAD(P)^+^, 20 mM alcohol substrate) was added in a Corning 96-well clear bottom black microplate. To initiate the reaction, TADH2 or TR1/2/7/9 enzyme was added at 0.4 μM final concentration, and 340 nm absorbance was monitored for the detection of NAD(P)H accumulation for 2 h. All assays were repeated in triplicate.

### Intracellular NADP and NAD quantification

The measurements were conducted by following the manufacture of NADP/NADPH Quantification Kit (MAK038) and NAD/NADH Quantification Kit (MAK037) from Sigma Aldrich. 10 O.D. (O.D.600) of *S. albus* cell tissue were harvested after culturing in R5 medium for 3 d at 30 °C, 200 r.p.m.. Cells were collected by centrifuge at 15,000*g*, 4 °C, 5 mins, the supernatants were discarded, and the cells were washed with pre-chilled PBS at 15,000*g*, 4 °C, 5 mins. Discarding the PBS and adding 500 µL of NADP/NADPH Extraction buffer or NAD/NADH Extraction buffer for NADP/NADPH measurement and NAD/NADH extraction respectively. 0.5 mm Glass beads (BioSpec Products, Cat. No. 11079105) were added for bead beating with the following process, 3,800 HZ for 30 s, on ice for 1 min, and 3 cycles total. The extracted samples were sitting on ice for 10 min, then centrifuge at 15,000 *g*, 4 °C, 10 min, the supernatant were filtering through a 10 kDa cut-off spin filter (Merck Millipore, UFC501096). The filtered samples were used for measurement, and the samples measurement is following the procedure in the Kits. All assays were repeated in at least duplicate.

### Data Availability

Coordinates and associated structure factors have been deposited with the PDB under accession code 8V1X.

## Acknowledgements

This work was part of the DOE Joint BioEnergy Institute (https://www.jbei.org) supported by the U.S. Department of Energy, Office of Science, Office of Biological and Environmental Research through Contract [DE-AC02-05CH11231] between Lawrence Berkeley National Laboratory and the U.S. Department of Energy, and the DOE Distinguished Scientist Fellow Program to J.D.K.. We thank Dr. Leonard Katz, Dr. J.L. (Clem) Fortman, Dr. Ning Qin, Dr. Jing Huang, and Dr. Matthias Schmidt for helpful discussions.

## Author Contributions

Q.D. and J.D.K. conceived the project. Q.D., Y.C., N.L., J.H.P., X.Z., Y.C., S.C., Y.R., J.G., A.R., T.W.H.B., and E.E.K.B. performed experiments. Q.D. and J.D.K. evaluated the data, and all authors contributed to preparing the manuscript.

## Author Information

J.D.K. has financial interests in Ansa Biotechnologies, Apertor Pharma, Berkeley Yeast, BioMia, Cyklos Materials, Demetrix, Lygos, Napigen, ResVita Bio, and Zero Acre Farms. The other authors declare no competing interests. Correspondence and requests for materials should be addressed to J.D.K.

