## Supplemental Figures 1-30 and Supplemental Tables 1-5 for "A polyketide-based biosynthetic platform for diols, amino alcohols and hydroxyacids"

**Table of Contents**

Supplementary Figure 1. Phylogenetic analysis of PKS TRs.	3

Supplementary Figure 2. Natural substrates of PKS TRs..	4

Supplementary Figure 3. SDS-PAGE analysis of purified TR1 (a), TR2 (b), TR7 (c), TR9 (d), maltose binding protein (MBP)-ACP1 (e), ACP9 (f), and TADH2 (g).	5

Supplementary Figure 4. TR substrate scope analysis and initial reaction rate measurement.	6

Supplementary Figure 5. Multiple sequence alignment of ACP-TRs.	7

Supplementary Figure 6. Terminal thioreductase structures in carrier protein-containing multi-domain complexes..	8

Supplementary Figure 7. Scheme of the rimocidin biosynthetic gene cluster..	9

Supplementary Figure 8. Purification of holo-PimS0 (pPant-attached PimS0) loading module in *E. coli* BAP1 cells.	10

Supplementary Figure 9. Substrate scope analysis of PimS0 loading module.	11

Supplementary Figure 10. Promoter screening results of *rimM0* in *S. albus* RimM0M1-TR2 (QD28-QD30)..	12

Supplementary Figure 11. 1,3-Diol volatility and consumption test..	13

Supplementary Figure 12. MS spectra of 1,3-diols produced in *S. albus* RimM0M1-TR1 (QD27). .	14

Supplementary Figure 13. *S. albus* RimM0M1-DEBS TE (QD18) production test.	15

Supplementary Figure 14. TR substrate scope analysis with 1,3-butanediol.	16

Supplementary Figure 15. TR substrate scope analysis with 1,3-pentanediol.	17

Supplementary Figure 16. Medium screening for *S. albus* RimM0M1-TR1 (QD27).	18

Supplementary Figure 17. [^13^C]labelled L-valine feeding experiment on *S. albus* RimM0M1-TR2 (QD28). .	19

Supplementary Figure 18. LC-MS detection of 2-methyl-1,3-diols produced in *S. albus* RimM0M1(RimM7 AT)-TR2 + FkbS (QD83)..	20

Supplementary Figure 19. MS spectra of 2-methyl-1,3-diols produced in *S. albus* RimM0M1(RimM7 AT)-TR2 + FkbS (QD83).	21

Supplementary Figure 20. Comparative analysis of 1,3-diols produced in *S. albus*.	22

Supplementary Figure 21. Detection of 2-methyl-3-hydroxyhexanoic acid produced in *S. albus*......	23

Supplementary Figure 22. Detection of 2-ethyl-3-hydroxyhexanoic acid produced in *S. albus*.	24

Supplementary Figure 23. [^13^C]labelled L-valine feeding experiment on *S. albus* RimM0M1(RimM13 AT)-TR2 (QD40). .	25

Supplementary Figure 24. Detection of 2-ethyl-1,3-hexanediol produced in *S. albus*.	26

Supplementary Figure 25. Detection of 2-E-1,3-BDO and 2-E-1,3-PDOproduced in *S. albus*.	27

Supplementary Figure 26. LC-MS detection of amino alcohols produced in *S. albus* RimM0M1-TR2 + TA1 (QD96)..	28

Supplementary Figure 27. MS spectra of amino alcohols produced in *S. albus* RimM0M1-TR2 + TA1 (QD96)..	29

Supplementary Figure 28. LC-MS detection of methyl amino alcohols produced in *S. albus* RimM0M1(RimM7 AT)-TR2 + TA1 + FkbS (QD99).	30

Supplementary Figure 29. MS spectra of amino alcohols produced in *S. albus* RimM0M1(RimM7 AT)-TR2 + TA1 + FkbS (QD99)..	31

Supplementary Figure 30. Intracellular NAD(P) level measurement in *S. albus*.	32

Supplementary Table 1. Plasmids and strains used in this study	33

Supplementary Table 2. Primers used in this study	39

Supplementary Table 3. Genes used in this study	43

Supplementary Table 4. Statistics for data collection and refinement of CpkC TR	53

Supplementary Table 5. Thermodynamic analysis of different synthesis pathways of 2-ethyl-1,3-hexanediol and 2-ethyl-3-hydroxyhexanoic acid	54

Supplementary References	55

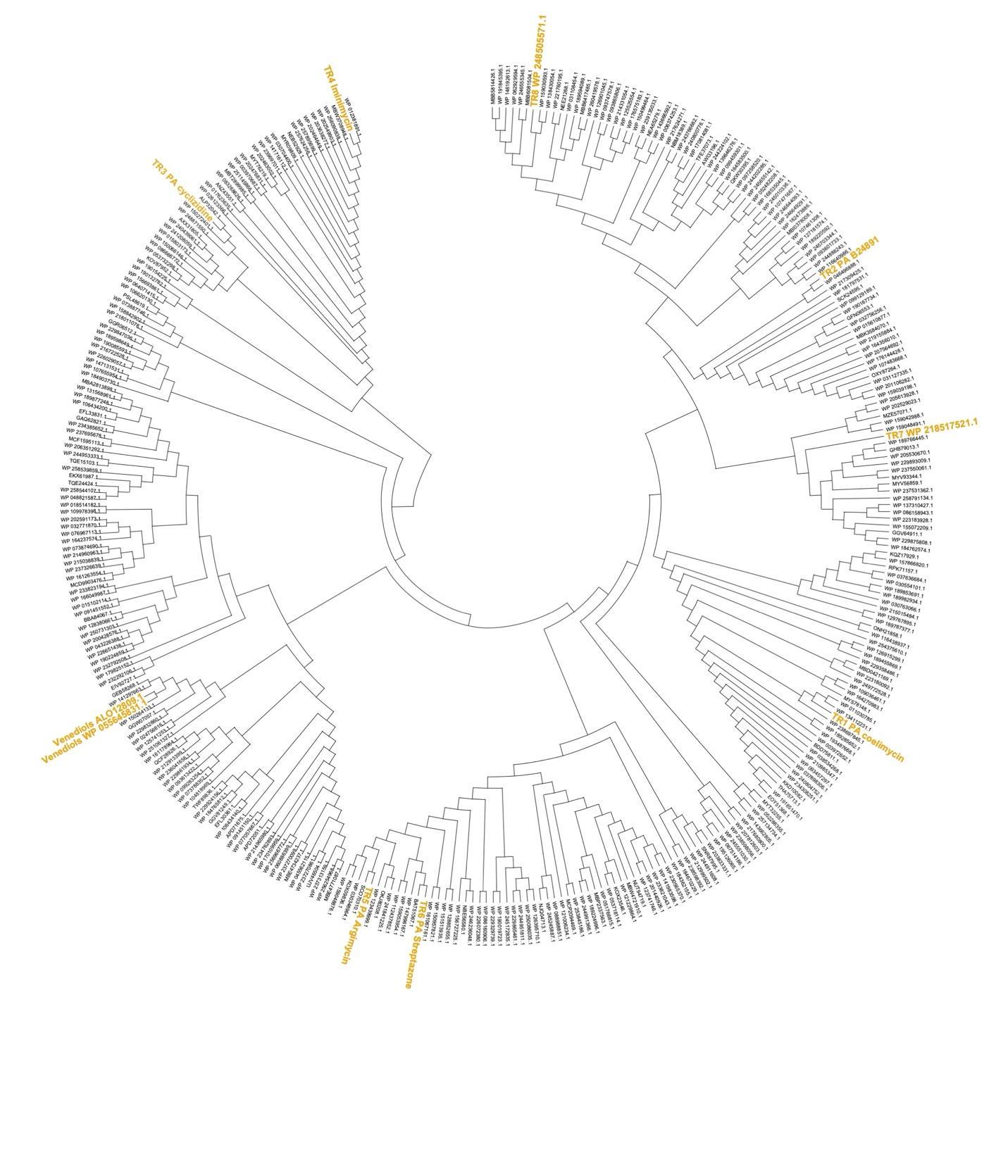

### **Supplementary Figure 1. Phylogenetic analysis of PKS TRs.** TR1 is from coelimycin pathway^1^, TR2 is from B24891 pathway, TR3 is from cyclizidine pathway^2^, TR4 is from iminimycin pathway^3^, TR5 is from argimycin pathway^4^, TR6 is from streptazone pathway^5^, TR7 and TR8 are from unknown pathways, and TR9 is from venediol pathway^6^.

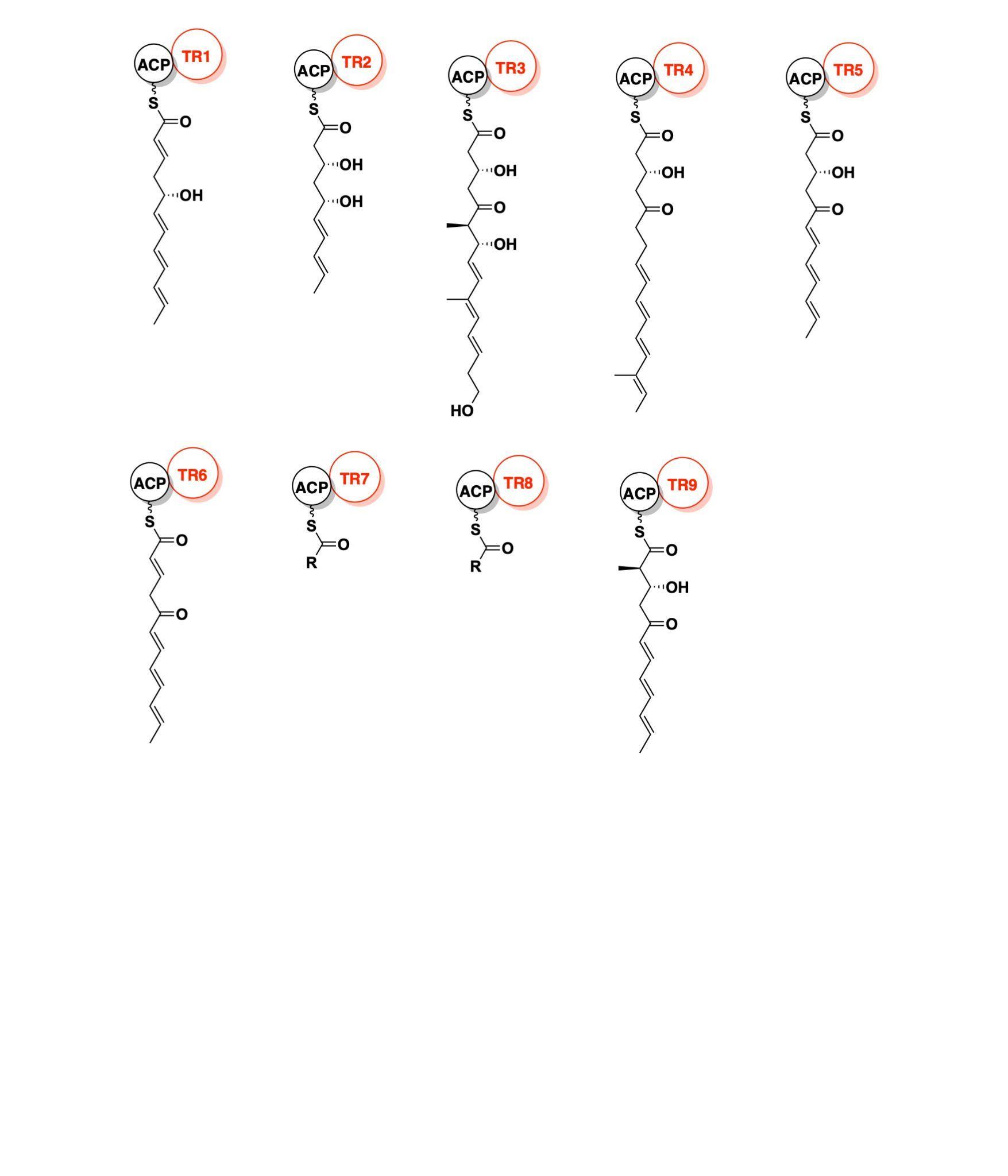

### **Supplementary Figure 2. Natural substrates of PKS TRs.** TR7- and TR8-containing pathway products are not known. All natural products are in the range of medium-chain or long-chain.

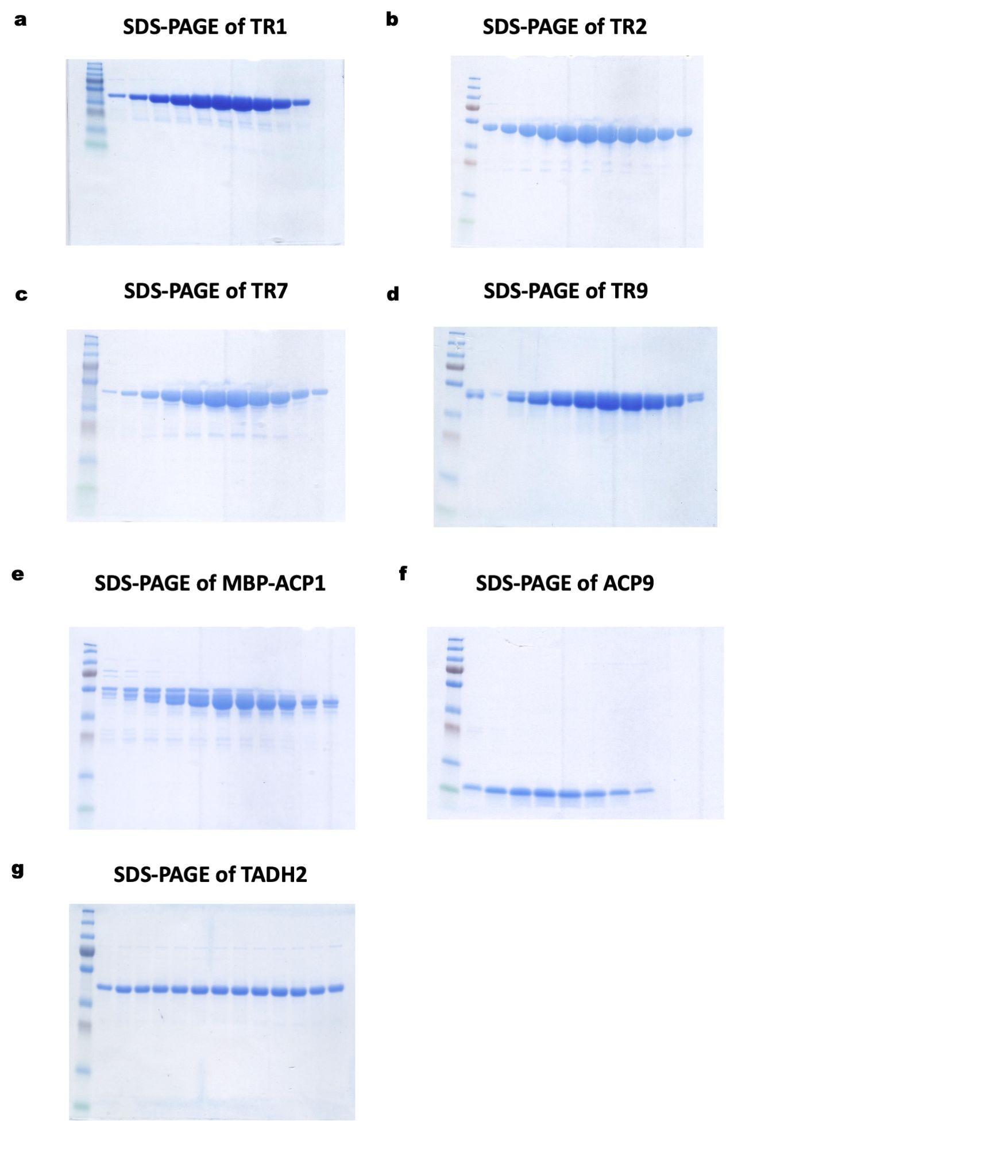

### **Supplementary Figure 3. SDS-PAGE analysis of purified TR1 (a), TR2 (b), TR7 (c), TR9 (d), maltose binding protein (MBP)-ACP1 (e), ACP9 (f), and TADH2 (g)^7^.** The protein ladders are PageRule Plus prestained protein ladders, 10 to 250 kDa.

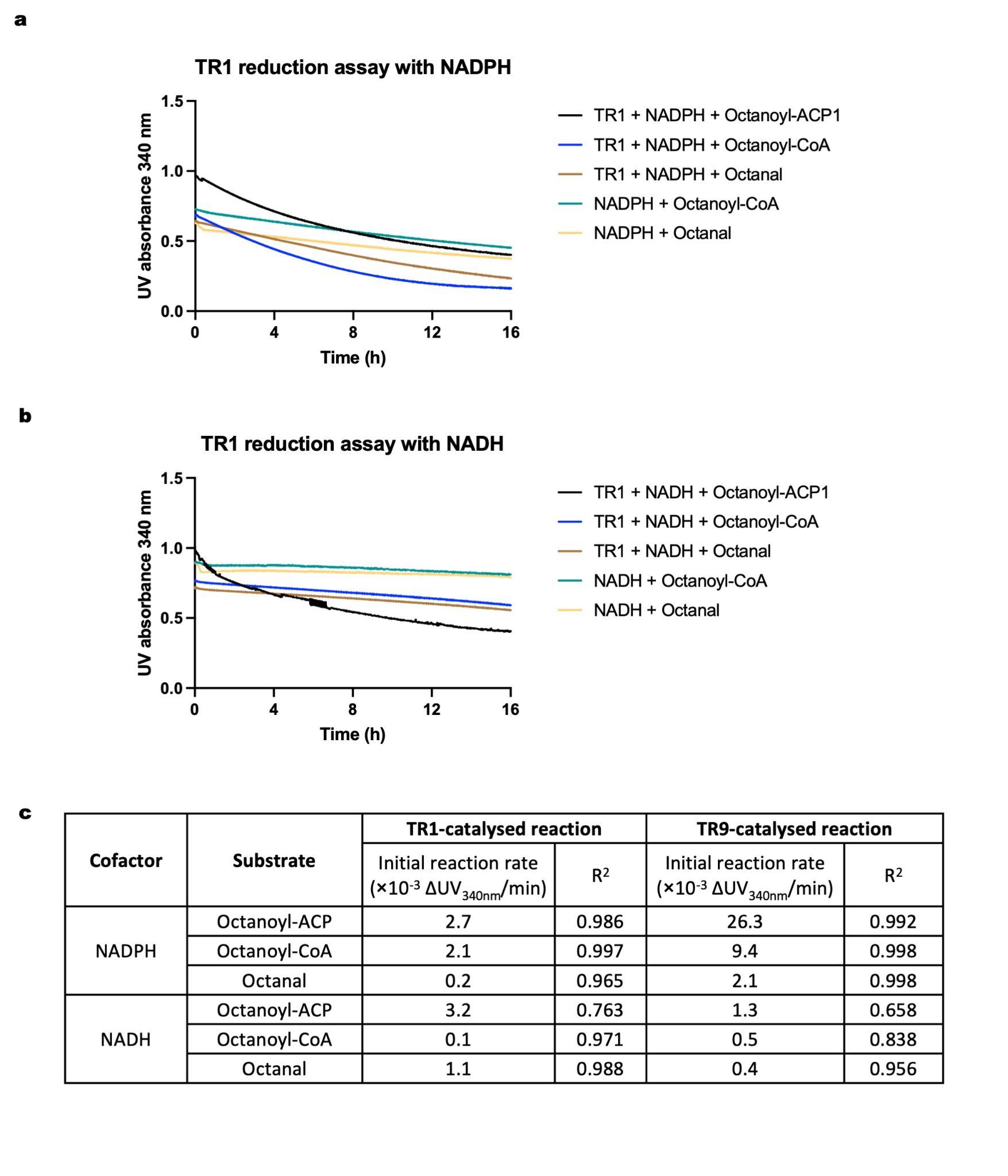

### **Supplementary Figure 4. TR substrate scope analysis and initial reaction rate measurement.** A-B, TR1 substrate scope analysis, with NADPH (a) or NADH (b) as the cofactor. Black: TR1 + NAD(P)H + octanoyl-MBP-ACP1; Blue: TR1 + NAD(P)H + octanoyl-CoA; Brown: TR1 + NAD(P)H + octanal; Green: NAD(P)H + octanoyl-CoA; Yellow: NAD(P)H + octanal. Each experiment was repeated in duplicate. C, Initial reaction rates of TR1- and TR9-catalysed reactions. R^2^: the coefficient of determination in linear regression. Octanoyl-ACP is octanoyl-MBP-ACP1 for the TR1 reaction and octanoyl-ACP9 for the TR9 reaction.

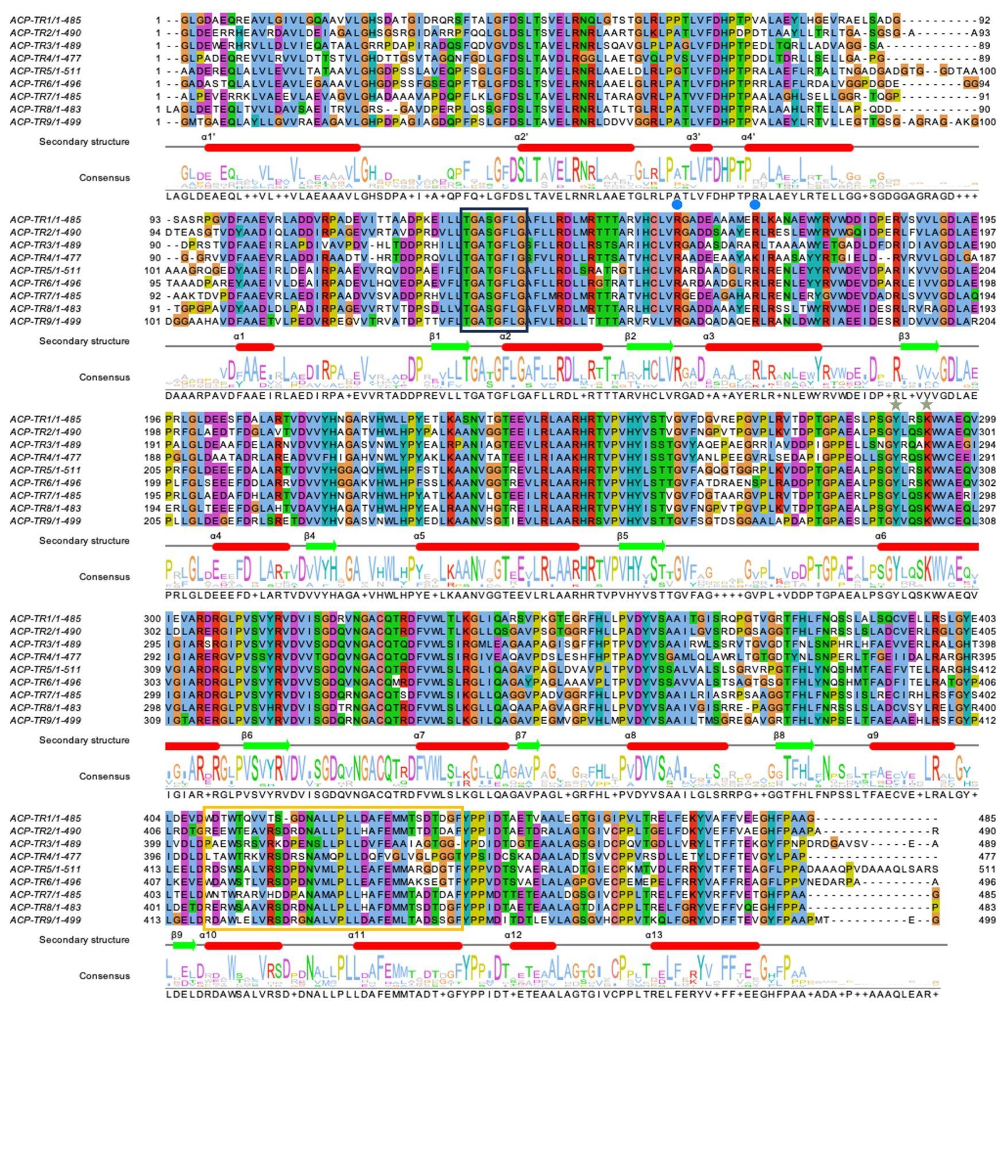

### **Supplementary Figure 5. Multiple sequence alignment of ACP-TRs.** Conserved Arg residues responsible for NADP(H) coordination are labelled with cyan dots, and catalytic Tyr and Lys are labelled with green stars. The “TGX_2_GX_2_G” P-loop motif is in the black box, and the “helix-turn-helix” motif is in the yellow box.

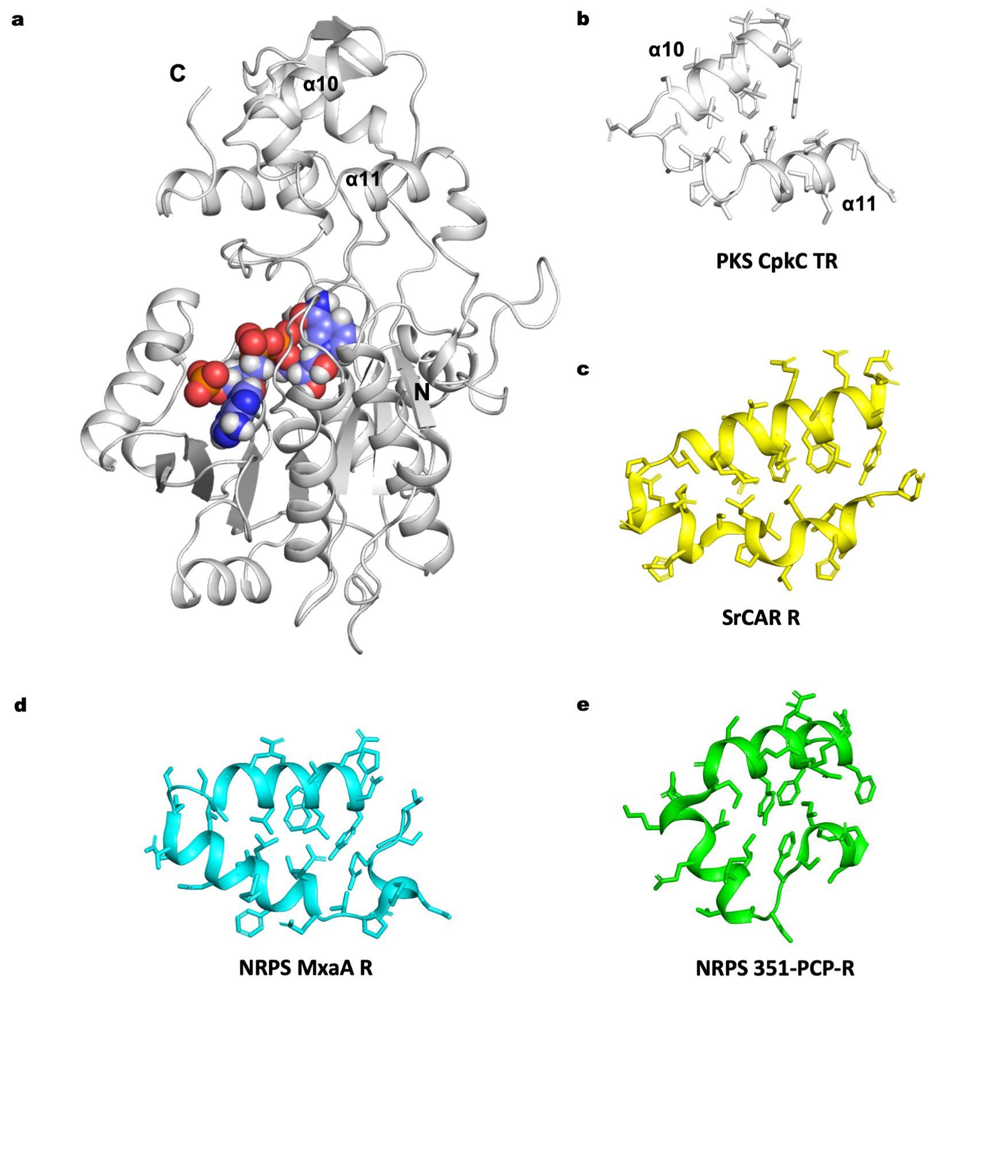

### **Supplementary Figure 6. Terminal thioreductase structures in carrier protein-containing multi-domain complexes.** A, CpkC TR (TR1) crystal structure, with the “helix-turn-helix” (HTH) motif composed of *α*10 and *α*11 labelled. B-E, The HTH motif in PKS CpkC TR (white), SrCAR R (yellow, PDB code: 5MSR), NRPS MxaA R (cyan, PDB code: 4U7W), and NPRS 351-PCP-R (green, PDB code: 6VTJ). Side chains are shown in sticks.

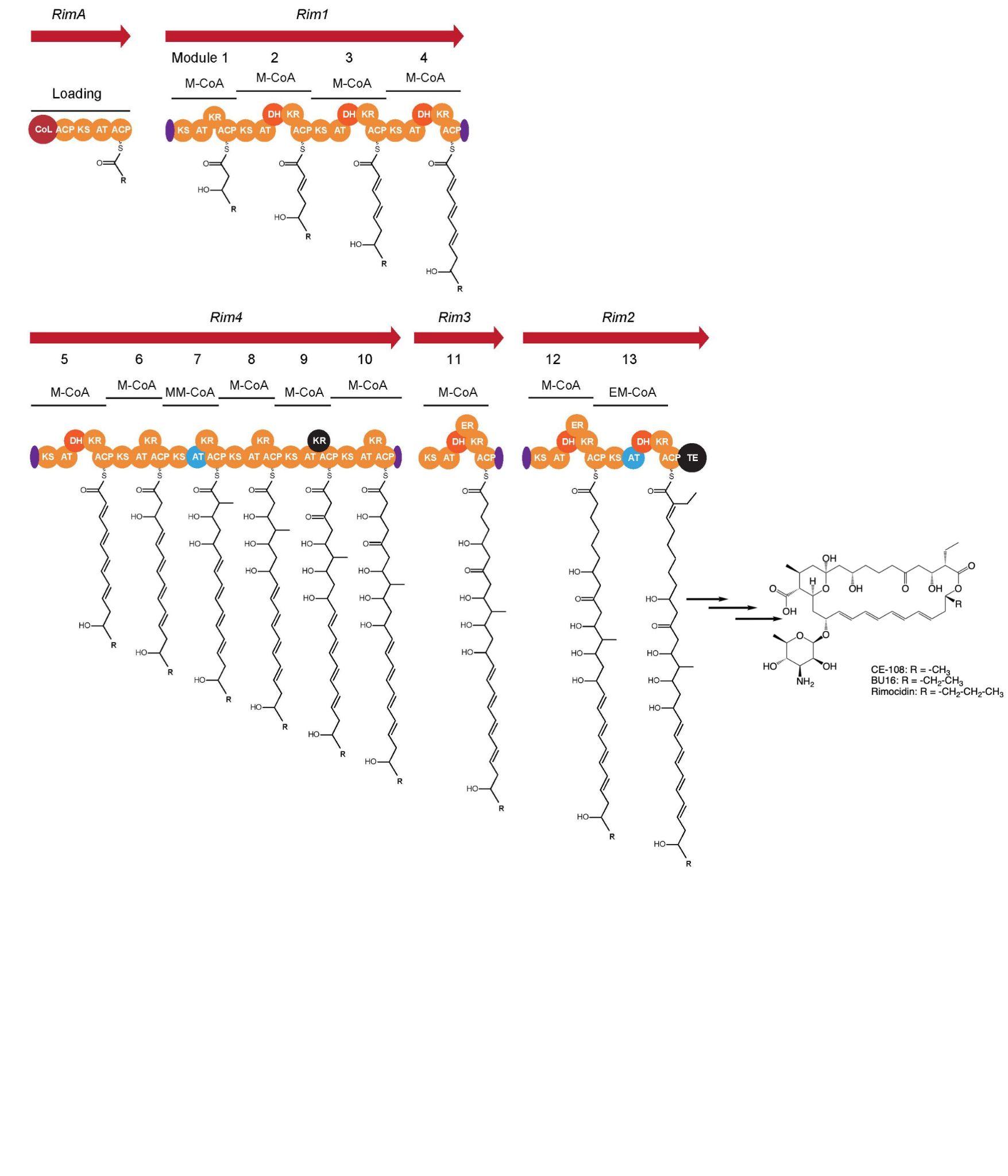

### **Supplementary Figure 7. Scheme of the rimocidin biosynthetic gene cluster.** Rimocidin loading module (RimA or RimM0) is predicted to promiscuously initiate with either acetyl-, propionyl-, or butyryl- starter units, producing CE-108, BU16, and rimocidin^8,9^ as the pathway products.

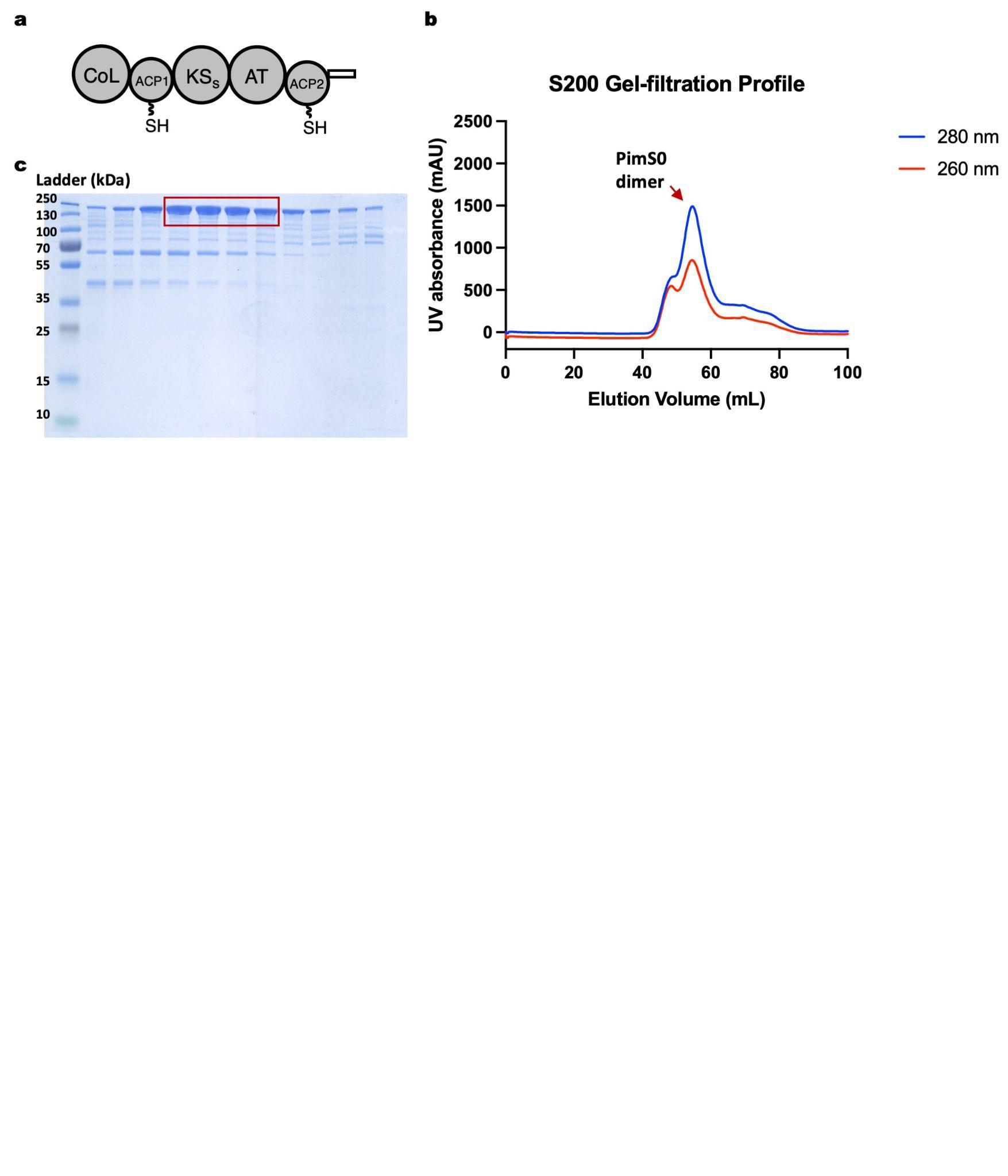

### **Supplementary Figure 8. Purification of *holo*-PimS0 (pPant-attached PimS0) loading module in *E. coli* BAP1 cells.** A. Domain organisation of PimS0 loading module^10^; B. Gel-filtration elution profile of PimS0 purification, with X-axis being elution volume, and Y-axis being UV 280 nm absorbance for the blue line and UV 260 nm absorbance for the red line. PimS0 dimer eluted at 54 min; C. SDS-PAGE analysis of PimS0 from gel-filtration. Furthermore, with coexpression of GroEL-GroES and DnaJ-DnaK-GrpE chaperones in *E. coli* BL21(DE3), we successfully purified the *apo*-PimS0 dimer to 80% homogeneity. Next we tested whether PimS0 can load acetate, malonate, acetyl-CoA, or malonyl-CoA *in vitro* (Supplementary Figure 9). *Holo*-PimS0 loaded two of the tested substrates, acetyl-CoA and malonyl-CoA. In comparison, neither acetate nor malonate was accepted by *apo*-PimS0 despite the presence of a putative CoA-ligase (CoL) domain. Both PimS0 ACP1 and ACP2 were functional, as demonstrated by detection of acetyl-ACP1 and acetyl-ACP2. Moreover, malonyl-ACP1 and malonyl-ACP2 were detected after 1 hour of malonyl-CoA incubation, but absent after 4 hours, suggesting that KSs may be capable of malonyl-ACP decarboxylation to generate acetyl-ACPs. The fact that PimS0 readily accepted acetyl-CoA and malonyl-CoA as substrates suggested a similar initiation mode for RimM0 with a putatively altered substrate pool, which may include acetyl-CoA, propionyl-CoA, and butyryl-CoA starter units.

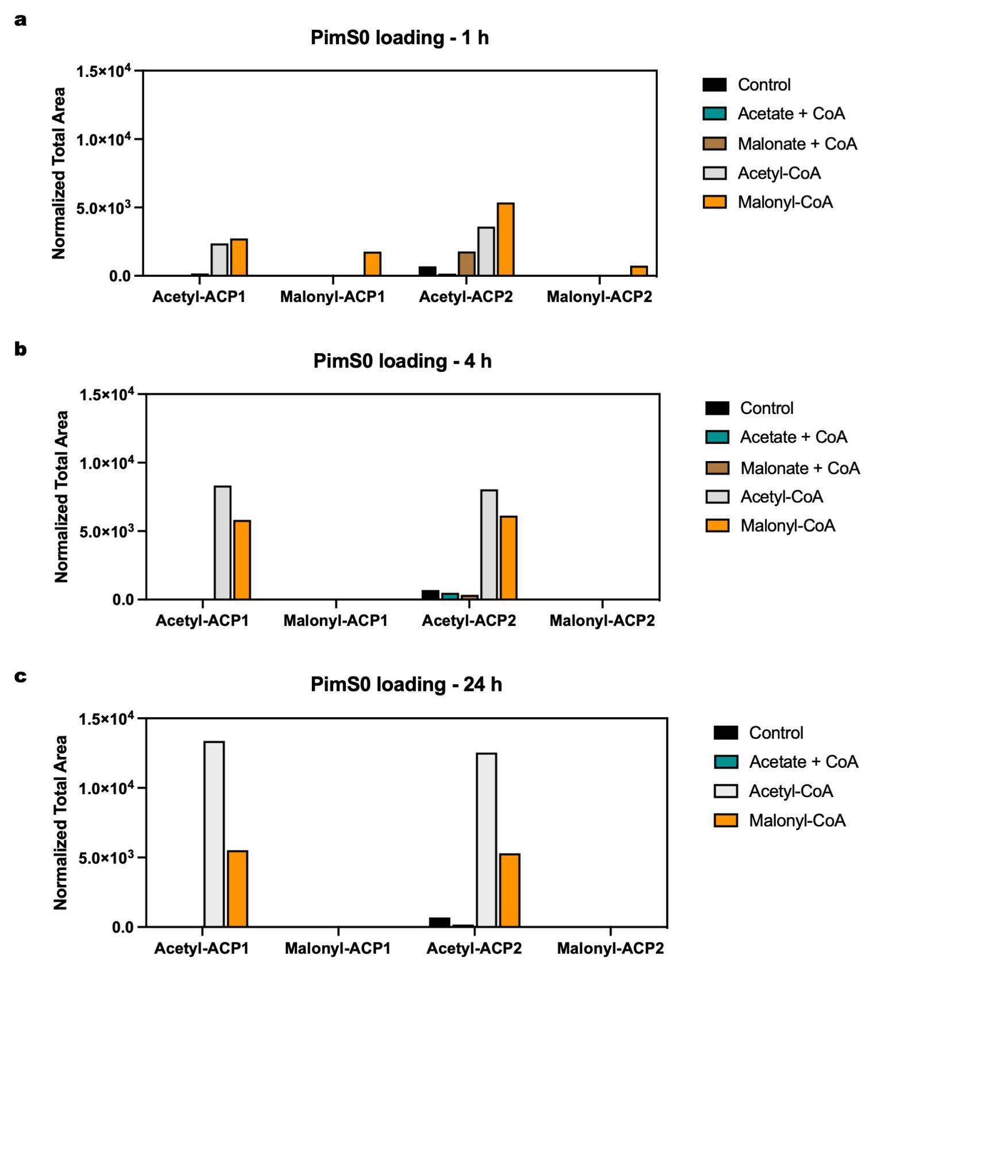

### **Supplementary Figure 9. Substrate scope analysis of PimS0 loading module.** Black: no substrate control; Cyan: acetate + CoA; Brown: malonate + CoA; White: acetyl-CoA; Orange: malonyl-CoA.

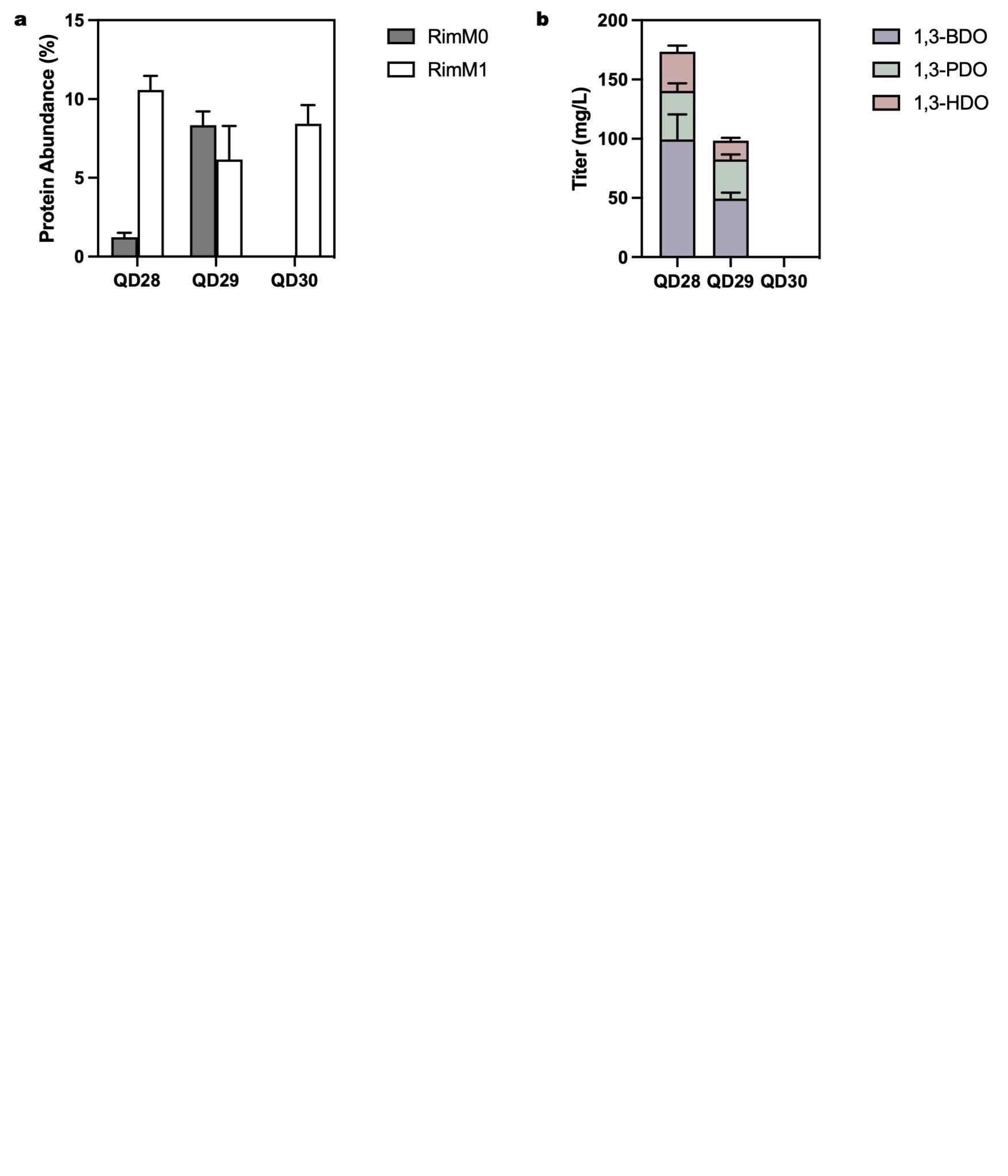

### **Supplementary Figure 10. Promoter screening results of *rimM0* in *S. albus* RimM0M1-TR2 (QD28-QD30).** QD28: *S. albus PrpsL(RO)-rimM0* + *Pgapdh(EL)-rimM1-TR2*; QD29: *S. albus Pgapdh(EL)-rimM0* + *Pgapdh(EL)-rimM1-TR2*; QD30: *S. albus kasOP***-rimM0* + *Pgapdh(EL)-rimM1-TR2*. A. Protein abundance of RimM0 and RimM1 in engineered *S. albus* when using *PrpsL(RO)*, *Pgapdh(EL)*, or *kasOP** promoter to drive *rimM0* transcription. The *rimM1-TR2* promoter is *Pgapdh(EL)*. Grey: RimM0 protein abundance; White: RimM1 protein abundance; B. 1,3-Diol production titers after 7 d cultivation in R5 medium. Purple: 1,3-BDO; Green: 1,3-PDO; Red: 1,3-HDO. Each experiment was repeated in quadruplicate.

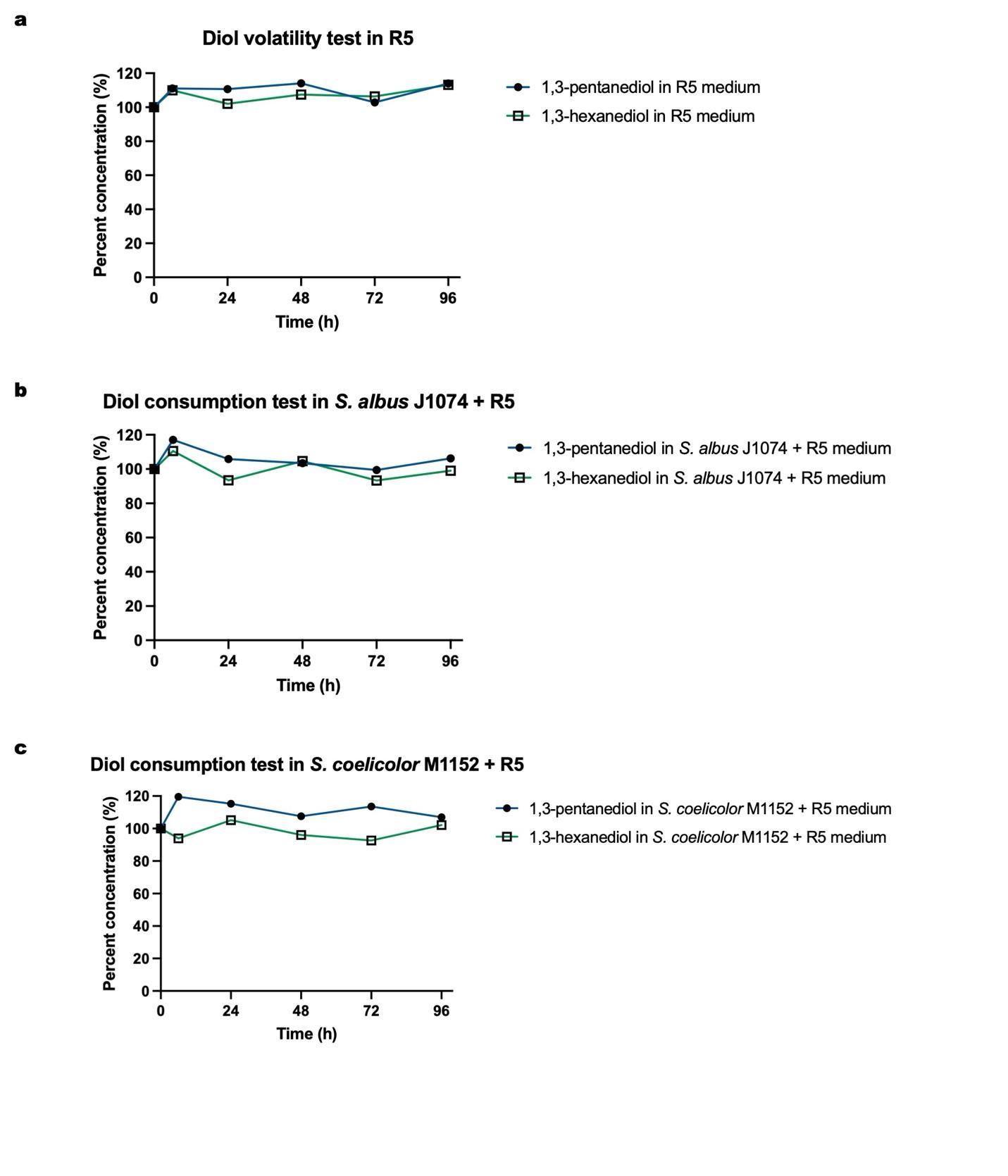

### **Supplementary Figure 11. 1,3-Diol volatility and consumption test.** A. Volatility test of 1,3-pentanediol (blue) and 1,3-hexanediol (green) in R5 medium; B. *S. albus* J1074 consumption test of 1,3-pentanediol (blue) and 1,3-hexanediol (green) in R5 medium; C. *S. coelicolor* M1152 consumption test of 1,3-pentanediol (blue) and 1,3-hexanediol (green) in R5 medium.

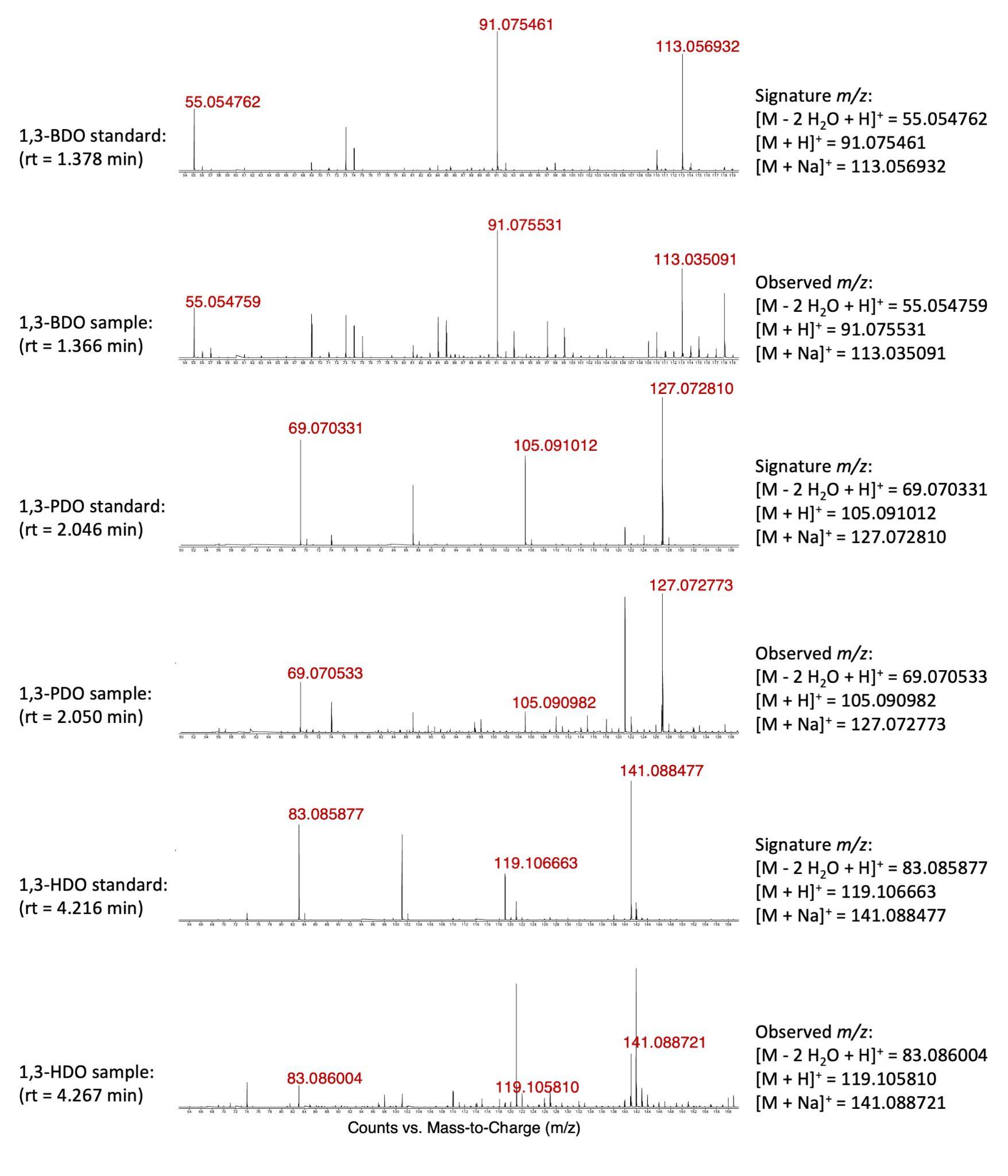

### **Supplementary Figure 12. MS spectra of 1,3-diols produced in *S. albus* RimM0M1-TR1 (QD27).** From top to bottom are MS spectrums of 1,3-BDO standard at rt = 1.378 min, QD27 at rt = 1.366 min, 1,3-PDO standard at rt = 2.046 min, QD27 at rt = 2.050 min, 1,3-HDO standard at rt = 4.216 min, QD27 at rt = 4.267 min. All observed *m/z* match well with expected signature *m/z* (dehydrated *m/z*^11^, [M + H]^+^ *m/z*, and [M + Na]^+^ *m/z*).

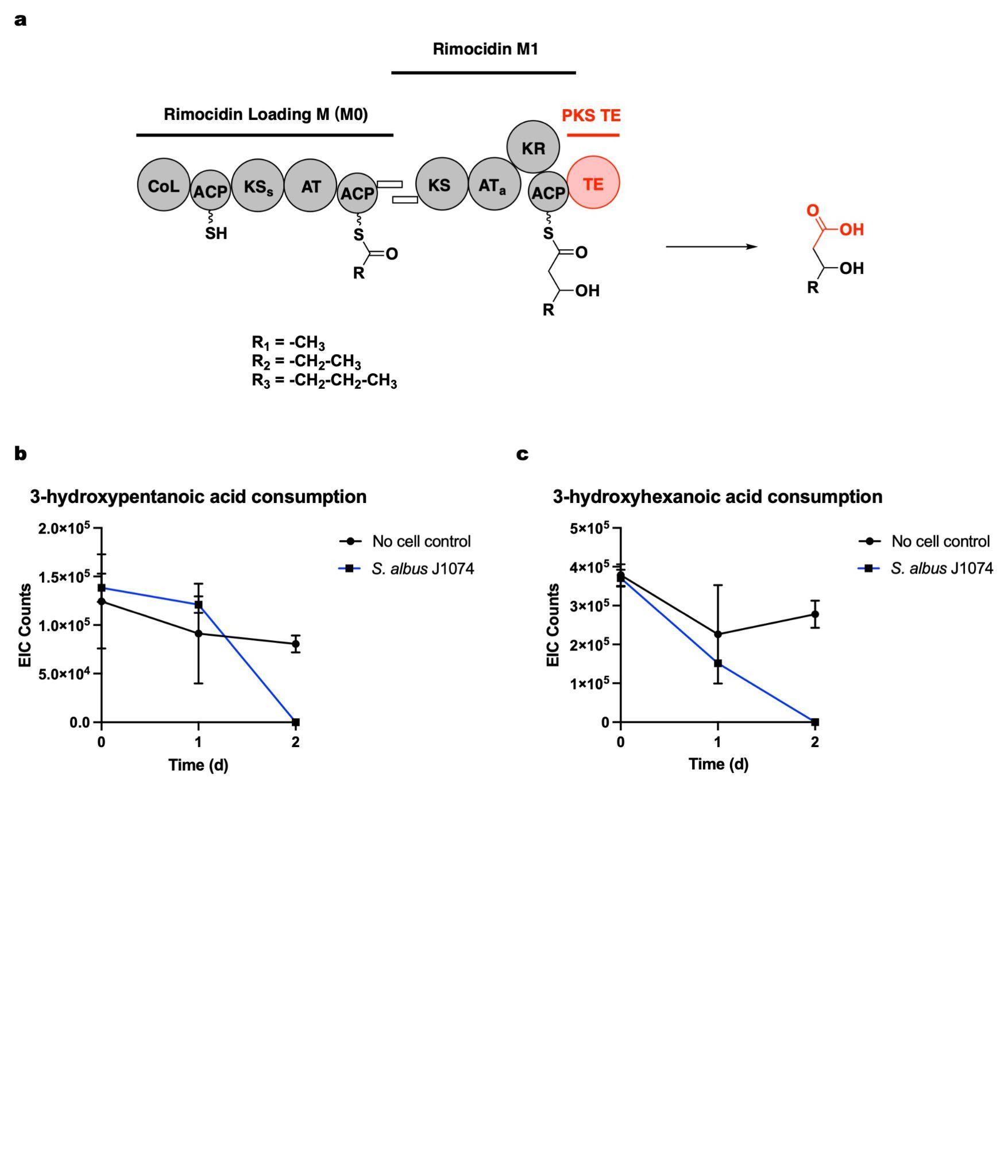

### **Supplementary Figure 13. *S. albus* RimM0M1-DEBS TE (QD18) production test.** A. Reaction scheme of RimM0M1-DEBS TE for 3-hydroxy acid production in *Streptomyces*. No 3-hydroxy acid was detected; B. Consumption test of 3-hydroxypentanoic acid in R5 (black) or R5 + *S. albus* (blue); C. Consumption test of 3-hydroxyhexanoic acid in R5 (black) or R5 + *S. albus* (blue). Each experiment was repeated in triplicate.

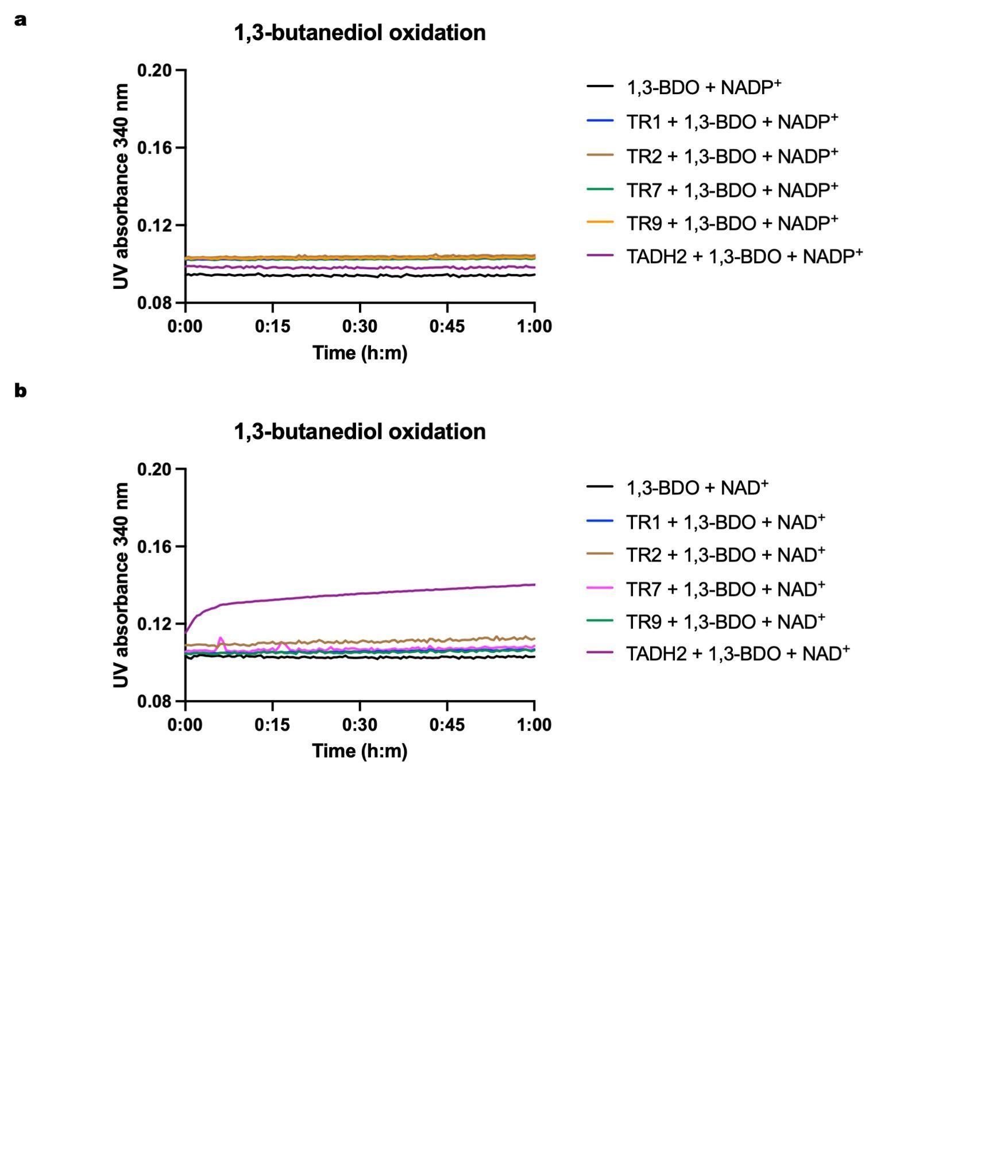

### **Supplementary Figure 14. TR substrate scope analysis with 1,3-butanediol, using NADP^+^ (a) or NAD^+^ (b) as the cofactor.** Black: no enzyme control; Blue: TR1; Brown: TR2; Magenta: TR7; Green: TR9; Purple: TADH2 positive control. Each experiment was repeated in triplicate.

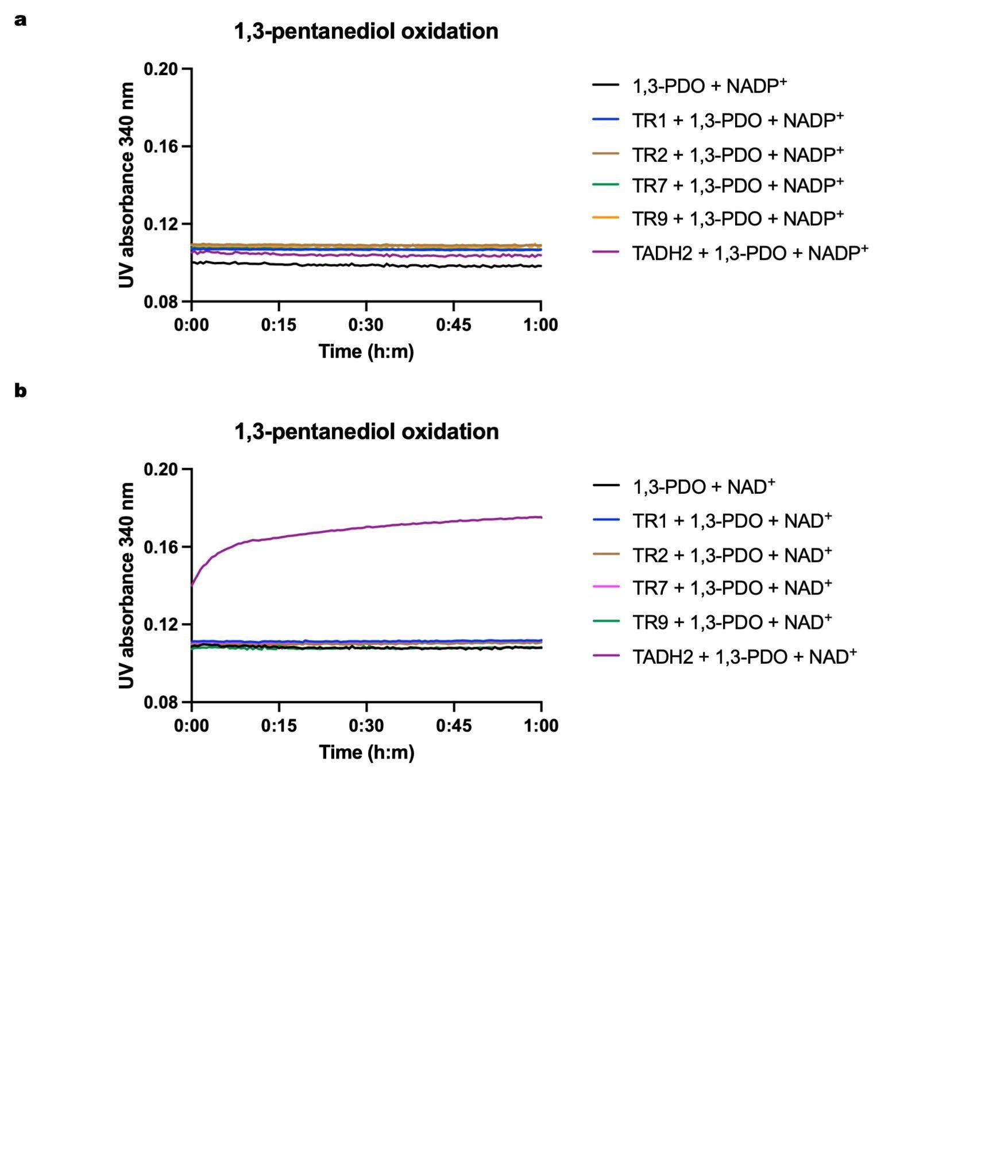

### **Supplementary Figure 15. TR substrate scope analysis with 1,3-pentanediol, using NADP^+^ (a) or NAD^+^ (b) as the cofactor.** Black: no enzyme control; Blue: TR1; Brown: TR2; Magenta: TR7; Green: TR9; Purple: TADH2 positive control. Each experiment was repeated in triplicate.

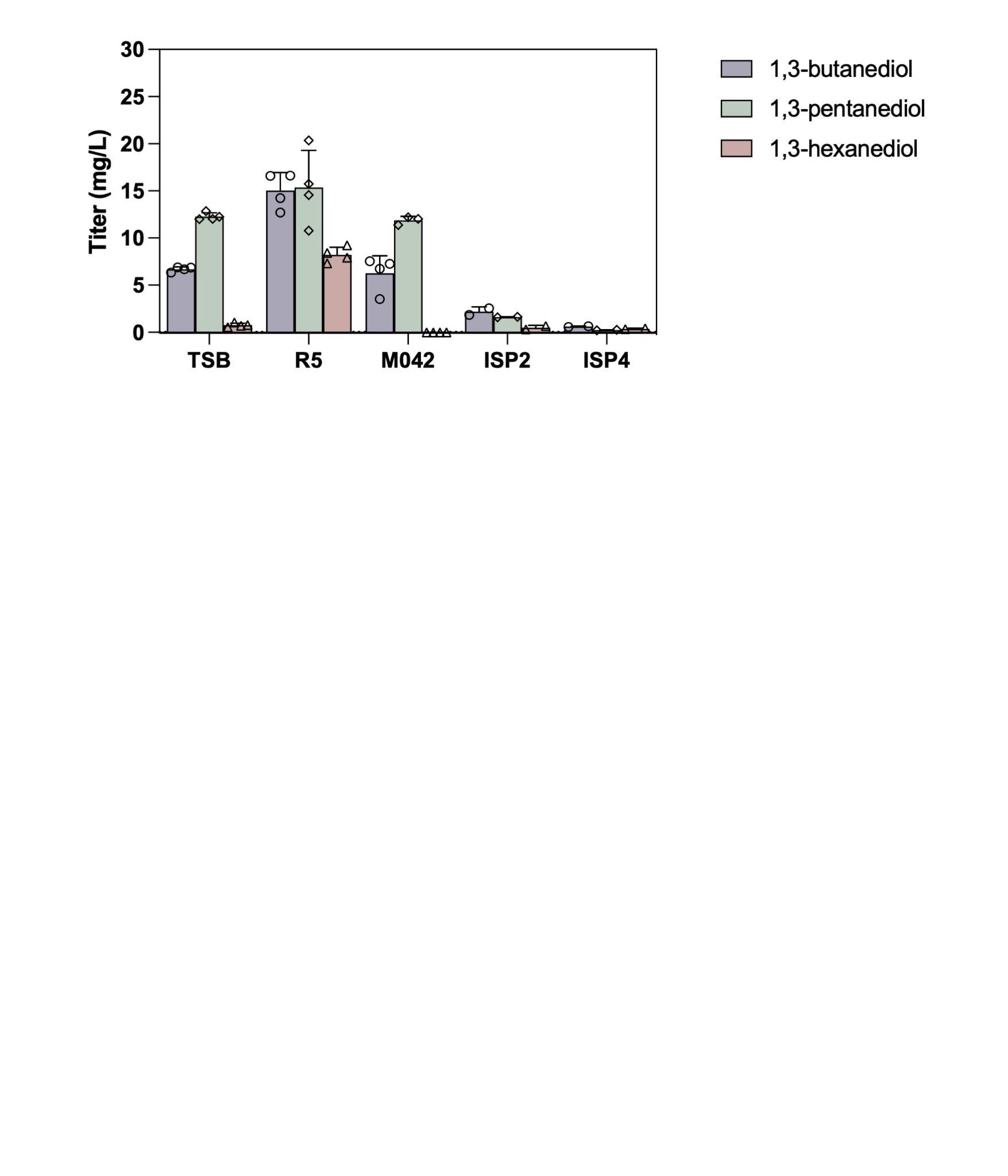

### **Supplementary Figure 16. Medium screening for *S. albus* RimM0M1-TR1 (QD27).** 3 d RimM0M1-TR1 cultivation results in TSB, R5, M042, and 5 d cultivation results in ISP2, ISP4 were presented here. 15.0 mg/L 1,3-BDO, 15.4 mg/L 1,3-PDO, and 8.2 mg/L 1,3-HDO were produced in R5 after 3 d. Each experiment was repeated in at least duplicate.

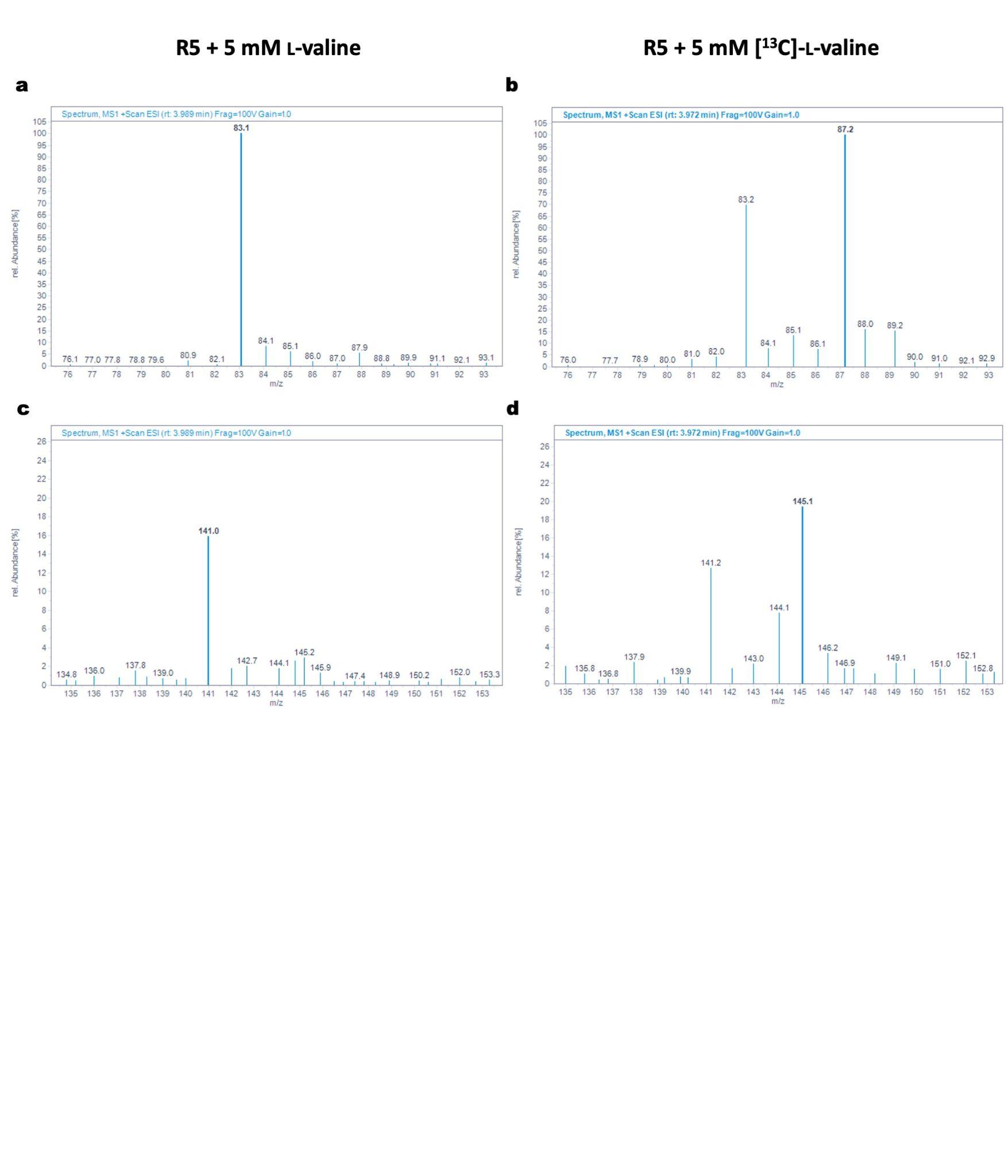

### **Supplementary Figure 17. [^13^C]labelled L-valine feeding experiment on *S. albus* RimM0M1-TR2 (QD28).** R5 medium + 5 mM L-valine or [^13^C]labelled L-valine was selected for comparison. A. 1,3-hexanediol MS spectrum in L-valine addition group (rt = 3.99 min, expected dehydrated 83.1 *m/z*, observed 83.1 *m/z*); B. 1,3-hexanediol MS spectrum in [^13^C]labelled L-valine addition group (rt = 3.97 min, expected dehydrated 87.1 *m/z*, observed 87.1 *m/z*); C. 1,3-hexanediol MS spectrum in L-valine addition group (rt = 3.99 min, expected [M + Na]^+^ 141.1 *m/z*, observed 141.0 *m/z*); D. 1,3-hexanediol MS spectrum in [^13^C]labelled L-valine addition group (rt = 3.97 min, expected [M + Na]^+^ 145.1 *m/z*, observed 145.1 *m/z*).

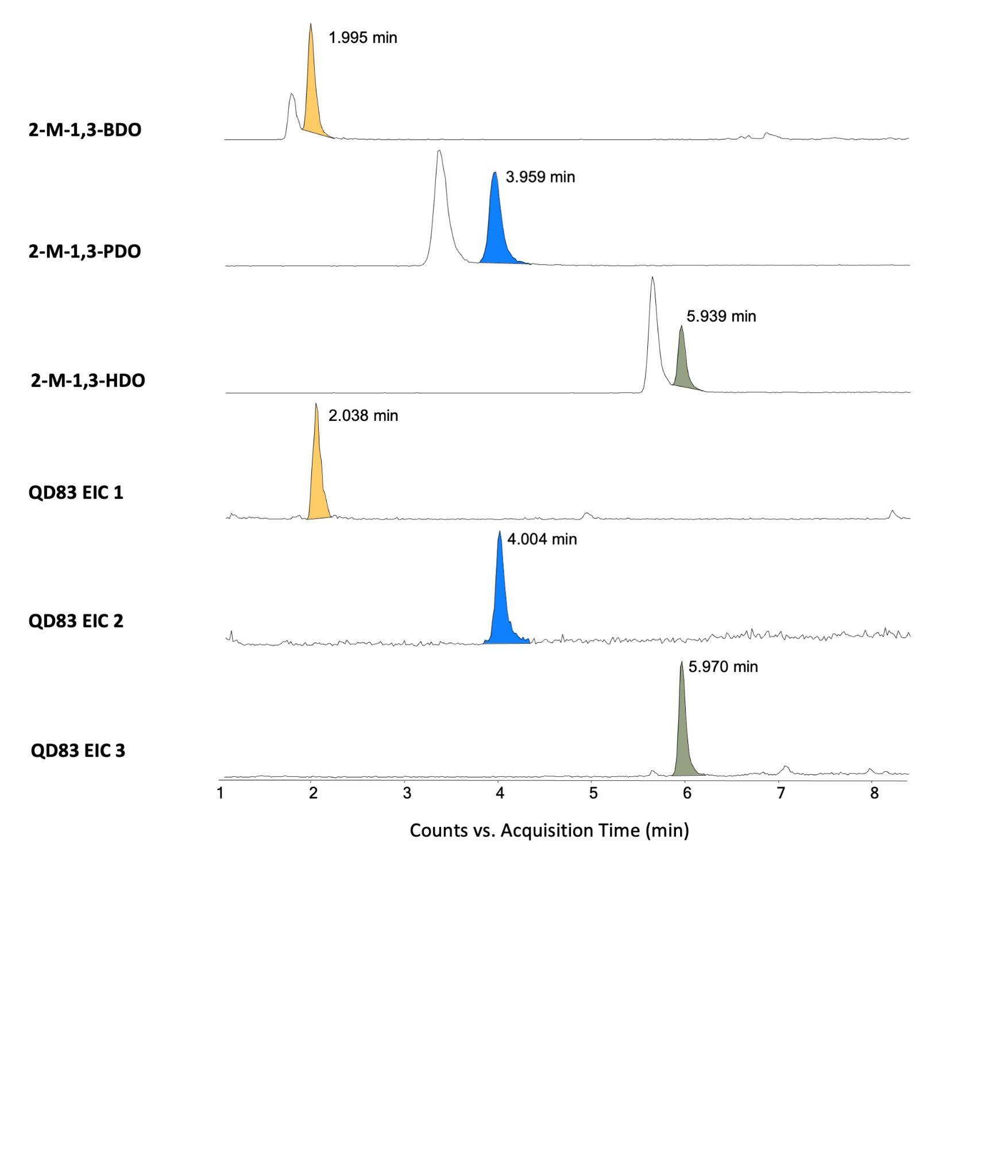

### **Supplementary Figure 18. LC-MS detection of 2-methyl-1,3-diols produced in *S. albus* RimM0M1(RimM7 AT)-TR2 + FkbS (QD83).** From top to bottom: 2-M-1,3-BDO standard EIC ([M + H]^+^ = 105.091006 *m/z*); 2-M-1,3-PDO standard EIC ([M + Na]^+^ = 141.088601 *m/z*); 2-M-1,3-HDO standard EIC ([M + Na]^+^ = 155.104251 *m/z*); QD83 EIC 1 = 105.091006 *m/z*; EIC 2 = 141.088601 *m/z*; EIC 3 = 155.104251 *m/z*.

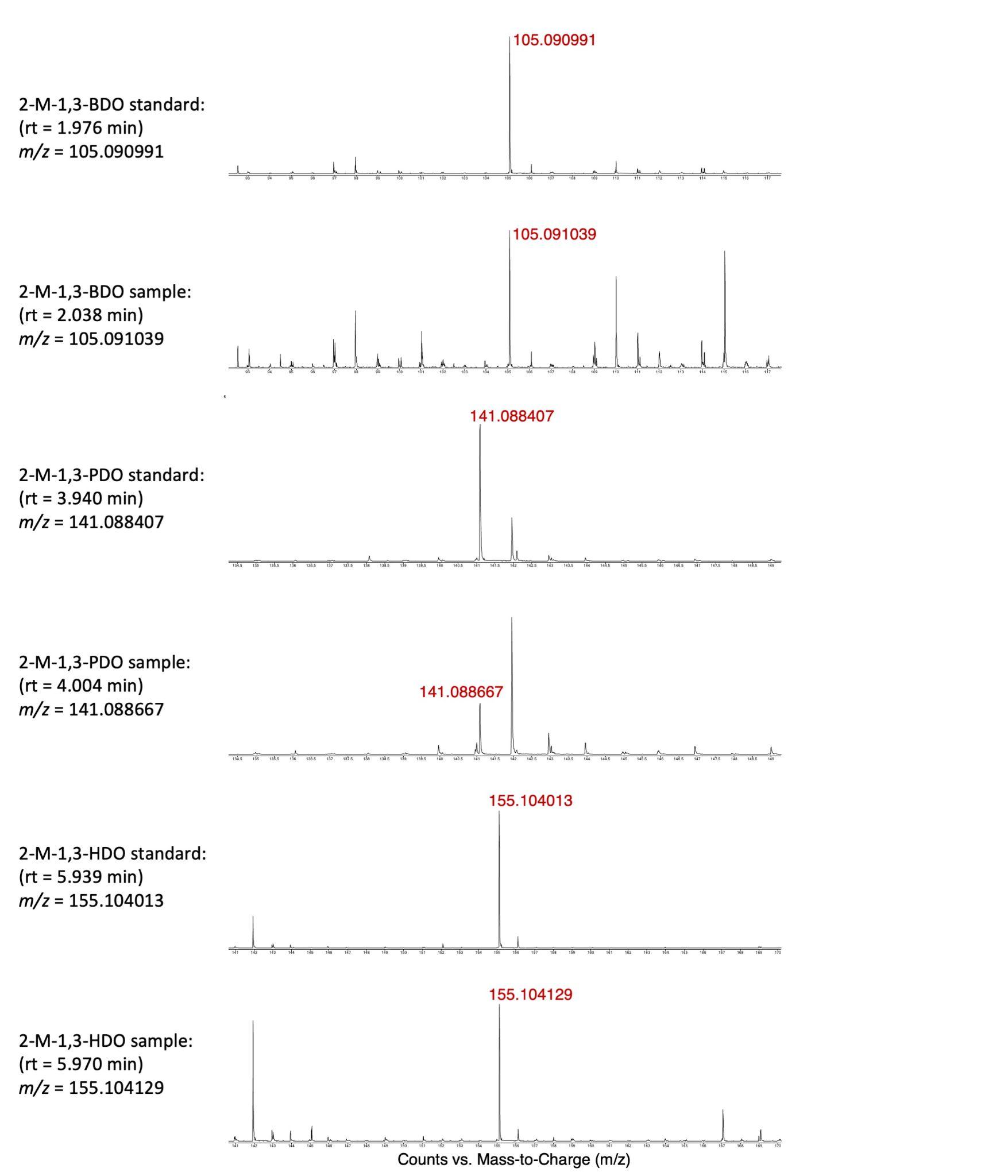

### **Supplementary Figure 19. MS spectra of 2-methyl-1,3-diols produced in *S. albus* RimM0M1(M7 AT)-TR2 + FkbS (QD83).** [M + H]^+^ *m/z* were presented for 2-M-1,3-BDO standard and sample, and [M + Na]^+^ *m/z* were shown here for 2-M-1,3-PDO and 2-M-1,3-HDO standards and samples.

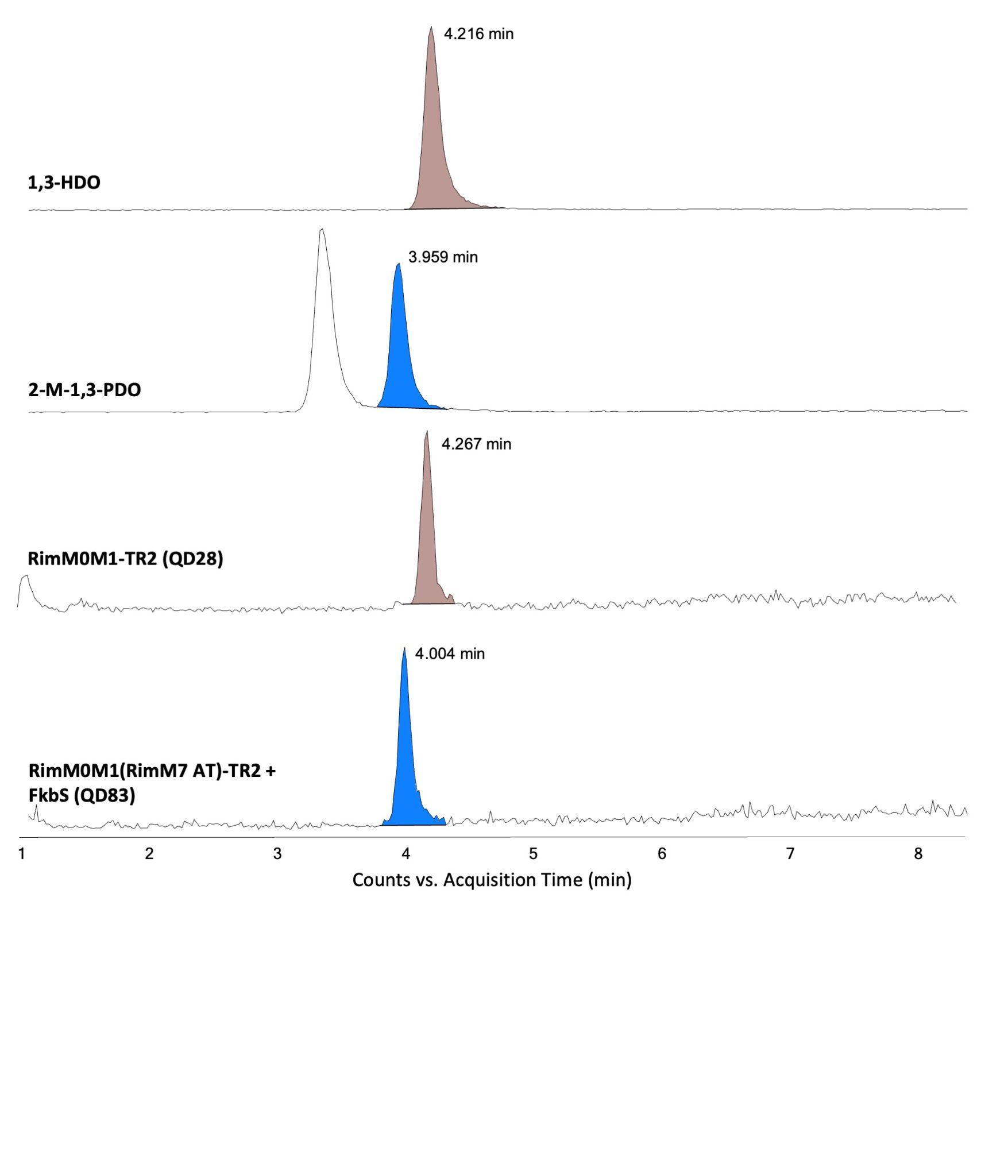

### **Supplementary Figure 20. Comparative analysis of 1,3-diols produced in *S. albus* QD28 and QD83.** From top to bottom are EIC traces of 1,3-HDO standard, 2-M-1,3-PDO standard, *S. albus* RimM0M1-TR2 (QD28) sample, and *S. albus* RimM0M1(RimM7 AT)-TR2 + FkbS (QD83) sample. 1,3-HDO and 2-M-1,3-PDO are isomers ([M + Na]^+^ = 141.088601 *m/z*). 1,3-HDO was produced only in QD28, whereas 2-M-1,3-PDO was produced only in QD83, confirming precision of PKS-TR engineering.

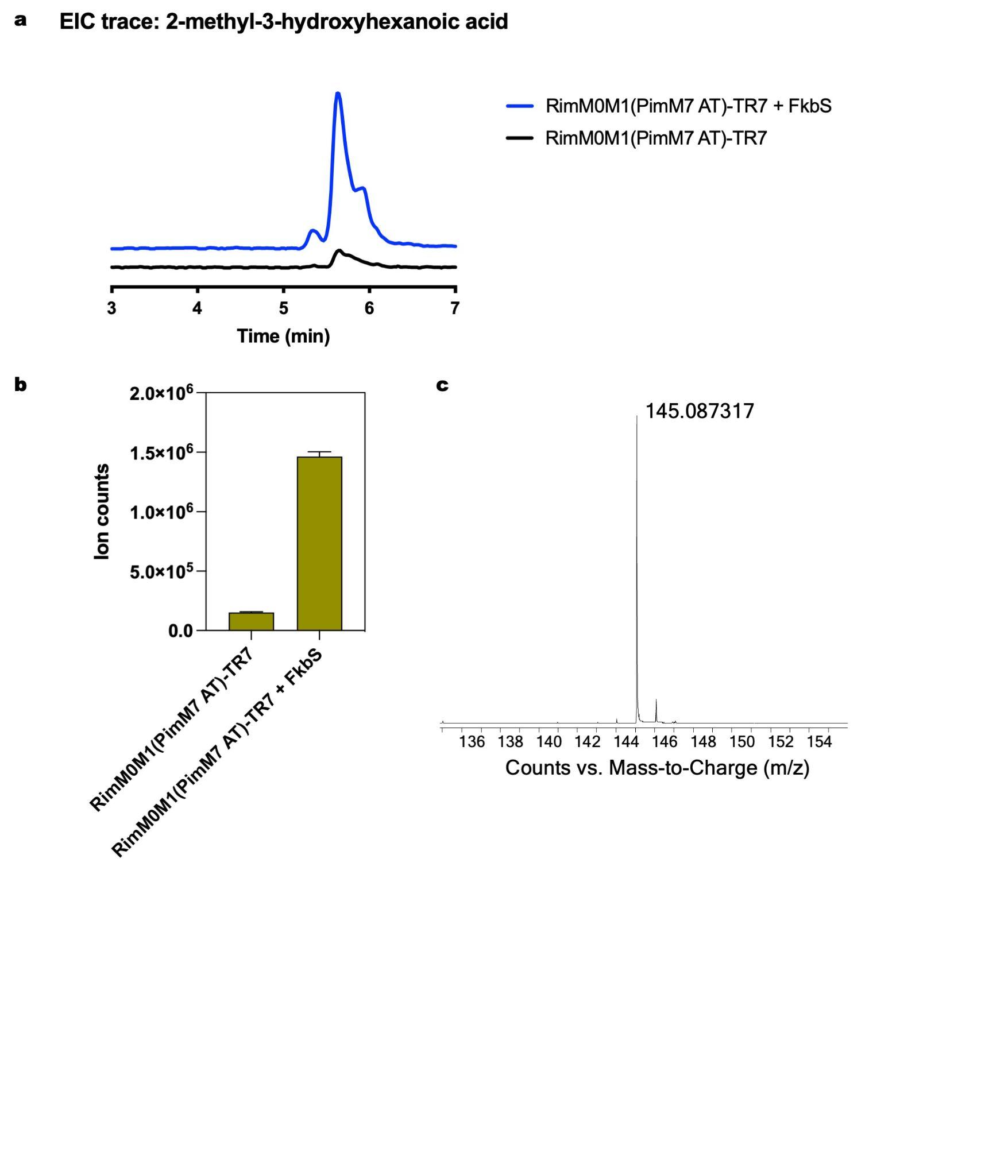

### **Supplementary Figure 21. Detection of 2-methyl-3-hydroxyhexanoic acid produced in *S. albus*.** A, EIC profiles of 2-methyl-3-hydroxyhexanoic acid: 7 d cultivation of RimM0M1(PiM7 AT)-TR7 (QD69) in R5 + 2% Glucose + 15 mM L-valine is shown in black, and 7 d cultivation of RimM0M1(PiM7 AT)-TR7 + FkbS (QD85) in R5 + 2% Glucose + 15 mM L-valine is shown in blue; B, Production profiles of 2-methyl-3-hydroxyhexanoic acid; C. Observed [M - H]^-^ of 2-methyl-3-hydroxyhexanoic acid in the QD85 sample is 145.087317 *m/z* (theoretical [M - H]^-^ 145.087018 *m/z*, mass error 2.06 ppm).

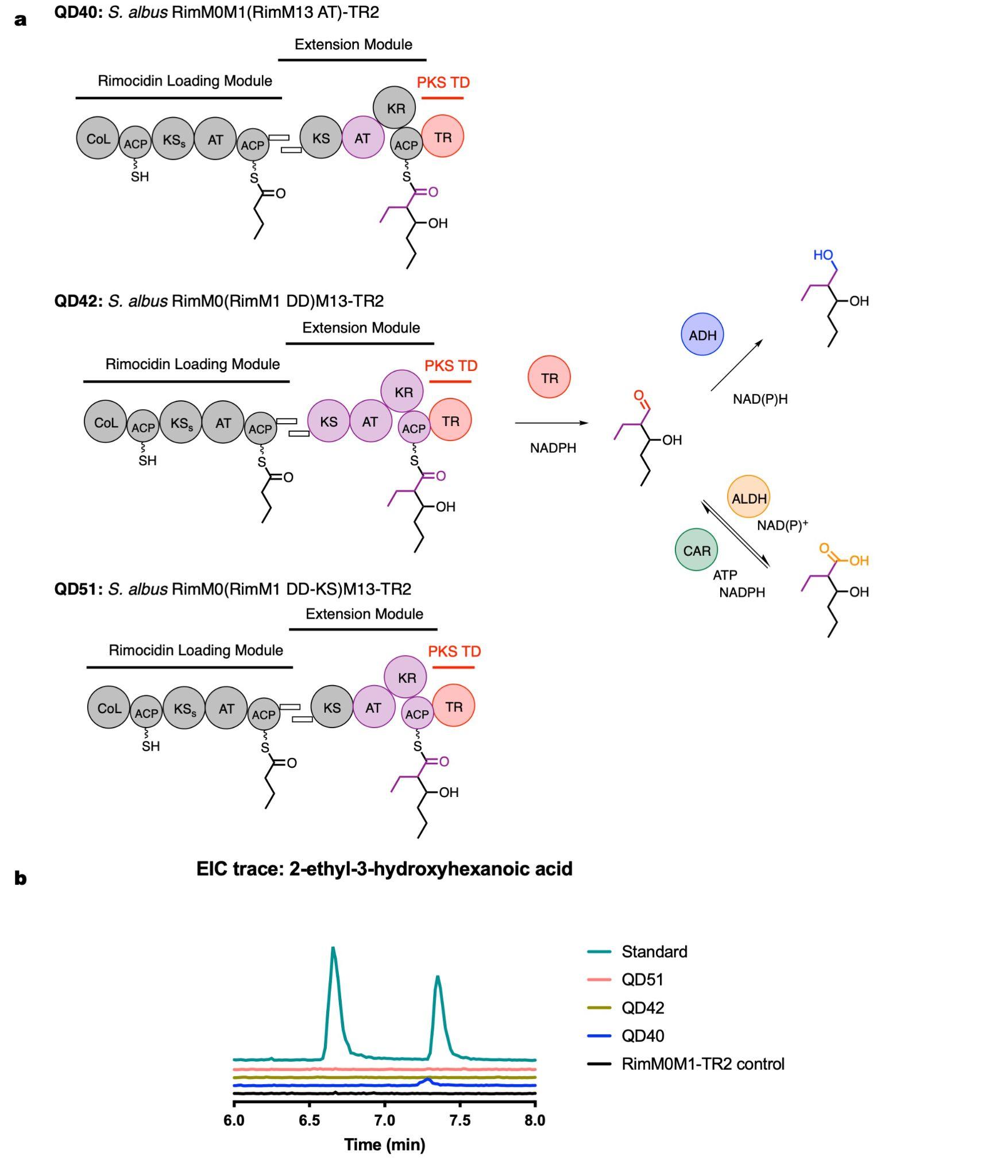

### **Supplementary Figure 22. Detection of 2-ethyl-3-hydroxyhexanoic acid produced in *S. albus*.** A, three PKS pathway designs for 2-ethyl-3-hydroxyhexanoic acid production were tested in *S. albus*. AT: acyltransferase; DD: docking domain; KS: ketosynthase; B, EIC profiles of 2-ethyl-3-hydroxyhexanoic acid in QD28 control (black), QD40 (blue), QD42 (brown), QD51 (pink) after 3 d cultivation in R5 + 2% Glucose + 5 mM L-valine, and 2-ethyl-3-hydroxyhexanoic acid standard (cyan).

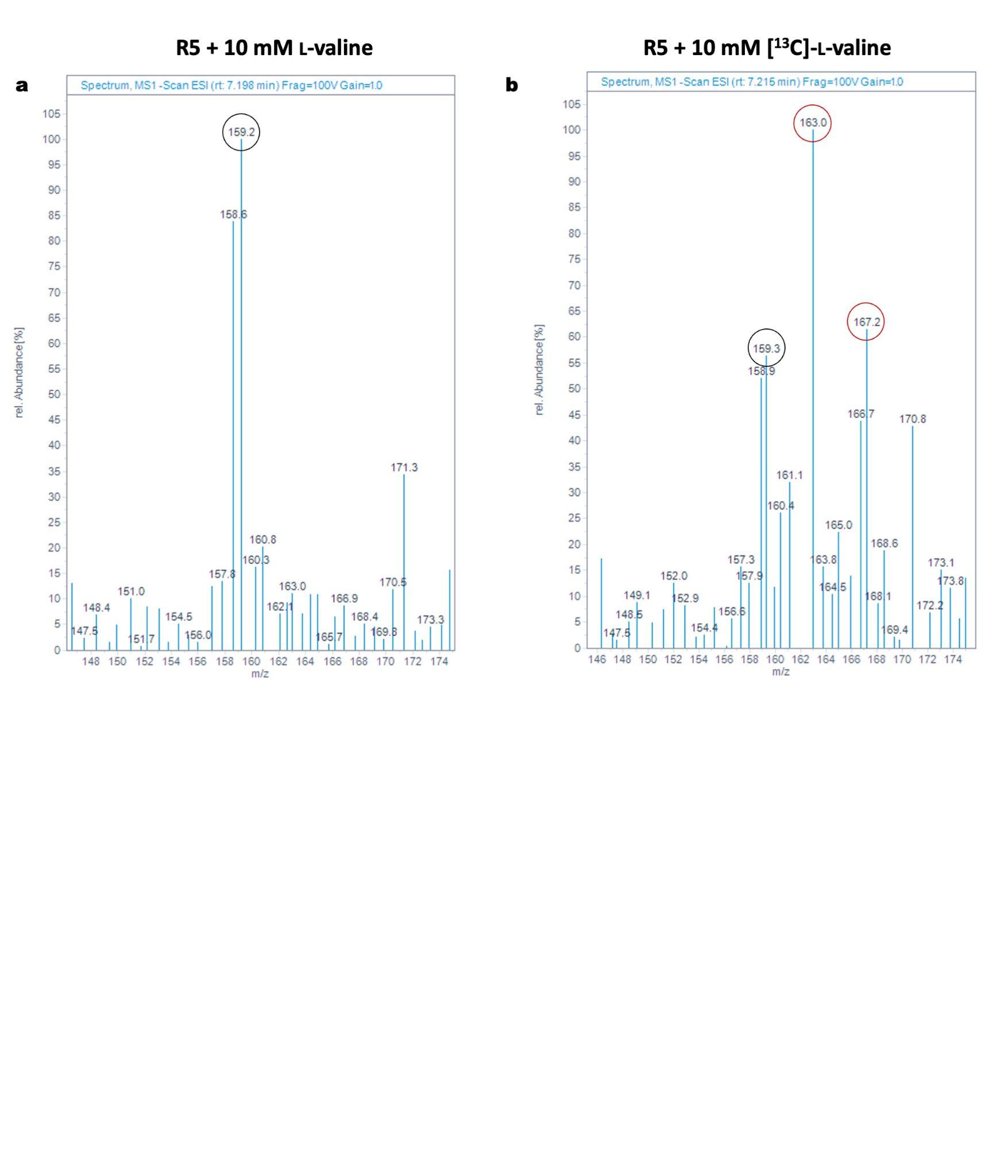

### **Supplementary Figure 23. [^13^C]labelled L-valine feeding experiment on *S. albus* RimM0M1(RimM13 AT)-TR2 (QD40).** R5 medium + 10 mM L-valine or [^13^C]labelled L-valine was selected for comparison. A. 2-ethyl-3-hydroxyhexanoic acid MS spectrum in L-valine addition group (rt = 7.20 min, expected [M - H]^-^ 159.2 *m/z*, observed 159.2 *m/z* in the black circle); B. 2-ethyl-3-hydroxyhexanoic acid MS spectrum in [^13^C]labelled L-valine addition group (rt = 7.21 min, expected [M - H]^-^ 163.2 *m/z* for a single [^13^C]labelled L-valine incorporation and 167.2 *m/z* for double [^13^C]labelled L-valine incorporation, observed 163.0 *m/z* and 167.2 *m/z* in red circles).

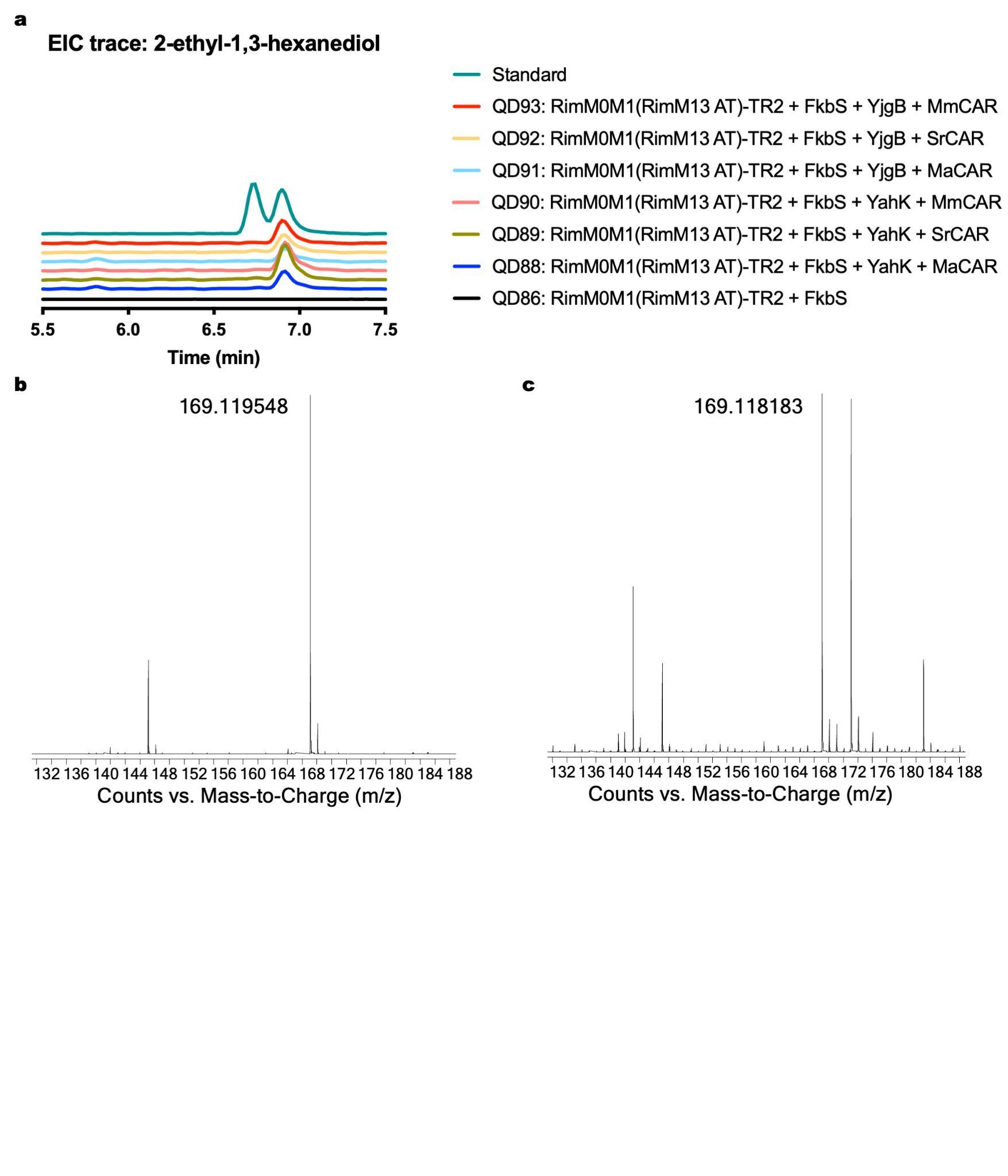

### **Supplementary Figure 24. Detection of 2-ethyl-1,3-hexanediol produced in *S. albus*.** A, EIC profiles of 2-E-1,3-HDO: 7 d cultivation of QD86, QD88 - QD93 in R5 + 2% Glucose + 15 mM L-valine; B, Observed [M + Na]^+^ *m/z* of 2-E-1,3-HDO standard is 169.119548; C. Observed [M + Na]^+^ *m/z* of 2-E-1,3-HDO in QD89 sample is 169.118183.

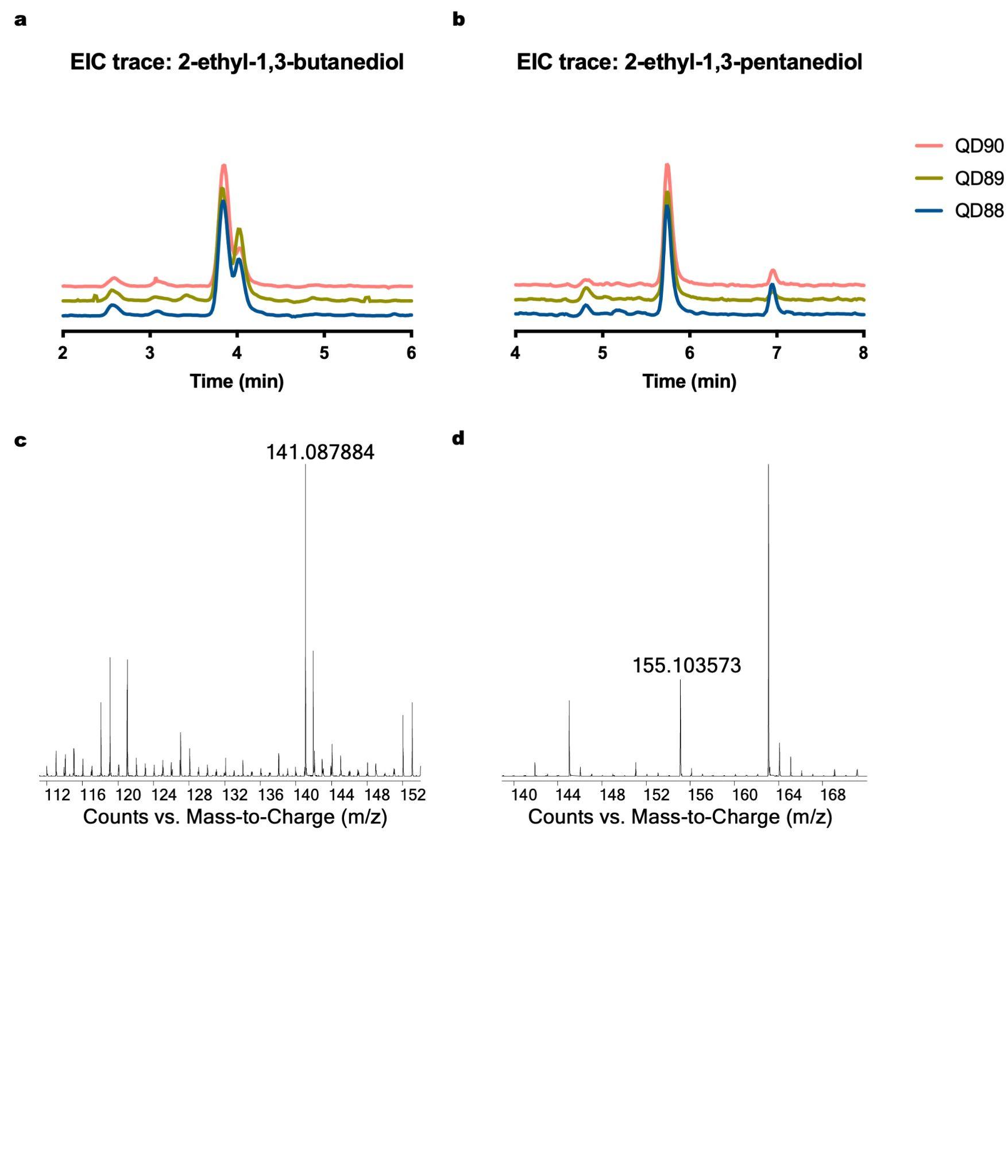

### **Supplementary Figure 25. Detection of 2-E-1,3-BDO and 2-E-1,3-PDOproduced in *S. albus*.** A, EIC profile of 2-E-1,3-BDO after 7 d cultivation of QD88 - QD90 in R5 + 2% Glucose + 15 mM L-valine; B, EIC profile of 2-E-1,3-PDO after 7 d cultivation of QD88 - QD90 in R5 + 2% Glucose + 15 mM L-valine; C, Observed [M + Na]^+^ *m/z* of 2-E-1,3-BDO in QD89 sample is 141.087884 (theoretical [M + Na]^+^ 141.088601 *m/z*, mass error -5.08 ppm); B, Observed [M + Na]^+^ *m/z* of 2-E-1,3-PDO in QD89 sample is 155.103573 (theoretical [M + Na]^+^ 155.104251 *m/z*, mass error -4.37 ppm).

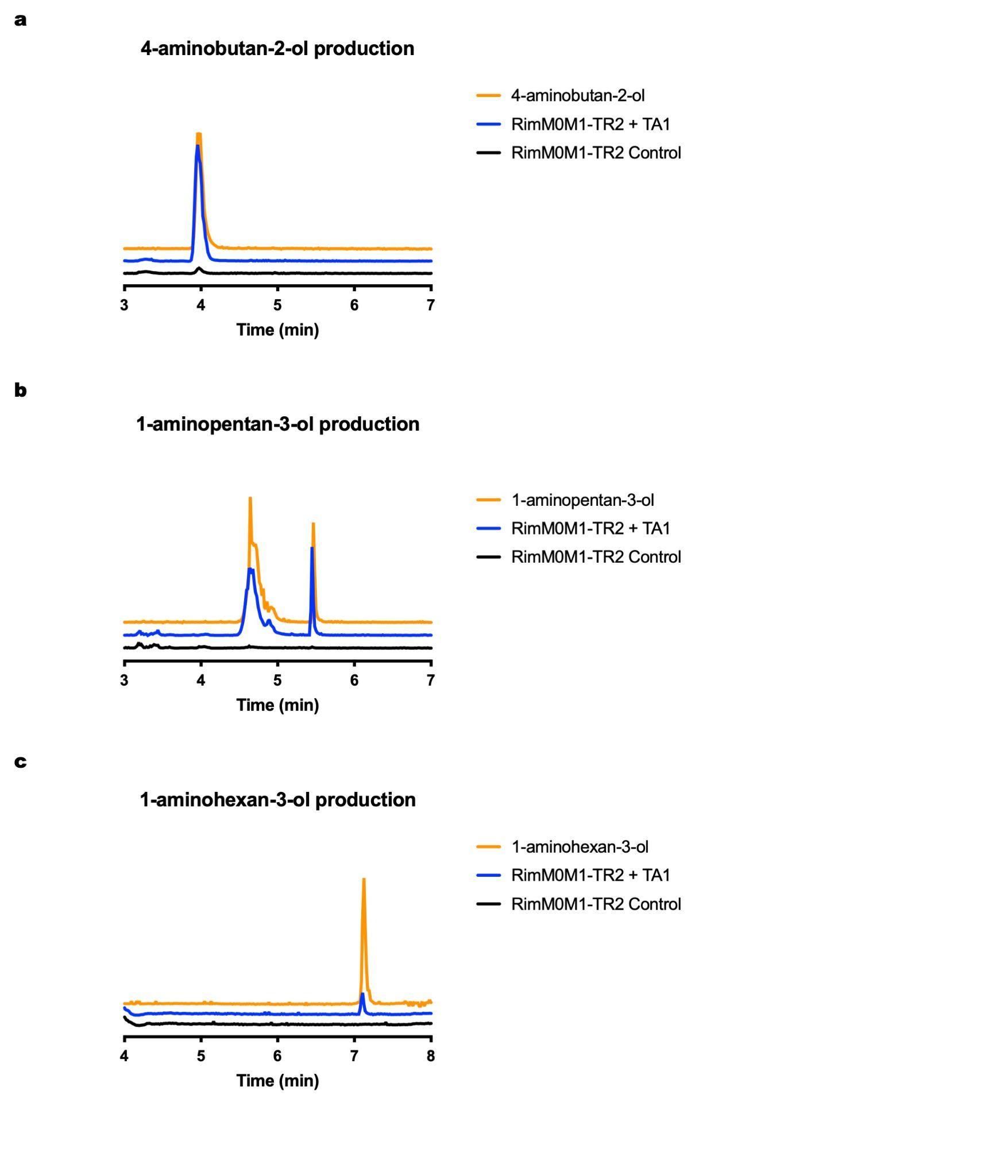

### **Supplementary Figure 26. LC-MS detection of amino alcohols produced in *S. albus* RimM0M1-TR2 + TA1 (QD96).** EIC traces: 4-aminobutan-2-ol in A, 1-aminopentan-3-ol in B, and 1-aminohexan-3-ol in C. RimM0M1-TR2 (QD28) control is coloured in black, RimM0M1-TR2 + TA1 (QD96) is coloured in blue, and the standards are coloured in orange. See Supplementary Figure 27 for detailed mass-to-charge (*m/z*) detection.

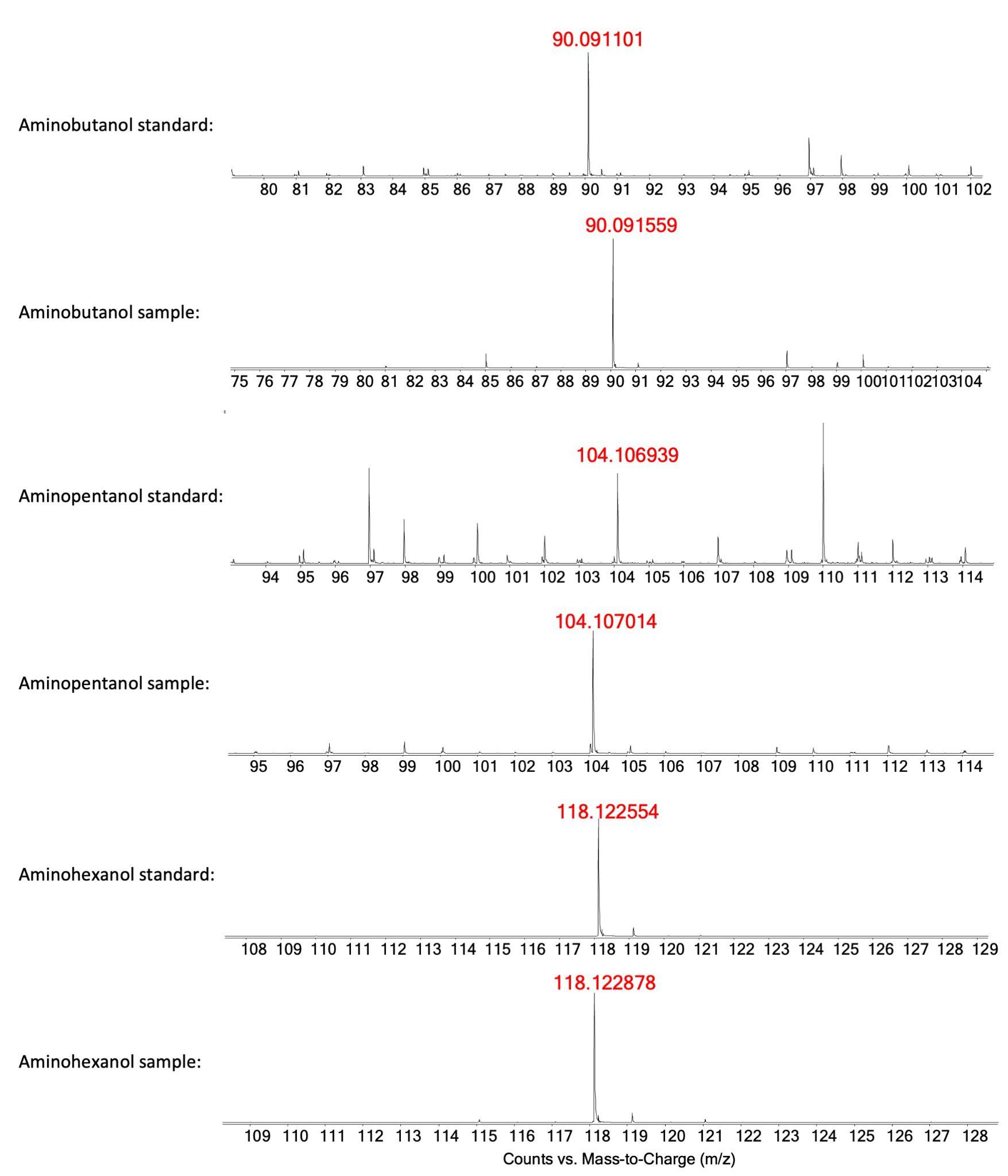

### **Supplementary Figure 27. MS spectra of amino alcohols produced in *S. albus* RimM0M1-TR2 + TA1 (QD96).** [M + H]^+^ *m/z* were presented for standards and samples of 4-aminobutan-2-ol, 1-aminopentan-3-ol, and 1-aminohexan-3-ol.

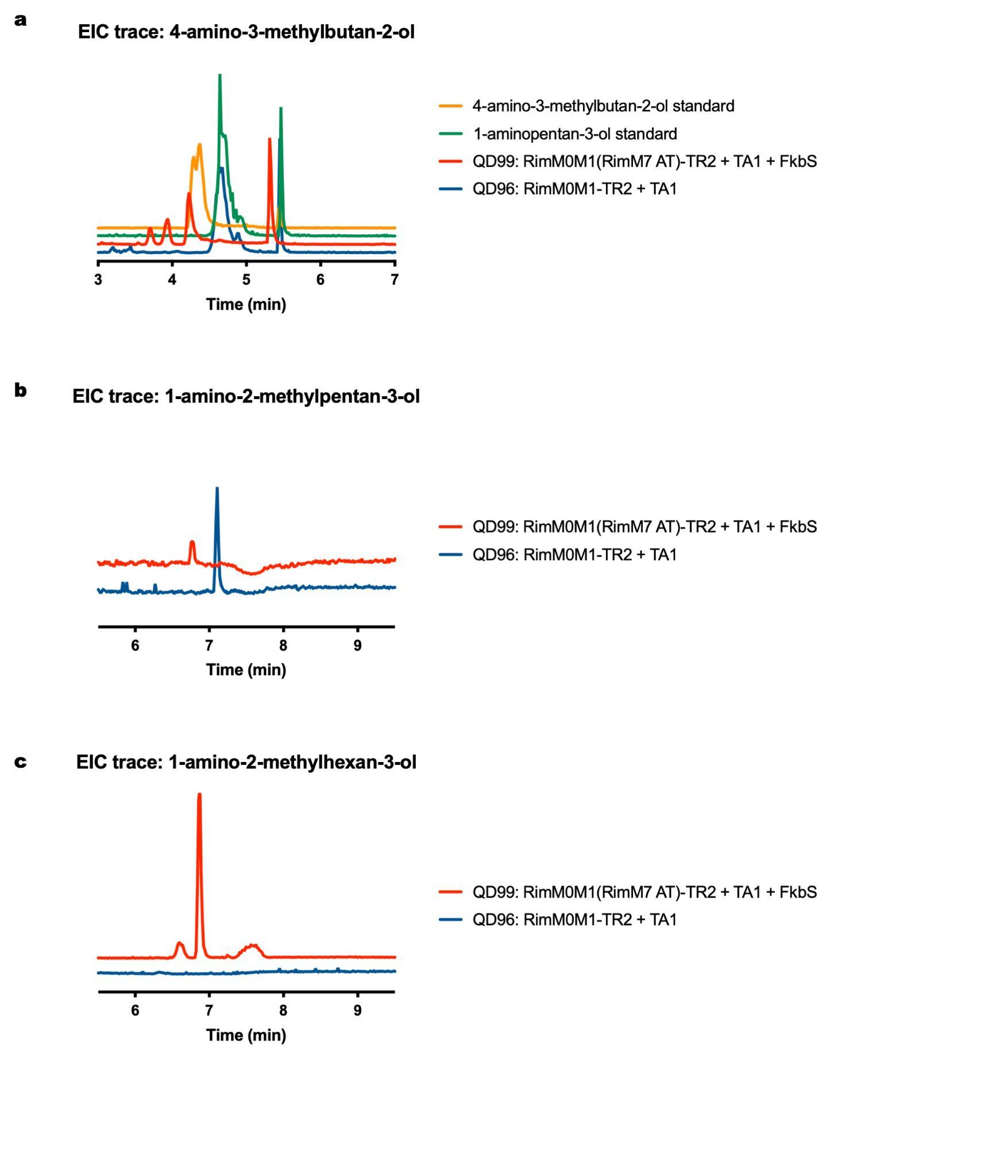

### **Supplementary Figure 28. LC-MS detection of methyl amino alcohols produced in *S. albus* RimM0M1(RimM7 AT)-TR2 + TA1 + FkbS (QD99).** *S. albus* RimM0M1-TR2 + TA1 (QD96) that produced 4-aminobutan-2-ol, 1-aminopentan-3-ol, and 1-aminohexan-3-ol serves as control. EIC traces: 4-amino-3-methyl-butan-2-ol in A, 1-amino-2-methylpentan-3-ol in B, and 1-amino-2-methylhexan-3-ol in C. QD96 control is coloured in blue, RimM0M1(RimM7 AT)-TR2 + TA1 + FkbS (QD99) is coloured in red. See Supplementary Figure 29 for detailed mass-to-charge (*m/z*) detection.

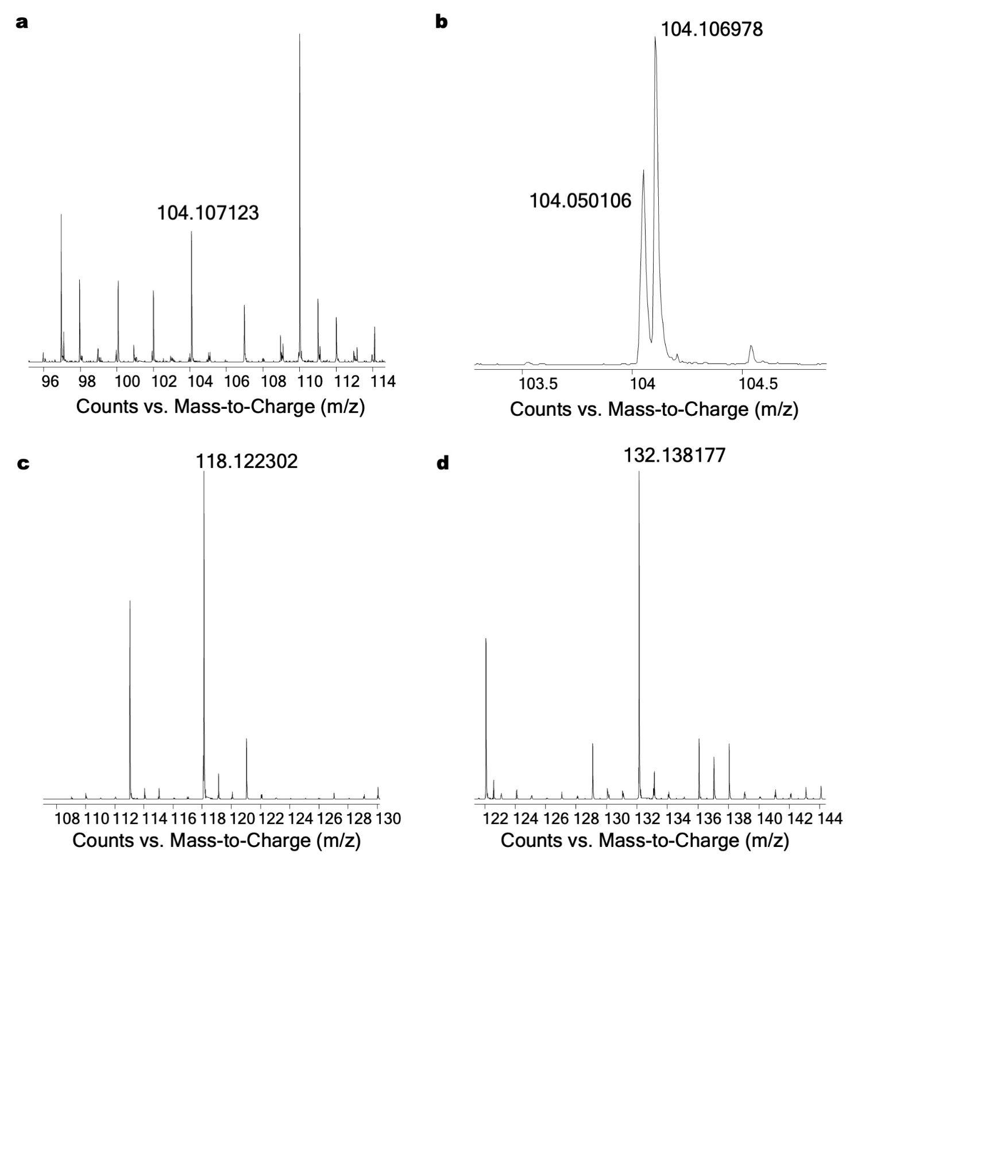

### **Supplementary Figure 29. MS spectra of amino alcohols produced in *S. albus* RimM0M1(RimM7 AT)-TR2 + TA1 + FkbS (QD99).** [M + H]^+^ *m/z* were presented for 4-amino-3-methyl-butan-2-ol standard in A (observed [M + H]^+^ 104.107123 *m/z*), and samples of 4-amino-3-methyl-butan-2-ol in B (observed [M + H]^+^ 104.106978 *m/z*), 1-amino-2-methylpentan-3-ol in C (theoretical [M + H]^+^ 118.122641 *m/z*, observed [M + H]^+^ 118.122302 *m/z*, mass error -2.87 ppm), and 1-amino-2-methylhexan-3-ol in D (theoretical [M + H]^+^ 132.138291 *m/z*, observed [M + H]^+^ 132.138177 *m/z*, mass error -0.86 ppm).

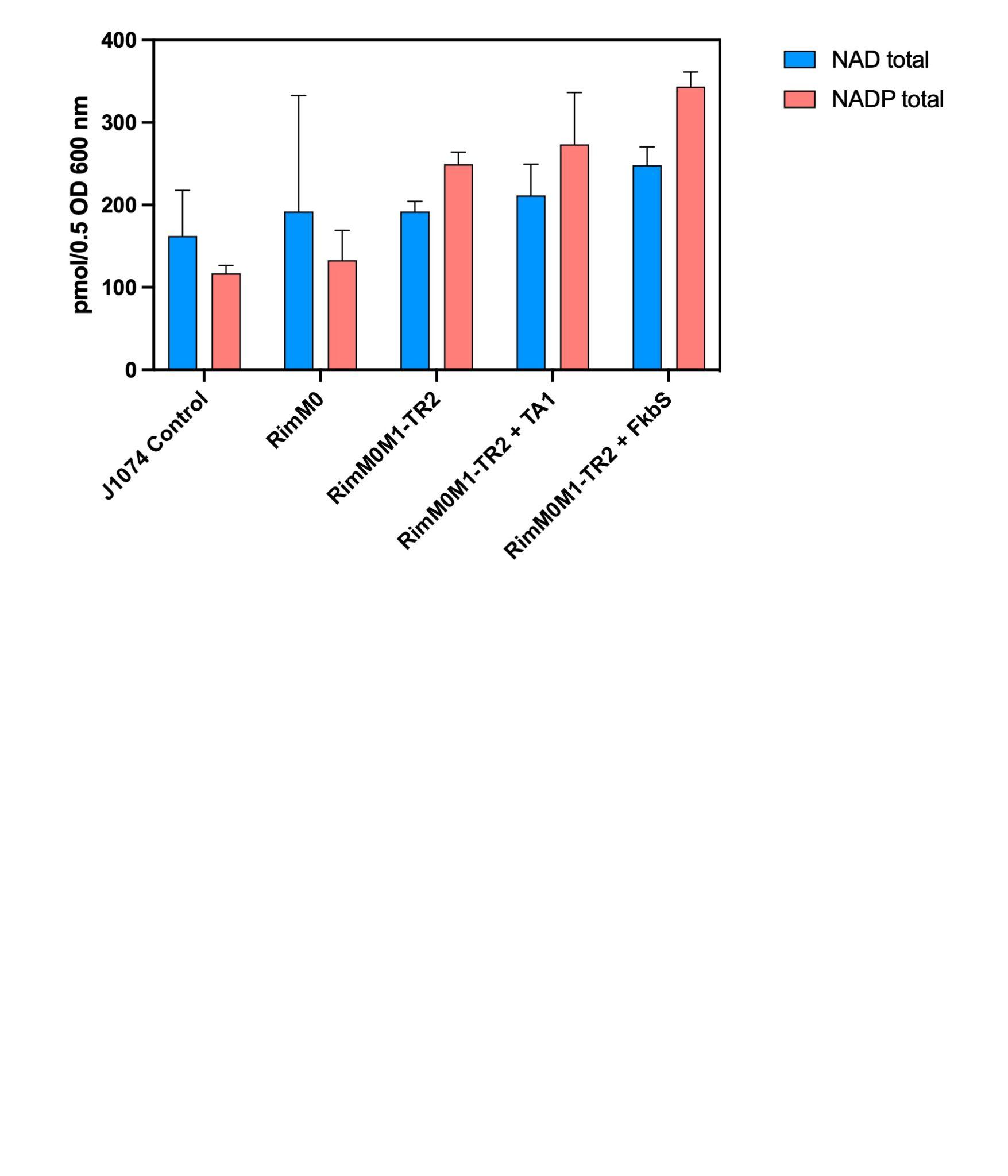

### **Supplementary Figure 30. Intracellular NAD(P) level measurement in *S. albus*.** Each experiment was repeated in quadruplicate except for RimM0M1-TR2 + TA1 which was repeated in duplicate.

### **Supplementary Table 1. Plasmids and strains used in this study**

| **Strains and Plasmids** | **Description** | **Source or Reference** |
| --- | --- | --- |
| **Plasmids** |  |  |
| pHIS | *E. coli* expression plasmid with N-terminal 6✕HIS-TEV tag, kanamycin resistance | Addgene plasmid 29653 |
| pMBP | *E. coli* expression plasmid with N-terminal 6✕HIS-MBP-TEV tag, kanamycin resistance | Addgene plasmid 29656 |
| pG-KJE8 | *E. coli* chaperone coexpression plasmid, chloramphenicol resistance | Takara |
| pHIS_rimA | pHIS plasmid carrying *rimM0* gene | This work |
| pHIS_pimS0 | pHIS plasmid carrying *pimS0* gene | This work |
| pMBP_ACP1 | pMBP plasmid carrying *ACP1* gene | This work |
| pHIS_ACP9 | pHIS plasmid carrying *ACP9* gene | This work |
| pHIS_TR1 | pHIS plasmid carrying *TR1* gene | This work |
| pHIS_TR2 | pHIS plasmid carrying *TR2* gene | This work |
| pHIS_TR7 | pHIS plasmid carrying *TR7* gene | This work |
| pHIS_TR9 | pHIS plasmid carrying *TR9* gene | This work |
| pHIS_TADH2 | pHIS plasmid carrying *TADH2* gene | This work |
| pSC | PhiC31-based *Streptomyces* integration plasmid with spectinomycin resistance | Cruz-Morales *et al.*^12^ |
| p41 | VWB-based *Streptomyces* integration plasmid with apramycin resistance | Cruz-Morales *et al.*^12^ |
| pOSV807 | pSAM2-based *Streptomyces* integration plasmid with hygromycin resistance | Addgene plasmid 126600 |
| pOSV809 | PhiBT1-based *Streptomyces* integration plasmid with kanamycin resistance | Addgene plasmid 126602 |
| pSC_rimA0 | pSC plasmid carrying *PrpsL(RO)-rimM0* | This work |
| pSC_rimA6 | pSC plasmid carrying *Pgapdh(EL)-rimM0* | This work |
| pSC_rimA7 | pSC plasmid carrying *kasOP*-rimM0* | This work |
| p41_rimB18 | p41 plasmid carrying *Pgapdh(EL)-rimM1-DEBS TE* | This work |
| p41_rimB27 | p41 plasmid carrying *Pgapdh(EL)-rimM1-TR1* | This work |
| p41_rimB28 | p41 plasmid carrying *Pgapdh(EL)-rimM1-TR2* | This work |
| p41_rimB31 | p41 plasmid carrying *Pgapdh(EL)-rimM1-TR3* | This work |
| p41_rimB33 | p41 plasmid carrying *Pgapdh(EL)-rimM1-TR4* | This work |
| p41_rimB34 | p41 plasmid carrying *Pgapdh(EL)-rimM1-TR6* | This work |
| p41_rimB35 | p41 plasmid carrying *Pgapdh(EL)-rimM1-TR7* | This work |
| p41_rimB36 | p41 plasmid carrying *Pgapdh(EL)-rimM1-TR8* | This work |
| p41_rimB37 | p41 plasmid carrying *Pgapdh(EL)-rimM1-TR9* | This work |
| p41_rimB40 | p41 plasmid carrying *Pgapdh(EL)-rimM1(rimM13 AT)-TR2* | This work |
| p41_rimB42 | p41 plasmid carrying *Pgapdh(EL)-rimM13(rimM1 docking domain)-TR2* | This work |
| p41_rimB51 | p41 plasmid carrying *Pgapdh(EL)-rimM13(rimM1 docking domain-KS)-TR2* | This work |
| p41_rimB66 | p41 plasmid carrying *Pgapdh(EL)-rimM1(rimM7 AT)-TR2* | This work |
| p41_rimB67 | p41 plasmid carrying *Pgapdh(EL)-rimM1(rimM7 AT)-TR7* | This work |
| p41_rimB69 | p41 plasmid carrying *Pgapdh(EL)-rimM1(pimM7 AT)-TR7* | This work |
| p41_rimB73 | p41 plasmid carrying *Pgapdh(EL)-rimM1(rimM13 AT)-DEBS TE* | This work |
| p41_rimB74 | p41 plasmid carrying *Pgapdh(EL)-rimM1(rimM13 AT)-Pik TE* | This work |
| p41_rimB75 | p41 plasmid carrying *Pgapdh(EL)-rimM1(rimM13 AT)-Rim TE* | This work |
| pOSV807_rimJ | pOSV807 plasmid carrying *Pgapdh(EL)-rimJ* | This work |
| pOSV807_fkbS | pOSV807 plasmid carrying *Pgapdh(EL)-fkbS* | This work |
| pOSV809_TA1 | pOSV809 plasmid carrying *Pgapdh(EL)-TA1* | This work |
| pOSV809_TA2 | pOSV809 plasmid carrying *Pgapdh(EL)-TA2* | This work |
| pOSV809_yqhD | pOSV809 plasmid carrying *Pgapdh(EL)-yqhD* | This work |
| pOSV809_chnD | pOSV809 plasmid carrying *Pgapdh(EL)-chnD* | This work |
| pOSV809_yahK | pOSV809 plasmid carrying *Pgapdh(EL)-yahK* | This work |
| pOSV809_yjgB | pOSV809 plasmid carrying *Pgapdh(EL)-yjgB* | This work |
| pOSV809_ScADH2 | pOSV809 plasmid carrying *Pgapdh(EL)-ScADH2* | This work |
| pOSV809_MaCAR | pOSV809 plasmid carrying *Pgapdh(EL)-MaCAR* | This work |
| pOSV809_yahK_MaCAR | pOSV809 plasmid carrying *Pgapdh(EL)-yahK* and *Pgapdh(EL)-MaCAR* | This work |
| pOSV809_yahK_SrCAR | pOSV809 plasmid carrying *Pgapdh(EL)-yahK* and *Pgapdh(EL)-SrCAR* | This work |
| pOSV809_yahK_MmCAR | pOSV809 plasmid carrying *Pgapdh(EL)-yahK* and *Pgapdh(EL)-MmCAR* | This work |
| pOSV809_yjgB_MaCAR | pOSV809 plasmid carrying *Pgapdh(EL)-yjgB* and *Pgapdh(EL)-MaCAR* | This work |
| pOSV809_yjgB_SrCAR | pOSV809 plasmid carrying *Pgapdh(EL)-yjgB* and *Pgapdh(EL)-SrCAR* | This work |
| pOSV809_yjgB_MmCAR | pOSV809 plasmid carrying *Pgapdh(EL)-yjgB* and *Pgapdh(EL)-MmCAR* | This work |
| **Strains** |  |  |
| *E. coli* BL21(DE3) |  | NEB |
| *E. coli* BAP1 |  | Pfeifer *et al.*^13^ |
| *S. albus* J1074 |  | ATCC |
| *S. coelicolor* M1152 |  | ATCC |
| *S. rimosus* ATCC 10970 |  | ATCC |
| *S. tsukubaensis* NRRL 18488 |  | NRRL |
| *S. natalensis* ATCC 27448 |  | ATCC |
| QD1 | *S. albus* J1074 strain with phiC31-based integration of *PrpsL(RO)-rimM0* | This work |
| QD2 | *S. albus* J1074 strain with phiC31-based integration of *Pgapdh(EL)-rimM0* | This work |
| QD3 | *S. albus* J1074 strain with phiC31-based integration of *kasOP*-rimM0* | This work |
| QD4 | *S. coelicolor* M1152 strain with phiC31-based integration of *PrpsL(RO)-rimM0* | This work |
| QD5 | *S. coelicolor* M1152 strain with phiC31-based integration of *Pgapdh(EL)-rimM0* | This work |
| QD6 | *S. coelicolor* M1152 strain with phiC31-based integration of *kasOP*-rimM0* | This work |
| QD18 | *S. albus* J1074 strain with phiC31-based integration of *PrpsL(RO)-rimM0* and VWB-based integration of *Pgapdh(EL)-rimM1-DEBS TE* | This work |
| QD27 | *S. albus* J1074 strain with phiC31-based integration of *PrpsL(RO)-rimM0* and VWB-based integration of *Pgapdh(EL)-rimM1-TR1* | This work |
| QD28 | *S. albus* J1074 strain with phiC31-based integration of *PrpsL(RO)-rimM0* and VWB-based integration of *Pgapdh(EL)-rimM1-TR2* | This work |
| QD29 | *S. albus* J1074 strain with phiC31-based integration of *Pgapdh(EL)-rimM0* and VWB-based integration of *Pgapdh(EL)-rimM1-TR2* | This work |
| QD30 | *S. albus* J1074 strain with phiC31-based integration of *kasOP*-rimM0* and VWB-based integration of *Pgapdh(EL)-rimM1-TR2* | This work |
| QD31 | *S. albus* J1074 strain with phiC31-based integration of *PrpsL(RO)-rimM0* and VWB-based integration of *Pgapdh(EL)-rimM1-TR3* | This work |
| QD33 | *S. albus* J1074 strain with phiC31-based integration of *PrpsL(RO)-rimM0* and VWB-based integration of *Pgapdh(EL)-rimM1-TR4* | This work |
| QD34 | *S. albus* J1074 strain with phiC31-based integration of *PrpsL(RO)-rimM0* and VWB-based integration of *Pgapdh(EL)-rimM1-TR6* | This work |
| QD35 | *S. albus* J1074 strain with phiC31-based integration of *PrpsL(RO)-rimM0* and VWB-based integration of *Pgapdh(EL)-rimM1-TR7* | This work |
| QD36 | *S. albus* J1074 strain with phiC31-based integration of *PrpsL(RO)-rimM0* and VWB-based integration of *Pgapdh(EL)-rimM1-TR8* | This work |
| QD37 | *S. albus* J1074 strain with phiC31-based integration of *PrpsL(RO)-rimM0* and VWB-based integration of *Pgapdh(EL)-rimM1-TR9* | This work |
| QD40 | *S. albus* J1074 strain with phiC31-based integration of *PrpsL(RO)-rimM0* and VWB-based integration of *Pgapdh(EL)-rimM1(rimM13 AT)-TR2* | This work |
| QD42 | *S. albus* J1074 strain with phiC31-based integration of *PrpsL(RO)-rimM0* and VWB-based integration of *Pgapdh(EL)-rimM13(rimM1 docking domain)-TR2* | This work |
| QD51 | *S. albus* J1074 strain with phiC31-based integration of *PrpsL(RO)-rimM0* and VWB-based integration of *Pgapdh(EL)-rimM13(rimM1 docking domain-KS)-TR2* | This work |
| QD66 | *S. albus* J1074 strain with phiC31-based integration of *PrpsL(RO)-rimM0* and VWB-based integration of *Pgapdh(EL)-rimM1(rimM7 AT)-TR2* | This work |
| QD67 | *S. albus* J1074 strain with phiC31-based integration of *PrpsL(RO)-rimM0* and VWB-based integration of *Pgapdh(EL)-rimM1(rimM7 AT)-TR7* | This work |
| QD69 | *S. albus* J1074 strain with phiC31-based integration of *PrpsL(RO)-rimM0* and VWB-based integration of *Pgapdh(EL)-rimM1(pimM7 AT)-TR7* | This work |
| QD73 | *S. albus* J1074 strain with phiC31-based integration of *PrpsL(RO)-rimM0* and VWB-based integration of *Pgapdh(EL)-rimM1(rimM13 AT)-DEBS TE* | This work |
| QD74 | *S. albus* J1074 strain with phiC31-based integration of *PrpsL(RO)-rimM0* and VWB-based integration of *Pgapdh(EL)-rimM1(rimM13 AT)-Pik TE* | This work |
| QD75 | *S. albus* J1074 strain with phiC31-based integration of *PrpsL(RO)-rimM0* and VWB-based integration of *Pgapdh(EL)-rimM1(rimM13 AT)-Rim TE* | This work |
| QD76 | *S. albus* J1074 strain with phiC31-based integration of *PrpsL(RO)-rimM0*, VWB-based integration of *Pgapdh(EL)-rimM1-TR2*, and pSAM2-based integration of *Pgapdh(EL)-fkbS* | This work |
| QD77 | *S. albus* J1074 strain with phiC31-based integration of *PrpsL(RO)-rimM0*, VWB-based integration of *Pgapdh(EL)-rimM1-TR2*, and pSAM2-based integration of *Pgapdh(EL)-rimJ* | This work |
| QD78 | *S. albus* J1074 strain with phiC31-based integration of *PrpsL(RO)-rimM0*, VWB-based integration of *Pgapdh(EL)-rimM1-TR2*, pSAM2-based integration of *Pgapdh(EL)-fkbS*, and phiBT1-based integration of *Pgapdh(EL)-yqhD* | This work |
| QD79 | *S. albus* J1074 strain with phiC31-based integration of *PrpsL(RO)-rimM0*, VWB-based integration of *Pgapdh(EL)-rimM1-TR2*, pSAM2-based integration of *Pgapdh(EL)-fkbS*, and phiBT1-based integration of *Pgapdh(EL)-chnD* | This work |
| QD80 | *S. albus* J1074 strain with phiC31-based integration of *PrpsL(RO)-rimM0*, VWB-based integration of *Pgapdh(EL)-rimM1-TR2*, pSAM2-based integration of *Pgapdh(EL)-fkbS*, and phiBT1-based integration of *Pgapdh(EL)-yahK* | This work |
| QD81 | *S. albus* J1074 strain with phiC31-based integration of *PrpsL(RO)-rimM0*, VWB-based integration of *Pgapdh(EL)-rimM1-TR2*, pSAM2-based integration of *Pgapdh(EL)-fkbS*, and phiBT1-based integration of *Pgapdh(EL)-yjgB* | This work |
| QD82 | *S. albus* J1074 strain with phiC31-based integration of *PrpsL(RO)-rimM0*, VWB-based integration of *Pgapdh(EL)-rimM1-TR2*, pSAM2-based integration of *Pgapdh(EL)-fkbS*, and phiBT1-based integration of *Pgapdh(EL)-ScADH2* | This work |
| QD83 | *S. albus* J1074 strain with phiC31-based integration of *PrpsL(RO)-rimM0*, VWB-based integration of *Pgapdh(EL)-rimM1(rimM7 AT)-TR2*, and pSAM2-based integration of *Pgapdh(EL)-fkbS* | This work |
| QD84 | *S. albus* J1074 strain with phiC31-based integration of *PrpsL(RO)-rimM0*, VWB-based integration of *Pgapdh(EL)-rimM1(rimM7 AT)-TR7*, and pSAM2-based integration of *Pgapdh(EL)-fkbS* | This work |
| QD85 | *S. albus* J1074 strain with phiC31-based integration of *PrpsL(RO)-rimM0*, VWB-based integration of *Pgapdh(EL)-rimM1(pimM7 AT)-TR7*, and pSAM2-based integration of *Pgapdh(EL)-fkbS* | This work |
| QD86 | *S. albus* J1074 strain with phiC31-based integration of *PrpsL(RO)-rimM0*, VWB-based integration of *Pgapdh(EL)-rimM1(rimM13 AT)-TR2*, and pSAM2-based integration of *Pgapdh(EL)-fkbS* | This work |
| QD87 | *S. albus* J1074 strain with phiC31-based integration of *PrpsL(RO)-rimM0*, VWB-based integration of *Pgapdh(EL)-rimM1(rimM13 AT)-TR2*, and phiBT1-based integration of *Pgapdh(EL)-MaCAR* | This work |
| QD88 | *S. albus* J1074 strain with phiC31-based integration of *PrpsL(RO)-rimM0*, VWB-based integration of *Pgapdh(EL)-rimM1(rimM13 AT)-TR2*, pSAM2-based integration of *Pgapdh(EL)-fkbS*, and phiBT1-based integration of *Pgapdh(EL)-yahK* and *Pgapdh(EL)-MaCAR* | This work |
| QD89 | *S. albus* J1074 strain with phiC31-based integration of *PrpsL(RO)-rimM0*, VWB-based integration of *Pgapdh(EL)-rimM1(rimM13 AT)-TR2*, pSAM2-based integration of *Pgapdh(EL)-fkbS*, and phiBT1-based integration of *Pgapdh(EL)-yahK* and *Pgapdh(EL)-SrCAR* | This work |
| QD90 | *S. albus* J1074 strain with phiC31-based integration of *PrpsL(RO)-rimM0*, VWB-based integration of *Pgapdh(EL)-rimM1(rimM13 AT)-TR2*, pSAM2-based integration of *Pgapdh(EL)-fkbS*, and phiBT1-based integration of *Pgapdh(EL)-yahK* and *Pgapdh(EL)-MmCAR* | This work |
| QD91 | *S. albus* J1074 strain with phiC31-based integration of *PrpsL(RO)-rimM0*, VWB-based integration of *Pgapdh(EL)-rimM1(rimM13 AT)-TR2*, pSAM2-based integration of *Pgapdh(EL)-fkbS*, and phiBT1-based integration of *Pgapdh(EL)-yjgB* and *Pgapdh(EL)-MaCAR* | This work |
| QD92 | *S. albus* J1074 strain with phiC31-based integration of *PrpsL(RO)-rimM0*, VWB-based integration of *Pgapdh(EL)-rimM1(rimM13 AT)-TR2*, pSAM2-based integration of *Pgapdh(EL)-fkbS*, and phiBT1-based integration of *Pgapdh(EL)-yjgB* and *Pgapdh(EL)-SrCAR* | This work |
| QD93 | *S. albus* J1074 strain with phiC31-based integration of *PrpsL(RO)-rimM0*, VWB-based integration of *Pgapdh(EL)-rimM1(rimM13 AT)-TR2*, pSAM2-based integration of *Pgapdh(EL)-fkbS*, and phiBT1-based integration of *Pgapdh(EL)-yjgB* and *Pgapdh(EL)-MmCAR* | This work |
| QD94 | *S. albus* J1074 strain with phiC31-based integration of *PrpsL(RO)-rimM0*, VWB-based integration of *Pgapdh(EL)-rimM1-TR1*, and phiBT1-based integration of *Pgapdh(EL)-TA1* | This work |
| QD95 | *S. albus* J1074 strain with phiC31-based integration of *PrpsL(RO)-rimM0*, VWB-based integration of *Pgapdh(EL)-rimM1-TR1*, and phiBT1-based integration of *Pgapdh(EL)-TA2* | This work |
| QD96 | *S. albus* J1074 strain with phiC31-based integration of *PrpsL(RO)-rimM0*, VWB-based integration of *Pgapdh(EL)-rimM1-TR2*, and phiBT1-based integration of *Pgapdh(EL)-TA1* | This work |
| QD97 | *S. albus* J1074 strain with phiC31-based integration of *PrpsL(RO)-rimM0*, VWB-based integration of *Pgapdh(EL)-rimM1-TR2*, and phiBT1-based integration of *Pgapdh(EL)-TA2* | This work |
| QD98 | *S. albus* J1074 strain with phiC31-based integration of *PrpsL(RO)-rimM0*, VWB-based integration of *Pgapdh(EL)-rimM1-TR2*, pSAM2-based integration of *Pgapdh(EL)-fkbS*, and phiBT1-based integration of *Pgapdh(EL)-TA1* | This work |
| QD99 | *S. albus* J1074 strain with phiC31-based integration of *PrpsL(RO)-rimM0*, VWB-based integration of *Pgapdh(EL)-rimM1(rimM7 AT)-TR2*, pSAM2-based integration of *Pgapdh(EL)-fkbS*, and phiBT1-based integration of *Pgapdh(EL)-TA1* | This work |

### **Supplementary Table 2. Primers used in this study**

| **Primer name** | **Sequence (5' to 3')** |
| --- | --- |
| pHIS_rimA-F | TACTTCCAATCCAATGCAatggtcccggtgcatactg |
| pHIS_rimA-R | TTATCCACTTCCAATGTTATCAATGTTCTGCATCCATAAGACGG |
| pHIS_pimS0-F | TACTTCCAATCCAATGCAatggtgcccgtccacacagatgactacgc |
| pHIS_pimS0-R | TTATCCACTTCCAATGTTAtcagctctccgcgtccatcagccg |
| pMBP_ACP1-F | TACTTCCAATCCAATGCAgctgggctaggagatgcag |
| pMBP_ACP1-R | TTATCCACTTCCAATGTTAtacttcaccatgcaggtactccg |
| pHIS_ACP9-F | TACTTCCAATCCAATGCAggaggcatgaccggcgcg |
| pHIS_ACP9-R | TTATCCACTTCCAATGTTActccagcaggaccgtccggagg |
| pHIS_TR1-F | TACTTCCAATCCAATGCAcgggcggaactgtcggcgg |
| pHIS_TR1-R | TTATCCACTTCCAATGTTAtcacccggccgccggg |
| pHIS_TR2-F | TACTTCCAATCCAATGCAaccggggccagcggct |
| pHIS_TR2-R | TTATCCACTTCCAATGTTActagcgggccggggccg |
| pHIS_TR7-F | TACTTCCAATCCAATGCAggcggacggacgcagggc |
| pHIS_TR7-R | TTATCCACTTCCAATGTTAtcagcctgccgcggggaag |
| pHIS_TR9-F | TACTTCCAATCCAATGCAggcaccacgggcagcggc |
| pHIS_TR9-R | TTATCCACTTCCAATGTTAtccttcggtcatgggggccgcc |
| pHIS_TADH2-F | TACTTCCAATCCAATGCAatgtctactacagtcggccaag |
| pHIS_TADH2-R | TTATCCACTTCCAATGTTAtcagtcagccatagtaatgatacatctc |
| pSC_rimA0-0F | gacgcggagcactgataactagcataaccccttggggc |
| pSC_rimA0-0R | gtggacgggcaccatttaccggctttcttcgtgttgaac |
| pSC_rimA0-1F | atggtgcccgtccacacagc |
| pSC_rimA0-1R | tcagtgctccgcgtccatcag |
| p41_rimB27-0F | cgcgatatcgaattcctcgag |
| p41_rimB27-0R | cttctcctcgttcgacatatggcgtatcccctttcag |
| p41_rimB27-1F | atgtcgaacgaggagaagctgcg |
| p41_rimB27-1R | gacctcggagcgcagatacg |
| p41_rimB27-2F | tgcgctccgaggtccgggcggaactgtcg |
| p41_rimB27-2R | cgaggaattcgatatcgcgttctttccgattcacccggc |
| p41_rimB28-2F | tgcgctccgaggtcaccggggccagcggct |
| p41_rimB28-2R | cgaggaattcgatatcgcgctagcgggccggggcc |
| p41_rimB31-2F | tgcgctccgaggtcgggtccgccgacccccg |
| p41_rimB31-2R | cgaggaattcgatatcgcgttacgcctcgacgctgaccgcg |
| p41_rimB33-2F | ctgcgctccgaggtcggcgcaccgggcggc |
| p41_rimB33-2R | gctagttactcgaggaattcgtcacggggcgggcaggtaacc |
| p41_rimB34-2F | ctgcgctccgaggtcggcgggccggacggc |
| p41_rimB34-2R | gctagttactcgaggaattcgtcaggcagcgggccgcg |
| p41_rimB35-2F | ctgcgctccgaggtcggcggacggacgcagggc |
| p41_rimB35-2R | gctagttactcgaggaattcgtcagcctgccgcggggaagt |
| p41_rimB36-2F | CTGCGCTCCGAGGTCGCGCCCCAGGACGACACCGGG |
| p41_rimB36-2R | GCTAGTTACTCGAGGAATTCGTCACGGGGCGGGCGGGAAGTG |
| p41_rimB37-2F | ctgcgctccgaggtcggcaccacgggcagcggc |
| p41_rimB37-2R | gctagttactcgaggaattcgtcatccttcggtcatgggggccgc |
| p41_RimB40-0F | cgtcgacctgcccacgtacgcc |
| p41_RimB40-0R | ggtgcctgctccaggatgacgtgg |
| p41_RimB40-1F | cctggagcaggcaccggcggacgcc |
| p41_RimB40-1R | gtgggcaggtcgacgcggcg |
| p41_RimB42-0F | ctgcgcagcgagttcaccggggccagcggctcc |
| p41_RimB42-0R | ggctcgcgctgcttcgcctcg |
| p41_RimB42-1F | aagcagcgcgagccggtcgtcatcgtcggcatggc |
| p41_RimB42-1R | gaactcgctgcgcagccgggc |
| p41_RimB51-0F | aaaggccaggaaccgtaaaaaggccgc |
| p41_RimB51-0R | cttctcctcgttcgacatatggcgtatcccc |
| p41_RimB51-1F | atgtcgaacgaggagaagctgcgcga |
| p41_RimB51-1R | ggtgcctgctccaggatgacgtgg |
| p41_RimB51-2F | cctggagcaggcaccggcggacgcc |
| p41_RimB51-2R | cggttcctggccttttgctggccttt |
| p41_RimB69-0F | cgctgcccacgtacgccttccagcg |
| p41_RimB69-0R | tcagccaatcgactggcgagcggcat |
| p41_RimB69-1F | ccagtcgattggctgagctcatgagcgg |
| p41_RimB69-1R | gcggggggcctcctccaggatgacgtgggcgttggtg |
| p41_RimB69-2F | gaggccccccgcccgcacga |
| p41_RimB69-2R | gtacgtgggcagcggcacccgg |
| p41_RimB73-0F | tagcgcgatatcgaattcctcgagtaactagcataacccc |
| p41_RimB73-0R | gacctcggagcgcagatacgcggc |
| p41_RimB73-1F | tgcgctccgaggtcgactccggtacgcctgcac |
| p41_RimB73-1R | gaattcgatatcgcgctatcagctgttgccgccac |
| p41_RimB74-0F | tagcgcgatatcgaattcctcgagtaactagcataacccc |
| p41_RimB74-0R | gacctcggagcgcagatacgcggc |
| p41_RimB74-1F | tgcgctccgaggtcgccgagcggaactgggccg |
| p41_RimB74-1R | gaattcgatatcgcgctacttgcccgccccctcgatgc |
| p41_RimB75-0F | tagcgcgatatcgaattcctcgagtaactagcataacccc |
| p41_RimB75-0R | gggctgtgcggcgacctcggagcgcagatacgcggc |
| p41_RimB75-1F | gccgcacagcccggcct |
| p41_RimB75-1R | gaattcgatatcgcgctatcatgcctggccgagggtggcc |
| pSC_RimA6-0F | ctagcataaccccttggggcc |
| pSC_RimA6-0R | gaccgaaggagcagcagtactctgctgcaacgtgtg |
| pSC_RimA6-1F | gctgctccttcggtcgga |
| pSC_RimA6-1R | atggcgtatcccctttcagatact |
| pSC_RimA6-2F | gaaaggggatacgccatatggtgcccgtccacaca |
| pSC_RimA6-2R | cccaaggggttatgctagttatcagt |
| pSC_RimA7-0F | ctagcataaccccttggggcc |
| pSC_RimA7-0R | gttcgaatgtgaacaagtactagtactctgctgcaacgtgtg |
| pSC_RimA7-1 (direct use) | agtacttgttcacattcgaacggtctctgctttgacaacatgctgtgcggtgttgtaaagtcgtggccaggagaatacgacagcgtgcaggactgggggagttcat |
| pSC_RimA7-2F | ggactgggggagttcatatggtgcccgtccacaca |
| pSC_RimA7-2R | cccaaggggttatgctagttatcagt |
| p41_RimB66-0F | ccggcgaggagctggccgccctgccgcccg |
| p41_RimB66-0R | aacacaccggtctgcggtgggg |
| p41_RimB66-1F | gcagaccggtgtgttcaaggaagacgacg |
| p41_RimB66-1R | cagccagtaccactggcgctggaagg |
| p41_RimB66-2F | ccagtggtactggctgcaggaaactccccacgcggcacca |
| p41_RimB66-2R | cagctcctcgccggacagcgg |
| p41_RimB67-0F | gccgccctgccgcccg |
| p41_RimB67-0R | aacacaccggtctgcggtgggg |
| p41_RimB67-2F | ccagtggtactggctgcccgccggc |
| p41_RimB67-2R | cggcagggcggcccgggccgcccagtcggc |
| pOSV807_rimJ-0F | GACGCTGGAGGACTGActagcataaccccttggggc |
| pOSV807_rimJ-0R | AGCGTTCAGTATTTCCTTCATgcgtatcccctttcagatactcg |
| pOSV807_rimJ-1F | ATGAAGGAAATACTGAACGCTATTCAGGC |
| pOSV807_rimJ-1R | TCAGTCCTCCAGCGTCCC |
| pOSV807_fkbS-0F | GGTTCCGCAATGTCTGActagcataaccccttggggc |
| pOSV807_fkbS-0R | CGTCCAGGATTTCCTTCATgcgtatcccctttcagatactcg |
| pOSV807_fkbS-1F | TGAAGGAAATCCTGGACGCGAT |
| pOSV807_fkbS-1R | TCAGACATTGCGGAACCGGT |
| pOSV809_TA1-0F | actagtagcggccgctt |
| pOSV809_TA1-0R | aaaaaacgctagcagcggc |
| pOSV809_TA1-1F | cgctgctagcgttttttgctgctccttcggtcgg |
| pOSV809_TA1-1R | gcggccgctactagtgctgtgcttcctgttcggcttaa |
| pOSV809_TA2-0F | actagtagcggccgctt |
| pOSV809_TA2-0R | aaaaaacgctagcagcggc |
| pOSV809_TA2-1F | cgctgctagcgttttttgctgctccttcggtcgg |
| pOSV809_TA2-1R | gcggccgctactagtgctgtgcttcctgttcggcttaa |
| pOSV809_MaCAR-0F | ctagcataaccccttggggcc |
| pOSV809_MaCAR-0R | gcgtatcccctttcagatactcg |
| pOSV809_MaCAR-1F | atctgaaaggggatacgccatatatgacggagacgatctccaccgccg |
| pOSV809_MaCAR-1R | ctcggtcatctccggggacagcggcgccgacc |
| pOSV809_MaCAR-2F | ccggagatgaccgagttcatggagagtctgc |
| pOSV809_MaCAR-2R | agccgctcgagccagtccagcacgagccagcggc |
| pOSV809_MaCAR-3F | tggctcgagcggctggcg |
| pOSV809_MaCAR-3R | ccaaggggttatgctagttagacgagacccaggagctggatgtcggag |
| pOSV809_ADH_CAR-0F | aacagggaagtgagagggcc |
| pOSV809_ADH_CAR-0R | accttgccgaaggcaattacat |
| pOSV809_ADH_CAR-1F | gccttcggcaaggtcgc |
| pOSV809_ADH_CAR-1R | aagtatcttcctggcatcttccag |
| pOSV809_ADH_CAR-2F | cctctcacttccctgttgctgctccttcggtcgg |
| pOSV809_ADH_CAR-2R | gatgccaggaagatacttcaaaaaacccctcaagacccgtt |

### **Supplementary Table 3. Genes used in this study**

| **Gene name** | **Sequence (5' to 3')** |
| --- | --- |
| *rimA (rimM0)* | atggtgcccgtccacacagctgactacgtgatccagccgcccaccgacgcggcggaggtacggggcggccacacgctgcccgaggtgttcgagaccgcgtcgagggccgcccccgacgcggtggccatcgtggacggggaccgttcgcggacctgggcgcagtggcgggcggacgtccgcgcgctggcccgtggcctccaggaatcgggcgtcgggcccggtgacgtggtggcggtgcggctgccgaacagctgggagttccagaccctgcacctggccgtcgcggccgtcggcgccgtactgctgcccgtccacgagggcaccccgaccgcggaggtccacgcgctgctgacccgcgcggaacccgtactcctcgtcctgtcggcttccgggagcgagggcacggcaacggcccgctcgctcctggagagcgtgccgacgctgcgcggcgtgctgctggccggggcgtcggaggcggagtgcgaggagccggggatcggggcgctggacgggttgcgggcagcctggacgggcagcgagccccggccggtgcacctcacccccgacatgccgctcgccctgatcccctcgtccggcaccacctcggcgcggcccaagctgtgcgtgcacacccacggcggcctgctggcgaacaccgcggccgtcgtggccgatgccgccgacgccttcggcggcacggtcctgaccgcctgcccgatgacccacctgttcgggctccagtccctgcacacggcgctgttcgccgcccggacgcaggtcctgctcaccggctgggacgtggaccgcttcctggagctggcgcgggagcacaacccccgcgtcgtcttcgccgtacccgcccaactgcgggacatcgtcgcgcgactcgccagggcgggcgagccggcgggcttcgcgccgcaccaggtgcgtacggcgggcgccgccctcgcacccgcgctcgccgcacagatacgcgccgccctcgactgcgaaccggtcgtggtgtggggcatgtccgagatcggcaccggcaccgcaacccgggcccaccaccccgacggcagcgtgggagaaccggtcgacggcgtgcgcgtacgggtcgtcgacgagcacggcgcggagtgcgcggcgggggagacgggcgagctccagtaccagggcccggcgatgttccgcggctatttccgcgagccggagctgacgcgctcggccctcaccgacgacggctggctgcggaccggcgacatcgccgccatcgacgcggacggcgtggtcgtcctgcacggccgggcggccgaagtgatcgccaccggaggccggaagttcggcgccaccgaaatcgagagcctgctcgcggagctcgcgggcctgggcccgctggccgtggcgggggcgccggacgaccggctcggcgagtacccgtgcctggtcgtgaccgaccgcgcggaccgtacgatcgggctgaccgaggtgaccgcgttcctgcgccggcggggactcgccgaccacaagatccccctcgaactggtcaccgtgcgcgagctgcccttcacccccgcggggaagctcgaccgcagggcactcaaggagcagctgcgcaccggcctggaagcgacgcccgtggcggcccgtctcggcgcggtcccgccggagacggccgaagaggcgctggaactggtacgcgactgcgtcggccaggtcctcggcagcagcggcgacccggccggccccgccgtcggtgcgccggaattccccgacaccgaattccgccggctcggcctcgactccgtcctcgcggtgcggctgcgcaacctgctgcgggaggagaccgggctgtccctcccggtcacgctggccttcgacttcccgacgccgcgggccgtggcgcacgcactggccgagcagaacgacccggcttcggcggcggcatcggggaagacggcggaggaggcgcggaggttcccggccgacgaggccgatccggtggcgatcgtcgccatggcctgccgcctgccgggcggcgccgactctcccgacgccctgtgggagctgctggccggcggaaccgacgcgatgcggcccttcccggacgaccgcggctgggacctggaccgcctcttcgacgaggaccccgaccggccgggcacctgctacgcacgcgaaggcggcttcctgcccggcgcgggcgacttcgacgcgggcttcttcggcctgtccgaccacgaggcgacggccaccgacccgcagcagcgcctgctcctggtggcggcctgggagaccttcgagcgggcgggcatcgacccggcgtccctgaagggcacccgtacgggcgtgttcaccggcgcgatggagcgcggctacggcgctctcgcgtccgccgtccccagcgagtgggagagcacgctcaccaccggggccgccaacagcgcgatctccgggcgcatcgcctacacctacggcctcgaaggccccgcgctgaccgtcgacaccgcctcctcgtcctccctcgtcgccctgcacctggcctgccgctccctgcggtcgggcgagaccgacctggcgctggccggcggcgtcaccgtcatggcgaccccggcgcccttcacccacttcgcccggctgcgcgcgctcgccaccgactcgcgcgccaaggcgtacgcggacaccgcgaacggctccgcgtgggcggaaggcgcgggactgctgctgctggagcggctgagcgacgcccgccgcaacggacaccgtgtactggcgctcgtacggggctccgcggtgaaccaggacggcgcctccaacgggctcaccgccccgagcggccccgcgcagcagcgcgtcatccgccaggccctggccgacgccggtctgacgccgcgggacgtggacgcggtggaggggcacggcaccggcactccgctcggcgacccgatcgaggcccaggccctgatggccacctacggccaggagcggccggagggacggccgctgtggctggggtcggtgaagtccaacctcgggcacacacaggccgctgccggggtcgtcggggtcatcaagacggtgctcgcgctgggccgcggcgtcctgcccaggacgctgcacgtggagactccttccaccaaggtcgactggtccgccggggcggtacggctgctgaccgaggcgcagccctggccccaggagagcgggcacacgcggcgggcaggagtgtcgtccttcgggctcaccggcaccaacgcccacgtgatcctggaggaggccccggacgggacgcagagcgcgcccggatcggaaccggccgacgacaccgccgtaccgtgggtgctctccgcacggagccggacggcgctgcgcgaacaggcccgccgcctggccgagcacgtgaccgctcaccccggcctgcgcacacaggacatcgcccacgccctggccaccacccgcacccggcaccggcaccgggccgtcgtcagcggctccgaccgggaccggatgctgtccgcgacggccgcgttcgggcgcggggagcgggccgcggacgtcaccccgctcgattccgcgccgggcggcctggccttcgtcttctccggacagggcggccagcaccccggcatggggcgcggggcggccgaggcgttccccgtgttcggggaggcgctgcgcgaggtgtgcgacaccctcgacccgctgctggcacgtccgctgacctcggtgatgtgggcggacgccgactccgaggaggcgacgctgctgcacaacgccgagttctcgcagccctcgctgttcgccctccaggtcgccctctaccggctgtacgagtcctggggcatggccccggaccgtctcgcgggccattcggccggcgagatcgccgccgcgcatgtcaccggcatcctcaccctccaggacgcctgcgccctggtggcctcccggggcaggctgatcagctcgctgccggtcggcggcgcgacggtggcggtgcgcatctcggaggacgaggtgcgggggtggctcgccgaggagacgaccggctcggtctcgatcgcggccgtcaacgggccgcactccctcgtactctccggtgccgaggccccgctcatcgccctcacggaccggctccgcgacgccggccacaagacccaccgcatccccatgagggtcgcggcccactcaccgctgatggaccccatcctgggggagttccgcgcggtcgtccgcacgctggcctacggcacgcccaccatccccctcgtctccaccgtcaccggccgcccgctgaccgacgaggaggcgcgcgaccctgagcactgggtacggcacgtgcggcagcccgtgcgcttcaaggacgcgatcggccggctccgggaggagcgcgtcaccggcttcctggagctgggcgccgagccgctgctcacccccatgatcgacgagtgcctggaggcggccggcccgcagcacggaaccgccgtggtgccgagcctgagctccggcgtaccggaccggcagatcctgctctccgcggccgcccgggtgcacacccacggcgcacccgtcgactgggacgcggtgctgcccggggcccggcccgtcgacctgccgacgtacgcgttccagcggcggcggttctggctggcggcggggccgggtgcggccgccgaggccggtttcgtgggtgccgcgatcggcgcgggcgcgccggcggccgatgcggaggagccgtcgggcctggaagcccggctggccggtctcgaagacgccgaacaggacgcgtacgtacgctccctggtgctcgccgagacctcggccgtgctcggcggccaggaaccgctcgacagcgaaggcacccacaccttcaaggagatgggcatcaactcggtgaacgccgtcgagctccgcaaccgcctcatcgcggccacggacctgcggctccccgccacgctcgtctacgaccaccccaccccgaacgcggtcgtccgcctcgtacgcgaacgcctcgcgcgcccggccaccgccgcacgcgacgtggactcggtcgtggccgagctggagtccctgctcacggccggcgcggaggtctcggcggaggcggtggcacggctgaaggcgatggcggcggggcgggacggcactacggacaccggttccggcgggcccctggacctgacgtcggccagcgacgaggacctgttccggctgatggacgcggagcactga |
| *pimS0* | atggtgcccgtccacacagatgactacgcgatccagccgcccgccgacaccgcgcatggagggggtggcttcacactgcctgctgtgttcgaggccgccgtggaatccgcccccgacgcagtggccctcgtcgacgggcaccgttcctggacctgggcgcagtggcgcgcagacgtcgatgcgctggcccgcgggttgcaggagtcgggtatcgcacccggtgacgtggtggcggtgcggctgccgaactgcgggaggtttcccaccctgcacctggccgtcgcggccgtcggagcggtactgctgcccatccaccagggcaccccgctcccggaggtcgacgcgctgctgacccgggcggaacccgcccttctcgtcctgtcggccgccgggagcgacggcctggcgacggcccgttcgcttctggagagcgtgccgtcgctgcgcggcgtgctgctggccggggcgtcggagacggggagtcgggatccgtcggaggtggggagtcaggatccgggcgtcgggtcgctggacgggctgctggcggcctgggcgggcagcgggccccggccggtggacgtcacccccgatatgccgctcgttctcgtcccctcgtccgggacggtctcggcacggcccaaactgtgtgtgcacagccacgacggtctgctgtcgaacaccgcagccgtcaccgccgaggccgccgacgccttcgacggtccggtcctcacggcctgcccgatgacccatctgttcggcctccagtccctgcacgcggcgctgttcgccgcctgcacgcaggtcctgctcaccggctgggacgtggaccgcttcctggaacaggctcgggagcacggcccccgtgtcgtgttcgccgtgcccgcccaactgcgggacgtcgtgacacgactggccaggaccggcgaaccggcgggcttcacgccgtaccaggtgcgtaccgcgggggccgccgtcgcgcccgcgctcgccgtacgggtacgcgccgtcctcgactgcgaactggtcgtggtgtgggggatgtccgagatcggcaccggcacccgcacccgggcccaccaccccgacggctgtgtgggagagccggtcagcggggtggacgtgcgcgtcgtcgacgagcacggccaagagtgcgcggcggacgagaggggtgaactccagtaccgggggccagggttgttccgcggctatttccgcgaaccggagctgacgcgctcggccctcaccgacgacggctggctgcggaccggcgatctcgcgaccgtcgacgcggacggcgtggtggtgctgcacggccgggcggccgagctgatcaatacggggggccggaagttcagcgccggcgaggtcgagggactgctctccggctttacggacctgggcccgctggccgtcgtcggcgcaccggacgaccggctcggggagtacccgtgcctggtcgtgaccgaccacgcggacggcaccatcggcctgagcgaggtgaccgcgttcctgcgccggctggggctcgccgaccacaagatccccctcgaactggtcaccgtgcgcgagcttcccttctcccctgccgggaagctcgaccgcggggcgctcaagcggctgctcgccaaccttgcggaggtgtccgttccggcccgtctcggcgcggtcccgccctacacggccgaggaggcgctggacctcgtacgcgactgcgtcggccgggtgctccgttacggcggcgcggccgtccccttccccccggacaaggacttcttctccccggacaaggacttccgccagctcggcctggactccatcggcgcggtgcggctgcggaatctgctgcgggaggagaccgggctgccgctcccggccaccctggccttcgactcccccactccgcgagccgtcgcgcgcgtcctggccgagcaggaggagccgtcccaggacgagccgagggagaacccggcggacggtgccgacccggtggcgatcgtgggcatggcctgccggctgccgggcggagccgactcccccgacgccctgtgggagctgctcgccgacgggaccgacgcgatgtcccccttccccacggaccgcggctgggacctggaccggctgttcgacgaggatgccgaccgcccgggtacctcgtacgcccgcgaaggcggcttcctgcacgacgcgggcgacttcgacgcgggcttcttcggcctgtcggaccaggaggcgacggcgaccgatccgcagcagcggctgcttctggaggcggcctgggagaccttcgagcgggcgggcatcgacccgcagtccctgaggggaagccgtacgggcgtgttcacgggcgcgatggaccgcggctatggaaccagcgcgtccgccgcgcccagcgcatgggagagcatgctcatcaccgggaccgccggcagcgcggtctcggggcgcatcgcctacacctacgggctcgaaggccccgcgctgacggtcgacaccgcctcctcgtcctccctcgtcgccctgcatctggcctgccggtcgctgcgctcgggcgagaccgacctggcgctggccggcggcgtcaccgtcatggcgaccccggcgcccttcgcacacttctcccggctgcgcgcgctgtcccccgactcccgctccatggcgtacgcggacgccgcgaacggctcggcgtggtcggagggcgcggggctgcttctgctggagcggctgagcgacgcccggcgcaacggacaccgtgtcctggcgctcgtacggggctccgccgtgaatcaggacggcgcctccaacgggctcaccgcgccgagcggacccgcacagcagcgcgtcatccgccaggccctggccgacgccgggctgacgccgcaggacgtggacgccgtggaggggcacggcaccggcacgccgctcggcgaccccatcgaggcgcaggcgctgctggccacgtacggccaacagcggcctgtggaacggccgttgtggctggggtcggtgaagtcgaacttcgggcacacacaagccgccgccggggtcgtcggcgtcatcaagacggtgctcgcgctgcgccacggcgtcctgccgcagacgctgcacgtggacgctccctcggccaaggtggactggtccgccggttcggtacggctgctgaccgaggcgcggccctggccacgggagagcggacgtacgcgccgggcgggggtgtcctcgttcgggctcaccggcaccaacgcgcacgtgatcctggaggaggcgccgggagaggcggcggcaggggcgcgggccgaggttcccgaggaggcgcggtgcgcctcctcaccggctcgactcccggagccgtccggcgacgcggccgcgccctgggtgctgtccgcccggagccgggcggcgctgcgcgcgcaggcgctccgcctggccgaccaggtggccgccgaccccggtctacgggcccaggatgtcgcccatgccctggccacctcccgtaccctgcaccggcaccgggccgtcgtcagcggctccgaccgggcacaaatgctcgccgcggcaaagcggttcgggctcggtgagcggaccgcgggcgtcaccccggacgattccgcgccgggcctgctggccttcgtcttctccgggcagggcagccagcgcagcggcatggggcgcgcggcggccgaggcgttcccggtcttcggacgggcgctgggcgaggtgtgcgccgcgctggacccgctgctgacacgcccactgacctcggtgatgtgggcggctcccggctccgaggaggcggcccgtctcgacgacaccacctacacgcagcccgccctgttcgccgtccaggtcgccctgtaccggctgttcgagtcctggggcgtggtgccggaccagctggtggggcattcggtcggcgagatctccgccgcccatgtggcaggcgtgctcggcctccgggacgcgtgcaccctggtggcggcccgtagcaggctgatgggcgcgctgccgcccggcggtgcgatggtggcggtacgcatcacggaacccgaagtgaccccatggctcgcggagttgacggacgaggtgtcgatcgcggccgtcaacggtccgcactccctcgtgctcgcgggcgccgaggccccgctcgtcgccctcacggaccggctcgccgccgccggacacaagacccggcgcctcatggtgagcaccgcgccccactcgccgctgatggaccccatgctggaggagttccgcgcggtcgtccgcacgctgtcctacgccgcgcccgccgttcccctcgtctccaccgtcaccggccgcccgctgaccggcgaggaggcgcgcgacccggaccactgggtgcggcatgtgcggcagtccgtccgcttcaaggacgcgatcggccggctccgggacgaacgcgtcaccgggttcctggagctgggtgccgaaccggcactcacaccgatgatcgacgagtgcctggagtccgccgacgggcagcccgggaccgccctggtgccgagtctgcgcgccggagtgccggagcgggatgccctgctcaccgcggtcgcccgggtgcacgcccagggcgttcccgtcgactgggacgcggtgctccccggggcccggcccgttgccctgccgacgtatgcgttccagcggcggcggttctggctggcgtcggctccggcgggctcggcggggccgaccgccgacgggggcttcgcgggcgtcgcggataccgcggacgggacggcggccggtgaagagccgccggggctggaggcccggctgtccggtctcgacgaggccgaacagcacgcgctcgtactggccctggtcctcgccgagacctcggccgtgctcggcggccaggagacgcctggcgaggagccgcacggcgaggagggccaccgcacgttcaaggagatgggcatcaactcgttgaacgccgtcgaactgcgcaaccgccttatcgcagccacggacctgcggcttcccgccacgctcgtctacgactaccccacgccgaaggccgtcgtccgactcgtacgcgaacgactcgcgcgaccggcctcccccgcacgcgatgtggcctccgtcgtggccgagttggagtccctgctgacggccggtgcggaggtctcggaggagaccgcggcgcggttgaaagcggtgacggcggtgtcgacggggacgacggggtcgggcagcggtacgggcgccggctccggcggggctctggacctggtatcggccagtgacgaggaactgttccggctgatggacgcggagagctga |
| *ACP1* | gctgggctaggagatgcagaacaaagggaggcggtgttgggcatcgtgctgggtcaagcggcagttgttctcggccacagcgatgctaccggtattgatcgtcagcgtagcttcaccgctttaggcttcgacagcctgacctctgtcgagctgcgcaaccagctgggcacctccaccggtctgcgtttgccaccgacgctggtttttgaccacccgactccggtggccttggcggagtacctgcatggtgaagta |
| *ACP9* | GGCGGGATGACCGGGGCCGAGCAGCTCGCGTACCTCCTGGGGGTCGTCCGGGCCGAGGCCGGTGCCGTGCTCGGCCATCCCGACCCGGCGGGCATCGCCGGCGACCAGCCGTTCCCGTCGCTCGGCTTCGACTCGCTGACCGCCGTGGAACTCCGCAACCGGCTCGACGACGTCGTCGGCGGCAGGCTCCCCGCCACGCTGGTGTTCGATCACCCCACCCCTGTGGCGCTCGCGGAGTACCTCCGCACGGTCCTGCTGGAG |
| *TADH2* | atgtctactacagtcggccaagttatcagatgtaaggccgccgttgcttgggaagccggtaagccattggttatggaagaagtcgatgttgccccaccacaaaagatggaagttagattgaaaatcttgtacacttctttgtgtcacaccgatgtctacttctgggaagccaagggtcaaaacccagtcttcccaagaattttgggtcacgaagctgctggtattgtcgaatctgttggtgaaggtgttaccgatttggctccaggtgaccatgtcttaccagtttttaccggtgaatgtaaggactgtgctcattgcaaatccgaagaatccaacatgtgttctttgttgcgtatcaacactgacagaggtgtcatgttgaatgacggtaagtccagattctccattaacggtaacccaatctaccacttcgttggtacctcaaccttttctgaatacaccgtggttcacgttggttgtgttgccaaaattaaccctttagctccattagacaaggtttgcgtattgtcttgtggtatctccactggtttgggtgcttccttgaacgttgctaagccaaccaagggttcttctgtcgctatcttcggtttaggtgctgtcggtctagccgctgctgagggggctagaatcgctggtgcctctagaatcattggtgttgacttgaatgcttctcgttttgaacaagctaagaaatttggtgtcactgaattcgtcaacccaaaggactattccaagcctgttcaagaagtcattgctgaaatgactgatggtggtgttgatagatccgttgaatgtactggtcacattgatgctatgattagtgctttcgaatgtgtccacgatggttggggtgttgctgtcttggtcggtgttccacacaaggaagcagtattcaagacccatccattgaacttccttaacgaaagaaccttgaagggtactttcttcggtaactacaagccaagatctgacattccatgtgttgttgaaaagtacatgaacaaggaattagaattggaaaaattcatcactcacactttgccattcgctgaaatcaacaaggctttcgacttgatgttgaagggtgaaggtttgagatgtatcattactatggctgactga |
| *rimB (rimM1)* | atgtcgaacgaggagaagctgcgcgagtacctcaagcgcgcgatcgcggacctccacgagacccgtcagcagttggacgagaccgaggcgaagcagcgcgagcccatcgcgatcgtctcgatggcctgccgcttccccggcggcgtccggtccccggaagacctctgggacctgctgcgggacggcgtcgacgcgatctcctccttcccccgcaaccgcggctgggacctggacacgctctaccaccccgagccgtcccaccagggcaccacctacgcccgtgagggcgggttcctgcacgaggcgggcgagttcgaccccggcttcttcgggatctccccgcgcgaggcgctggccatggacccccagcagcggctgctgctggagaccgcgtgggaagccgtcgaacgcgccggcatcgacccggaatccctcgcgggcagcagcaccggcgtgttcgtcggcaccggccacggggagtacgacaccgagggcggccggcgcgccgacgaggtcggcgggcacctgctgaccggcaaccacatcagcatcgcctccggccggatctcctacgtgctgggcctggagggccccgcgctgaccgtggacacggcctgctcctcgtcgctggtcgccctgcacctggccgcgcatgcgctgcggcgcggcgagtgctccatggccctggtgggcggcgcgaccgtgatgtccacgccgaagatcttcgtggagttctcccgccagcgcggcctggcgcccgacggccgctgcaagccgtttgcggccgccgccgacggcaccggctggagcgagggcgtcggcatgctgtgcgtcgagcggctgtcggacgcggtacgcaacggccatcccgtactcgccgtactgaagggctcggccgtcaaccaggacggcgcgtccaacggcctgaccgcccccaacggcccctcgcagcaacgcgtcatccgtcaggcgctcaccggagccgggctggccgccgcggacatcgacgccgtggaggcgcacggcaccggcaccaccctcggcgaccccgtggaggcgcacgcgctgctggccacgtacggacagcagcgccccgccgaccggcccgtgctgatcggcgccatgaagtccaacatcgggcacacccaggccgccgccggtatcgcgggcgtgatgaagatggtcctggcgatgcggcacgggcagctgcccaggaccctgcacctggacgagcccaccgggcacgtcgactggagcgagggcaacgccagactcctcgcggagcccgagccctggccgagcgccgaccggccccgccgcgccgccgtctcctcgttcggcatcagcggcaccaacgcccacgtcatcctggagcaggcacccgcccaggacgccacaccggcccccgaaccggccgcccggccgggcgcgctgcccctggtcctgtccgcccgcaccgaagaggccctgcgcgcccaggccgaacggctcggccgccacctgcgggaccgggccgacctggaaccggccgcggtcgcacgcgccctcgcgagcagccgtacgctcatggagcaccgcgcggtcgtcgtcgcggacgaccgggaagcgctgctcagcggcctggacgcgctggccgccggccgtaccgccaccggcctggccggcggcgtcgccgtcaacgcccccaccgcgttcctcttcgccggacagggctcgcagcgcgccggcatgggacgcgagctgtacgcggcgtaccccgtgttcgccgcggccttcgacgcggtgtgcgcggaactcgacccgcacctggaccggccgctgcgcgacatcgtcttcgccggggaaggcagtgacgaggccgccctgctggaccgcaccgcctacacgcagaccgccctgttcgccctggagaccgccctgttccggctggtcgaatcctggggcatggcgccccggttcgtcgccggacactccatcggcgagctgaccgccgcccacgtcagcggcgtactgaccctccaggacgccgcacggctggtcgccgcgcgcggcaccctgatgcaggcgctgcccgaaggcggcgcgatggtggcgatccaggccaccgaggaggagatacgaggccacctcgcggaccgcgagaacgtggccctggcagccgccaacgggcccgactccaccgtcatttcgggcgacgagcaggccgtggccgagatcgcggagcactgggcggcgcagggccgccgcaccaagcggctgcgggtcagccacgccttccactccccgcacatggacggcatgctggaggaattccggcgcgtcgcgcgcggcctgaccttccacgcgccccgtatccccgtcgtgtccacggtgaccggcgcgctcgccgccgaggaggacctgcgctcgcccgactactgggtacggcaggtccgcgaagcggtgcgcttctgcgccgcggtccgcaccctcgaagccgaaggcgtcaccacgtacgtggagatcggccccggcggcgtcctgacgcccatggtccaggacagcctgaccaccctcgaagcgcccgtgctcgtcccgctgctgcgcaccggacgccccgaggcgcacgccctcaccgaggccgtcgccaccgccttcgcgcacggcgagcgtgtcgactggaccgcctgcttgggcgcgcccggcacctcgcacgtcgacctgcccacgtacgccttccagcgccagtggtactggctggacccggccggccgcgacgaggagcagacggccgccgccgaagcgggcgaggccggcttctgggcggccgtcgaacgcgaggacctccaggaactctcggccgtcctggccatcgacggcagcgaggcggactccctcggcagcctcctgcccaccctctcctcctggcaccggcagcgcaggacgcaggccgccgcggaccgtttcagctaccgcgtccactggaccccgcgcaccgcctccggcggccccgccgccaccggccactggctcgtcgtcctgcccgaaggcggcaccgacgacccgtggaccgcccgcctcctggacgcgctgcacgacctgggcctgcacaccgacgtacgcgaactgcccgccgaccacgaccccgacgcgcccatcgcctggcccgacaccccgctcgacggcgtgctctccctcctggccctggacgagcggccccacccggacctgccctccgtgccccgcggcctggccgccaccaccacactgctgcacgccctggagagcgcgggcgtccaggcgccgctgtggtgcgcgacccgcggcgccgtcgccgtcgaccggcacgacgcgctcgacagccccgtacaggctcagatctggggcctgggccgggtggccgccctggaagcgccgcagagctggggcggcctcgtcgacctccccgagaacctggaccggcgcgccgtctccgcgctgctggacgccctcgcctcggaagaggaccaggtcgcgatccgcccggccgggaccttcgcccgccgcctggaaaggatcgccccaggcggcgacaccggcgcccggtggagcatccacggcaccgtcctggtcaccggcggcaccggcgccctcggcggccacctcgcccactggctggccgacgccggggccgaacacctcgtgctcaccggccgccgcggcccgcaggcccccggcgcccaggaactcgccgccgcgctcaccgaccggggcgtcaaggtcaccctcgccgcctgcgacgccgccgaccgcgaagcgctggcggccgtcctcgcggacatcccgccggagctgccgctgaccggcgtcgtgcacgccgcgggcgtcctggacgacggtgtactggccgcgctcacccccgagcgcttcgagaccgtactgcgccccaaggcgcgcgccgcacagcacctgcacgacctcacccagggcatggacctggacctcttcgtcctgttctcctcgatcgtcggcgtcctgggcaacgccggacaggccggctacgccgccgccaacgcccacctggacgcccttgccgcacaccgccgccaacagggcctcccggccacctccgtgtcctggggcccctgggcgggcgagggcatggccaccgacagcgacgcggccgaccggctgagccgcgacggactgctgcccatggccgcggcccccgcactcgccgccctgcgccaggccctcgcccaggacatgacgcacgtgaccgtggccgacatcgactggagcgcgtacgccccggccctgaccgccgtccgccccagccccctcatcggcgacctgcccgaggcccgccgcgcgctcggccccgcggacggcccgcgccgggaacgctcccccctgcgcgaccggatcgccgccctgccgcccgccgagcaggaacaggccctcgtcaccacggtcagggaagaagccgcgaaggtcctcggacacccctcgccggacaccgtcgacgtccagcgcgccttccgcgagcagggcttcgactcgctgatgtcggtcgacctgcgcaaccggctctccgccgccaccggcctgcggctgcccgccaccctgctgtacgaccacccctccaccgtcgcggtcgccgcgtatctgcgctccgaggtc |
| *TR1* | cgggcggaactgtcggcggacggttccgcgagccggccgggcgtggacttcgccgccgaggtccggctcgccgacgacgtccggccggccgacgaggtgatcaccacggccgcggacccgaaggagatcctgctgaccggtgcgagcgggttcctcggcgcgttcctgctgcgcgacctgatgcggaccaccaccgcccgggtgcactgcctggtccgcggggcggacgaggcggcggccatggagcggctgaaggcgaacgccgagtggtaccgggtgtgggacgacatcgacccggagcgggtctcggtcgtcctcggcgatctcgccgagccgcggctgggcctggacgaggagagcttcgacgcgctcgcccgtaccgtcgacgtggtctaccacaacggcgcgcgggtgcactggctgctgccgtacgagaccctcaaggcctccaacgtcacgggtacggaggaggtgctgcggctggccgcccggcaccgcacggttccggtgcactacgtttccaccgtcggcgtgttcgacggggtgcgggagcccggcgtgccgctgagggtcaccgatccgaccgggcccgccgagtccctgccgagcggatatctgcgctccaagtgggtggcggagcaggtcatcgaggtggcccgggaccggggtctgcccgtgtcggtctaccgggtggacgtcatctccggcgaccgggtgaacggtgcctgccagacccgtgacttcgtctggctgacgctgaaggggctcatccaggcccgcagtgtgcccaagggcaccgagggccgcttccacctgctgccggtggactacgtgagcgccgcgatcaccggtatctcccggcagcccggtacggtcggccgtaccttccacctcttcaaccagagctcgctggcgctcagccagtgcgtggagctcctgcgctcgctcggatacgagctggacgaggtcgactgggacacctggactcaggtggtgacgtccggtgacaacgcactgctgccgctgctcgacgccttcgagatgatgacgtcggacaccgacggcttctacccgccgatcgacaccgctgagacggtggcggcgctggagggaaccggtatcgggattccggtgctgacgcgggagctgttcgagaagtatgtggccttcttcgtcgaggagggacacttcccggcggccgggtga |
| *TR2* | accggggccagcggctccggggccgcggacacggaagcctccggcaccgtggactacgcggcggacatccagctcgcggacgacatccgccccgccggcgaggtcgtgcgtaccgccgtcgatccgcgcgacgtactgctcaccggcgccagcggattcctcggcgccttcctgctgcgggatctgatgcgtaccaccaccgcccggatccactgcctggtccggggcgccgacgactccgccgcgtacgagcggctgcgcgagagcctggagtggtaccgggtctggggccagatcgatccggagcggctcttcgtcctcgcgggcgatctggccgagccccggttcggtctcgccgaggacaccttcgacggcctggccgttaccgtcgatgtcgtctaccacgccggggcgaccgtgcactggctgcatccgtacccggccctgaaggcggccaatgtcggcggtacggaggagattctgcggctggccgcgcgccaccgtacggtgccggtccactatgtctccacggtcggcgtgttcaacgggcccgtgacaccgggcgttccgctgaaggtgaccgatcccacgggaccggccgaagccctgcccagcggctatctccagagcaagtgggtcgccgaacaggtgctggacctggccagggagcggggcattccggtgtcggtgtaccgggtcgacgtgatctccggcgaccaggtcaacggcgcctgccagacccgcgacttcgtatggctcacgctcaagggcctgcttcagtcgggggccgtgccgtccggtacgggagggcgcttccatctgctgccggcggactatgtgagcgcggcgatcctcggggtctcgcgcgatcccggctcggcgggcggcacgttccacctgttcaaccgcagctcgctgagcctggcggactgtgtggagcggctgcgcggtctggggtacgaactccgggacaccgggcgggaggagtggaccgaggcggtgcgctcggaccgcgacaacgcgctgctgccgctgctccacgccttcgagatgatgaccacggacacggacgccttctatccgcccatcgacacggccgagacggaccgggcgctggcgggcaccgggatcgtctgcccgccgctgacgggtgaactcttcgacaagtacgtggagttcttcgtggcggagggccacttcccggcggccccggcccgctag |
| *TR3* | ggggggtccgccgacccccgttccacggtcgactttgccgccgagatccgacttgcgcccgacatcgtcgccgtcccggacgtgcacctcacggacgacccccgccacatcctgctcaccggcgcgacgggcttcctgggctccttcctcctccgtgacctcctgcggtcaacgtccgcgcgcatccactgcctggtgcggggggcggacgcgagcgacgctcgggcccggctcacggccgcagcggcctggtacgaaaccggcgccgacctcgacttcgaccggatcgacatcgccgtgggcgacctcgccgagccggccctgggcctcgacgaggccgcgttcgacgagctcgccagaaatgtcgacgtcgtctaccacgccggcgcgtccgtcaactggctgtacccgtacgaggccctgcggcccgcgaacatcgccggcaccgaggaggtactgcgtctggccgcgcgacaccggaccgtcccagtccactacatctcgtcgactggagtgtacgcgcaggagccggcagagggccggcggattgccgtggacgacccgatcggcccgccggagctgctttctaacgggtatcggcaggccaagtgggtcgccgagggcatcatcggcatcgcgcggagcaggggcatcccggtgtccgtgtaccgcgtcgacgtcgtgagcggggaccaggtgaacggcgcgtgccagacccaggacttcgtctggctgagcatccgcgggatgctggaagccggggccgcccccgcgggaatatccggtttcttccacccaacacccgtcgactacgtgtccgcggccatccgctggctgtcgtcgcgggtcaccggcgtcggtgacaccttcaacctgagcaacccgcatcggctccatttcgccgaggtcgtcgagcgcctacgggcgctcggccacaccctcgtggacctggacccagccgagtggtcgcgttccgtgaggaaagaccccgagaactccctgctgccgctgctcgacgtgttcgaggcggcgatcgccggaaccggtggctacccggacatcgacaccgacggaacggaggccgcactggccggctccggtatcgactgcccacaggtcaccggcgacctcctggtccggtacctgacgttcttcaccgagaagggctacttcccgaaccccgatcgcgatggcgcggtcagcgtcgaggcgtaa |
| *TR4* | ggcgcaccgggcggcggcagggtcgtggacttcgcggccgaggtgcggctggccgacgacatccgggccgcggacaccgtgcacaggacggacgatccgcggcaggtgctgctgaccggcgccaccggcttcatcggctcgttcgtgctgcgcgacctgctgcgtaccacctccgccaccgtgcactgcctggtgcgggccgccgacgaggaggcggcgtacgccaagatccgtgccgcttccgcgtactaccggacgggcatcgagctggaccgggtgcgcgtggtcctgggcgacctcggtgcgccggggctggggctcgacgcggcgacggcggaccggctggcccgcgaggccgacgtggtgttccacatcggcgcccatgtcaactggctctatccgtacgcgaagctgaaggcggccaacgtcaccgcgacggaggagatcctgcggatcgcggcgaagcaccgcacggtgccggtgcactacatctcgtccaccggtgtctacgcgaacctgccggaggagggtgtacgcctctccgaggacgcgccgatcggtccgccggagcagctcctcagcggctaccggcagtcgaagtgggtgtgcgaggagatcatcggcatcgcccgcgagcgcggtgtcccggtgtcgtcgtaccgggtggacgtggtgacgggcgatcaggtcaacggcgcctgccagacacaggacttcgtctggctgagtatccgcggcatcgtggaggcgcaggccgtgccggacagtctggagagccacttccacccgacgcccgccgactacgtcagcggcgcgatgctccagctggcgtggcggctcaccggcaccggtgacacctacaacctctcgaacccggagcggttgacgttcggggagatcatcgacgcgctgcgggcccggggccaccgcatcgacgacctggacctcacggcgtggacccgcaaggtccgctcggaccggtccaacgcgatgcagccgctgctcgaccagttcgtggggctcgtggggctgccgggcggcacctacccgagcatcgactgctccaaggccgatgccgcgctggcggacacctcggtcgtctgcccgcccgtgcggagcgacctgctggagacctacctggacttcttcaccgaagtcggttacctgcccgccccgtga |
| *TR6* | ggcgggccggacggcgacgagggcgggaccgcggcggccgacccggcacgggagtacgcggcggagatcgtgctggacgaggcgatccggcccgccgacgaggtgctgcaccaggtcgaggacccggcggaggtcttcctgaccggcgcgaccgggttcctcggcgcgttcctgctgcgcgacctgctgcgggggactcgggccacgctgcactgcctggtccgtgcccgggacgcggcggacgggctgcggcggctgcgggagaacctggagtactaccgcgtgtgggacgaggtggacccggcccgcctggagatcgtcgtcggggacctcgcagagccgcttttcgggctgagcgaggaggagttcgacgacctcgcccgccgggtcgacgtggtctaccacggcggcgcgaaggtccactggctgcacccgttcagctcgctcaaggcggccaacgtgggcggcacgcgggaggtcctgcggctcgccgcgcgccaccgcaccgtcccggtccactacctgtccacgaccggggtcttcgccaccgaccgggcggagaacagcccgctgagggcggacgatcccaccggccccgccgaggcgctgcccagcggctatctgcgcagcaagtgggtggccgagcaggtcgtcggcatcgctcgcgacagggggctgccggtgtcggtgtaccgcgtggacgtcgtctccggcgaccaggtcaacggggcctgccagatgcgtgacttcgtctggctgagcctgcgcggtctgatccaggcgggcgcgtacccggccgggctggccgccgcggtccccctgaccccggtggactacgtcagctcggccgtggtcgccctgtccacgtcggccgggaccggttcgggcaccttccatctctacaaccagagtcacatgacgttcgccgacttcattaccgagttgagggcgacgggttacccgctgaaggaagtcgagtgggacgcgtggagcacactcgtgaggtcggaccccgacaacgtcatgcttccgctgctggaagcgttcgagatgatggcgaagagcgagggcaccttctatccacccgtggacacgagcgtggccgaactggccctggcgggcccgggagttgagtgccccgagatggagccggagctgttccggcggtacgtcgccttcttccgggaggccggtttccttccgccggtgaacgaggacgcgcggcccgctgcctga |
| *TR7* | ggcggacggacgcagggcccggcggccaagacggacgtcccggacttcgccgctgaggtgcggctggccgaggacatccggcccgcggccgatgtggtgtccgtggccgacgacccccggcacgtgctgctgaccggcgccagcggtttcctcggcgccttcctgatgcgcgatctgatgcgcacgacacgggcgacggtgcactgcctggtgcggggtgaggacgaggccggggcgcacgcccggctgcgggagaacctggaacgctacggcgtctgggacgaggtggacgccgaccgtctctcggtggtcgtgggcgatctggcgcagccgcggctcgggctggccgaggacgccttcgaccatctggcgaggacggtcgacgccgtctaccacaacggcgcccgggtgcactggctgcacccgtacgcgacgctcaaggcggccaacgtcctcggcacggaggagatcctgaggctggcggcgcgccaccgcacggtgccggtgcactacgtgtcgaccgtcggcgtcttcgacggcacggcggccaggggcgtaccgctgcgggtgaccgacccgaccggtcccgccgagcggctgcccagcggctatctgcagagcaagtgggtcgccgagcggatcatcgggatcgcccgtgaccggggcctgccggtgtccgtctaccgagtggacgtcatctcgggcgaccagcgcaacggcgcctgccagaccagcgatttcgtgtggctgagcatcaagggcctactgcaggcgggtggggtcccggccgatgtcgggggtcgtttccacctgcttcccgtggactacgtcagcgcggccatcctccgcatcgcgagccgcccctcggccgcgggcggcaccttccatctcttcaacccgagttccatcagcctgcgggagtgcattcgccatctgcggtcgttcggatactcgctgaccgagctggactggaacacctggcgggcccgcgtgcatgacgacccggccaacgccatggcgccgctgctgcacgccttcgagatgatgacggccgacaccgacgcgttctatccgcccatggacacaacggagaccgaggcagcgctggacggcagtggcatcgtttgcccgccgttgacccgagaactcttcgagaagtacgtacggttcttcgtcgagacggggcacttccccgcggcaggctga |
| *TR8* | gcgccccaggacgacaccgggccgggcccggccgtggactacgccgccgacctcgacctgcccgccgacatccgcccggccggcgaggtcgtccgcaccgtcaccgacccgtcggacctcctcgtcaccggagcgagcggcttcctcggagcgttcctggtgcgcgacctcatgcgcaccacgaccgcccgcctgcactgcctggtgcgcggcgccgacgacgcggcggcgtacgagcggctgcgctccagcctcacctggtaccgcgtgtgggacgagatcgacgagagccggctgcgggtccgcgccggggacctggccgaggagcgactcggcctgaccgaggaggagttcgacggcctcgcccacaccgtggacgccgtctaccacgcgggcgctaccgtgcactggctccacccctacgaggcactgcgcgcggccaacgtccacggcacccgcgagatcctgcgcctcgccgcccgccaccgcaccgtcccggtgcactacatctccacggtcggtgtcttcaacggcccggtcacccccggcgtccccctgaaggtcaccgacccgaccggccccgccgaagcgctgcccagcggctacctccagtccaagtgggtcgccgaacagctcgtcggactcgcccgggagcggggtctgccggtgtccgtgcaccgcgtcgacgtcatctccggagacacccgcaacggggcctgccagacccgcgacttcgtctggctcagcctcaagggcctcctccaggcgcaggccgcaccggccggcgtcgcaggccgcttccatctgctgcccgtcgactacgtcagcgccgccatcgtcgggatctcgcggcgcgagccggccggcggcaccttccacctgttcaaccggagctccctgagcctcgccgactgcgtgtcgtacctgcgcgaactcgggtaccggctcgacgagacggaccgggagcggtggagcgccgcggtgcggtccgaccgggacaacgccctgctgccgctgctgcacgccttcgacatgatgacgtccgacaccgacggcttctacccgcccatcgacacggccgagaccgaggcggcactggccggcacggacatcgcctgcccgccgctgacgcgcgaactcttcggcaggtacgtggagttcttcgtccaggaagggcacttcccgcccgccccgtga |
| *TR9* | ggcaccacgggcagcggcgcgggtagggccggggccaagggcgacggcggcgccgcccatgccgtggacttcgccgccgagacggtcctgcccgaggacgtccgccccgaaggcgtggtcacccgagttgcgaccgaccccacgacggtgttcctcaccggcgccacggggttcctgggcgccttcgtactgcgggacctgctgacgaccaccaccgcccgcgtccgggtgctcgtccggggcgccgaccaggccgacgcccaggagcggctgcgcgccaacctggactggtaccgcatcgccgaggagatcgatgagagccgcatcgacgtcgtggtcggcgacctggcgcgtcccctgctcggactcgacgagggggagttcgaccgtctgtcgcgcgagaccgacgtggtctaccacgtcggggcctccgtcaactggctgcacccctacgaggacctgaaggcggccaacgtctcgggtaccatcgaggtcctccggctcgccgcccggcaccgctcggtgcccgtgcactacgtgtccaccaccggcgtcttctccggaacggactcgggaggcgccgcgctcgcgccggacgcgccgaccggccccgcggagtccctgccgacgggttacgtgcagagcaagtgggtctgcgagcagctgatcggcacagcccgggagcgcgggctccccgtcagcgtgtaccgggtcgacgtcatctccggcgaccagcggaacggcgcctgccagacccgcgacttcgtctggctgtcgctcaagggcatcctccaggcgggcgcggtgccggagggcatggtcggcccggtccacctgatgccggtggactacgtctcggcagcgatcctgaccatgagcgggcgagagggggccgtcggccgcaccttccacctctacaacccgtccgagctgaccttcgcggaggcggccgagcacctgcgctccttcggctacccgcttggcgaactggaccgggacgcctggctcgaactggtgcggtccgaccgcggcaacgccctggtcccgctgctcgacgcgttcgagatgctcacggcggactcctcaggcttctacccgccgatggacatcaccgacacgctggaggtgttagctggttcgggggtgcactgcccgcccgtgaccaagcagctcttcggccgctacgtcgacttcttcaccgaggtgggctacttcccggcggcccccatgaccgaagga |
| *rimM13 (excluding TE)* | gagccggtcgtcatcgtcggcatggcctgccggctgcccggcggcgtgcggtccccggaggacctgtggcggctcgtcaccgacgagcgcgacggcatctccgccttcccgaccgaccggggctgggacctggagaccctgctcggcggggccgagggcgaccagggccgcagctccgcgtccaagggcggcttcctggacggcctcggcgacttcgacgcccccttcttcggcatctccccgcgcgaggccatggccatggacccgcagcagcggctgatgctggaaacctcctgggaggccgtcgaacgggccggcatcgacccggtgtccctgcgcggcaccgagaccggcgtcttcgtcggcaccagcggctccgactacgccagcatcctgatgaactccggcgaggacgtggaggcccacgccagcaccggcctcgcgggcagcgtcctgtccgggcggctgtcctacaccttcggcctggagggcccggccgtcaccgtcgacaccgcgtgctcctcctcgctcgtctccctccatctggcggcccatgcgctgcgcaacggcgagtgctcgctcgcgctggccggcggtgtgacgctgctgtccacgccgatgagcatcgccggattcagccggcagggcgcgctggcgacggacggccggtgcaaggcgttctccgacgacgccgacggcacgagctgggccgagggcgtcggcgtcctcgtactggaacggcagtccgacgcccgccgcaacggccacgagatcctcgccgtcgtccgcggttcggccgtcaaccaggacggcgcgtccaacggcctgaccgcccccaacggcccctcccagcagcgcgtcatcggccaggccctggccaacgccgggctgtccgccgacgaggtggacgcggtcgaggcacacggcaccggcacctcgctcggtgacccgatcgaggcccaggcgctgctcgccacctacggccaggaccggcccgaggggcagccgctgctcctcggctcgctgaagtcgaacatcggccacaccatggcggcggccggtgtctccggcgtcatcaagatggtcctggcgctgcggcacggcgtcctgccgcgctccctgcacatcgacacgccctcctcccacgtcgactggacggcgggagacgtccggctcctcaccgagcggaccgactggccggacaccggccgcccgcgccgcggcgccgtctcctccttcggcatgagcggcaccaacgcccacgccatcctcgaacaggcaccggcggacgccgggcggccggccgaggacacccggcccgcccagccgggagccgtcccgtggctcgtctcgggcaagtccgcccaggcgctggacgacgggatcggacagcttcgggagtgggtcgccgagcggccggaactcgcgccggccgacgtgggcttctcgctggccaccgggcggccgctgttcgcgcaccgcgcggtgctgctggacggtgccgaggtggcgcgcggcaccgccgaggaccgctcgctggccatgctcttcccgggccagggcggacagcgcatcggcatgggccgcgagctgtacgaccggttcccggtgttcgccgaggccctggacgcggtactggcacacttcgacgacgaactgcgcgaggtgatgcgcagcgacgccgagcggctggactgcaccgagttcacccagccggccatgttcgccatccaggtggcactgttccggctcgtcgaatcctggggcgtacggcccgacttcgtcggcgggcactcgttcggcgaaatcgccgccgcgcacgtcgccggggtcctctccctcaaggacgcctgcaccctggtggccgcccgcgggcggctgatgggcaccctggagggccgcggcgccatcatggtggcggtccaggcgagcgagcaggaagtggccgcgcggctggtcgacggcgtctcgatcgcggcggtcaacggaccggagtccgtcgtcatcgccggcgacgggcccgccacccagcgcatcaccgacgcgttcgccgccgagggccgcaagacgcggcaggtggcggtgcaggtcgccggccactgcccgctgatggacccgatcctggacgagttccgccaggtcgccgagggcctgtcctaccaggagccgcgcatcccgctcgtctccaccgtgaccggcgggcgggccccggaggagctgatgcgctccgccgactactgggtccgcaacgtgcgtgagcccgtccgcttcgccgacgcgatccgcaccctggccgacgaggacgtctcggccttcctggagctgggcctcgacggggcactgacgggcatggtcccgcacaacctggatggcaccgccgtcaccgtcagcgccctgcgcaaggaccaccccgaagagacggccctgctcaccaccctggcgcagctgcacgtcgccggggtcgacgcggactgggcccgcgtcttcgacggcaccggcgcccgccgcgtcgacctgcccacctaccccttccagcgccagcggtactggcccaagcgcagggccctggcgggcgatgtgacctccgccggcctgctgcccaccgaacacccgctgctcggcgcggcggtaccgctggcggacggcggcggcacgctgttcaccagccggctgtcgctggaaacccacccctggctcaaggaccacgtggcgggcggcacggccgtcttcccggccgccggcttcgtcgagctggccctgagcgccggggaccgcttcgactgcgaccggatcgccgaactgaccctcgacacgccgctggggctcaccgacggcgcggccctggtgctccaggtgtgggtcggcgcccccgacgacgacggcacccggcagatccggttcttcacgcggccccaggccgcgctggacgagccctgggtccggcacgccacgggcatcctcgccgccgaccggcacaccgacaccctcgacgtgtccggctggccgcccagcgacgcgaccgagatggacgtggacgagtactacgagggcaccgaattcggacccgccctccaggggctgcgggcggtctgggtccgcgacgacgagaccttcgtcgaagccgaactccaggacgacgccgcctccgacgccccgtccttcggcctgcacccggcactgctcgaagcggtcctgcacggcacctccttcagcggcgtgggtgacagggagcacatctcctacggcgcctcctggaacgacgtgtcgctgcacgcctcgggtgccgcggcgctgcgggcacggatcacgaagaccggcgaggacaccgtcaccgtcaccgcggtcgacgccgccggggaacccgtcctgtcggcccgcaccgtcacactgcgcgcgctggccgaagcccccgagcgcacctccagcggccagagccagaaactgctccgcctggagtggatccccgcaccgcgcacggaggacgcgccgagcgcccggagcaccgtcctggcgcccctgggcaccacgtccctcgccggcctcacggacacccccgacctggtcgtggtccccgtgccccgtacggcggacggctacccggaggccgtgcacgagctgacgacccaggccctgctcctggtacagcagtggatcgccgacgagcgcttcaccacgtcccagatggtgttcctgacctccggcgccgtcgacggcggcgacctggccggctccgccctgtggggcctgctgcgctcgacgatcgcggaacaccccggccgtttcgtgctggccgacgtcgaggacccggacgagaacggcggcgacctcgccacgctcctcgacgccctgccgggtctgctgcccaccggtgagacccagttcgtcgtccgcgacggcgcggtgctcatggggcggctggcctggctgcccgccgacgccggcacccggccggcgcagtgggaccccgaggggaccgtactgatcaccggcggcaccggcggtctgggcatcgagctcgcccggtacctggccacgaccggcacccggcacctgctgctggtcagccgcagtggcccggcggcccgcggggcggacgaactggccgaagagctgcgggccctgggcgcgcagcccacgatcgcggcctgcgacacggcggaccccgccgcgctcgccggcctgctggccgccatccccgaggagcacccgctgaccgcggtcgtccacctcgccggcgtcctcgacgacggcgtggtcaccgcgctgaccccggagcgcctgacgccggtgctgcggccgaaggtggacgccgcctggcacctgcacgaggccaccaaggacctcggcctgtccgcgttcatcatgttctcctccgtctccggtgtgctgggcgcccccggcgtggccaactacgcctccgcgaacgccttcctcgacggcctgatccggcaccgcgccgccctgggcctgccgggccagtcactggtgtggggccactgggacggcggcggcatggccgacgcgatgaaccaggccaccgtgcagcgcatgcgcctcaacggcatggcgccggtctccatcgaggagggcatggaactgttcgacatcgcgcgcggctacgccgagcccgtcgtgatggcggtgggcctgatccccggcgcccgtgtgccggaaggacagatcccgccgctcttcagggggctggtgcggcacagcaggcggaccgccgtcaacgccgcgagcacgtccgcctccccggacaccttcgcccagcagctgctggcgctgccggagggggagcgcgtacgccacctcgtcgacctggtccgtaccgaggccgccaccatcctcggccatgtctccgccgaggagatcgaggccggccgggacttctacgagctgggcatcgactcgctgacctcgatcgagctgcgcaaccggctcgccaccgtcaccgggctccagctcccggcgaccctggtcttcgacaacaagacccccgacgccctcggcgcccggctgcgcagcgagttc |
| *rimM7 AT* | gaggccccccgcccgcaggagacggctcccaccgagccggtgcccccggccggtgacgccgtgccctgggtgctgtccgcccgtaccccgggcgcactgcgcgcccaggcggccaagctcgccgcccacctcgacggcgagcaggcacccggcgcgctggacgtcgcccacagcctggtggcctcgcgcacgctcttcgaccaccgcgccgtcgtcgtcggcaccgatgacaccgcacggcgggccgcactggacgccctcgccaccggcggctccgcgcccggcgtcgtgcagggcacggccgacaccgacggcaagaccgtcttcgtcttccccggccagggctcccagtgggtcggcatgggcgcgcggctcctggaggagtccccggtcttcgccgagcgcctgaccgagtgcgccaccgccctgtcggccttcaccgactggtccctgctggacgtactgcgccagaccgaaggcgcgccgaccctggaccgcgtcgacgtggtccagcccgcgtccttcgccgtcatggtctccctggccgcgctgtggtccgcccacggcatcacccccgacgccgtgatcggccactcgcagggcgagatcgccgcggccgccgtcgccggggcgctgtccctggaggacgcggcccgcgtcgtcgcgctgcgcagccaggccatcgcccgcggcctcgccgggtccggcggcatgatgtcggtcccgctgcccgccgccgaggtcgaacagcgcctcgccgcgtacgagcacctgtccatcgccgccgtcaacggcccccgctccaccgtcgtctccggcgccaccgcgccgctggacgcgctccaggccgaactggtcggcgcggacatccgcgcccggcgcatcgccgtggactacgcctcgcactccgcgcaggtcgagatggtgcgcgacgagctgcacaccgtcctcgcgccggtccgcccgcgcccggccgaggtgcccttcttctccaccgtcaccggcgactggctcgacaccaccgccatggacggcgactactggtacaccaacctgcgccagaccgtccgcttccagcacggcatcggtgaactcctcgcccagggccaccgcttcttcatcgaggtcagctcccacccggtgctgtccatcggcgtccaggagaccgtcgaggaagccggcggcacggccgcggtgctcggcacgctccggcgcgacaccggcggtgtcgaccgcttcctgacctcgctcgccgaggctttcgtacggggcgccggcgccgactgggccgccgtcctcgcggacaccggcgcccgccgcgtcccgctgcccacctac |
| *pimM7 AT* | gaggccccccgcccgcacgaggcgacccccgcggggccggtgcccccggccggggacgccatcccctgggtgctgtccgcccggaccccgggcgcgctgcgcgcccaggcggcccagctcgccgcccacctcgacggcgaggcacccgatgccctcgacgtcggccacaccctggtggccgcgcgcaccctcttcgaccaccgcgccgtcgtcgtcggcaccgacgacgcatcccggcgcgcggccctggacgccctcgccaccggcggctccgcacccggcatcgtccagggcacggccgacaccgacggcaggaccgtcttcgtcttccccggccagggctcccagtgggccggcatgggagcgcggctccttgaggagtccccggtcttcgccgcgcgcctgaccgaatgcgccaccgccctctcggagttcgtcgactggtccctgctcgacgtgctgcgccaggccgacggcgcgccgaccctggaccgcgtcgacgtcgtccagcccgcctccttcgccgtcatggtctccctggccgcgctgtggacctcccacggcatcaccccggacgccgtggtcggccactcgcagggcgagatcgccgcggccgccgtcgccggggcgctctccctggaggacgcggcccgcgtcgtcgcgctgcgcagccaggccatcgcccgcgggctggccggaaccggcggcatgctgtcggtcccgctgcccgccgccgacgtcgaacagcgcctcgccgcctatgaggacctgtcgatcgccgccgtcaacggcccccggtccaccgtcgtctccggcgccaccgcgccgctggacgccctccaggccgaactggtcggcgaggacatccgtgcccggcgcatcgccgtggactacgcctcgcactccgcccaggtcgagcgggtgcgcgacgaactgcgcaccgtcctcgaaccggtgcgcccgcgcccggcgcaggtgcccttcttctccaccgtcaccggtgactggctggacaccaccgtcatggacgcggagtactggttcaccaacctgcgccggaccgtccacttccagcccgccatcggcgaactcctcgcccagggccaccacttcttcatcgaggtcagctcccaccccgtcctgtccatgggcatccaggcgaccgccgaggaagccggggccgccgccgcggtgctcggcacgctccggcgcgacaccggcgccaccgaccgcttcctggcctccctggccgaagccttcgtccggggcgcggacgccgactggtccgccgtcttcgccggcaccggcgcccgccgggtgccg |
| *rimM13 AT* | caggcaccggcggacgccgggcggccggccgaggacacccggcccgcccagccgggagccgtcccgtggctcgtctcgggcaagtccgcccaggcgctggacgacgggatcggacagcttcgggagtgggtcgccgagcggccggaactcgcgccggccgacgtgggcttctcgctggccaccgggcggccgctgttcgcgcaccgcgcggtgctgctggacggtgccgaggtggcgcgcggcaccgccgaggaccgctcgctggccatgctcttcccgggccagggcggacagcgcatcggcatgggccgcgagctgtacgaccggttcccggtgttcgccgaggccctggacgcggtactggcacacttcgacgacgaactgcgcgaggtgatgcgcagcgacgccgagcggctggactgcaccgagttcacccagccggccatgttcgccatccaggtggcactgttccggctcgtcgaatcctggggcgtacggcccgacttcgtcggcgggcactcgttcggcgaaatcgccgccgcgcacgtcgccggggtcctctccctcaaggacgcctgcaccctggtggccgcccgcgggcggctgatgggcaccctggagggccgcggcgccatcatggtggcggtccaggcgagcgagcaggaagtggccgcgcggctggtcgacggcgtctcgatcgcggcggtcaacggaccggagtccgtcgtcatcgccggcgacgggcccgccacccagcgcatcaccgacgcgttcgccgccgagggccgcaagacgcggcaggtggcggtgcaggtcgccggccactgcccgctgatggacccgatcctggacgagttccgccaggtcgccgagggcctgtcctaccaggagccgcgcatcccgctcgtctccaccgtgaccggcgggcgggccccggaggagctgatgcgctccgccgactactgggtccgcaacgtgcgtgagcccgtccgcttcgccgacgcgatccgcaccctggccgacgaggacgtctcggccttcctggagctgggcctcgacggggcactgacgggcatggtcccgcacaacctggatggcaccgccgtcaccgtcagcgccctgcgcaaggaccaccccgaagagacggccctgctcaccaccctggcgcagctgcacgtcgccggggtcgacgcggactgggcccgcgtcttcgacggcaccggcgcccgccgcgtcgac |
| *fkbS* | ATGAAGGAAATCCTGGACGCGATCTCGTCGGCGGATGCGACGCCGGCGGACTTCGCCGCCCTCGCAGTCCCCGAGTCCTACCGCGCGGTGACCGTGCACAAGGACGAAGCCGAGATGTTCGCCGGCCTGCCGAGCCGGGACAAGGACCCCCGTAAGTCGCTGCATGTCGAAGACGTCGCGGTGCCCGAACTCGGGCCCGGTGAGGCGCTCGTCGCCGTGATGGCCAGCTCCGTCAACTACAACTCCGTGTGGACGTCGATCTTCGAGCCGCTGTCCACCTTCGGCTTCCTGGAGCGCTACGGGCGCGTCAGCGAACTCACCCGGCGGCACGATCTGCCGTACCACGTCATCGGCTCGGACCTGGCGGGCGTCGTCCTGCGCACCGGGCCCGGGGTGAACGCCTGGAAACCGGGGGACGAGGTCGTCGCCCACTGTCTGTCGGTGGAGCTGGAGTCCTCGGACGGGCACAACGACACCATGCTCGACCCCGAGCAGCGGATCTGGGGCTTCGAGACCAACTTCGGCGGCCTCGCCGAACTCGCGCTGGTCAAGTCGAACCAGCTGATGCCCAAGCCAGCCCATCTGTCCTGGGAGGAGGCCGCGGCGCCGGGGCTGGTGAACTCCACCGCGTACCGCCAGCTGGTCTCCCGCAACGGCGCCCGGATGAAGCAGGGCGACAACGTCCTGATCTGGGGTGCGAGCGGCGGGCTCGGCTCGTACGCCACCCAGTTCGCGCTCGCCGGGGGCGCCAACCCGATCTGTGTGGTCTCCAGCGACCGCAAGGCGGACATCTGCCGGTCGATGGGCGCGGAGGCGATCATCGACCGGAGCGCCGAGGACTACCGGTTCTGGAAGGACGAGCGGTCGCAGGACCCGCGTGAGTGGAAGCGGTTCGGCGCCCGGATCCGTGAGCTGACCGGCGGCGAGGACGTCGACATCGTCTTCGAGCACCCCGGCCGGGAGACCTTCGGGGCCTCCGTGTACGTCACCCGCAAGGGCGGCACGATCGTCACCTGCGCCTCGACCTCGGGCTATCAGCACGAGTACGACAACCGCTACCTCTGGATGTCGCTGAAGCGGATCATCGGCTCCCACTTCGCCAACTACCGGGAGGCGTGGGAGGCCAACCGGCTGATCGCCAAGGGGAAGATCCACCCCACGCTGTCGAAGGTGTACCCCCTGGCGGAGACCGGCCAGGCGGCGCACGACGTCCACCGCAACGCCCACCAGGGCAAGGTCGGCGTCCTCTGTCTGGCACCCCGTGAGGGCATGGGTGTGCGGGACGAGGAGACGCGCGCCCGGCACCTCGGCGCCATCAACCGGTTCCGCAATGTCTGA |
| *rimJ* | ATGAAGGAAATACTGAACGCTATTCAGGCGCCGGACACCACGGCCGCCGACTTCGCCGCGCTGCCGCTGCCCGAGTCGTACCGCGCGATCACCGTCCACAAGGACGAGACCGACATGTTCGAGGGGCTCGCGGGGTGCGACAAGGACCCGCGCAAGTCGCTGCACCTCGACGAGGTGGCCGTCCCCGAACTCGGTCCCGGCGAGGCCCTGGTGGCCGTCATGGCGTCCTCGGTGAACTACAACAGCGTCTGGACCTCGATCTTCGAGCCGCTGCCGACGTTCGGGTTCCTGGAGCGGTACGGCCGGCGCAGCGAGCTCGCCAAGCGGCACGACCTGCCGTACCACGTCATCGGCTCCGACCTGGCGGGCGTCGTGCTGCGCACCGGGCCCGGCGTCACCGTCTGGAAGCCCGGTGACGAGGTCGTCGCGCACTGCCTGTCCGTCGAGCTGGAGAGCGCGGACGGCCACGGTGACACGATGCTCGACCCCGAGCAGCGCATCTGGGGCTTCGAGACCAATTTCGGCGGTCTCGCCGAACTCGCGCTGGTGAAGTCGAATCAGCTGATGCCCAAGCCCGGCCATCTCACGTGGGAGGAGGCGGCGGCCTCCGGGCTGGTGAATTCCACCGCGTACCGCCAGCTGGTGTCCCGCAACGGCGCGGGCATGAAGGTCGGTGACAATGTGCTGATCTGGGGCGCGAGCGGCGGACTCGGTTCGTACGCGACGCAGTTGGTGCTGGCCGGCGGCGGCACACCCGTCTGTGTGGTGTCCAACGACCAGAAGGCGGAGATCTGCCGGTCCATGGGCGCGGAAGCGGTCATCGACCGCAACGCCGAGGGGTACCGGTTCTGGAAGGACGAGCAGACGCAGGACCCCCGGGAGTGGAAGCGGTTCGGCAAGCGCATCCGCGAGCTGACCGGCGGCGAGGACGTGGACATCGTTTTCGAGCACCCGGGGCGGGAGACGTTCGGCGCGAGTGTGTTCGTCACGCGCAAGGGCGGCACGGTCGTCACCTGCGCCTCCACCTCGGGCTACCAGCACCAGTACGACAACCGCTATCTGTGGATGTCGCTCAAGCGCATCGTGGGCTCGCACTTCGCCAACTACCGCGAGGCGTGGGAGGCCAACCGCCTGATCGCCAAGGGCAGGATCCACCCGACGCTGTCCCAGACCTACCCCCTGAGGGAGACCGGGCAGGCGGCGTACGACGTGCACCGCAACCTCCACCAGGGCAAGGTCGGCATTCTGGCGCTCGCTCCCGAGGAAGGACTCGGCGTACGGGACGCCGAGTTCCGGGCGCGGCACGCCAGCGCCATCAACCGCTTCCGCGGGACGCTGGAGGACTGA |
| *TA1* | atgaccgaccagccgaccacacgggacggcttcgccgagccgtttctgctggaggtgctcgcctccggcggcctggacgccgcctacgtgcgcgcggagggcaacaccctgtaccggcgcggtgaggacggcgaggagatcgccgtactcgacttcgtcggcggctacggctcgctgatgctcgggcacaacaacccggagatcaacgaccgggccagagaactgctggaccggcagacgcccgtccacgcgcagttctcccgccatccgtacgccgacgaactggcggcggagctgaaccggatcgtccagcgcgaacgcggcgacgacgagtcgtactacgccatcttcgccaacagcggcgcggaggccgtcgaggcggcgatgaagcacgccgaactcgaccgcggactgcggctgtcggcactcaccgaggagatcgacgcccacctggaggaggtccgggcccgggtcgccgacggcaccgccacggtcccgccacgggtcgccgggtccgccgacgagctgatcgcggacgtccggcgcaggaacgaggaacagctggcccgcggcccgctgttcctcaccccggaaggcgccttccacggcaagctcgccggcagcgtccagctcacccacaacccgggctaccggctgccgttcaagtccctcgccgcgcaggccaggttcgtgccccgggaccagccgggggcgctgcgcaagatcgtcgacgaggagcggccgcacctgctcgacctggtcgtggagggcggccgggtacaggtcgtcgagcgggaccacccgctgttcacggccttcgtcctggagcccgtccagggtgagggcggcatccacgagctgtccgccgagttcgtggcggaggtccaggaggtctgtgccgaggcccggatcccggtcgtcgtcgacgagatccagagcggcatgggacgcaccggcagcttcctggcggccacccggctggggctgaagggcgactactacacgctggccaagagcctgggcggcggtatcgccaagtccgcggtgctgctcgtccgcaagccgctctaccacgggcagttcgaactggcccacagctccaccttcgccaaggacgccttctcctgcctcatcgggctcaaggtgctcgagatcatggaggccgacgacggtgcggtctaccgccgggcggcggagcgcggtgagcgcctgctcggaatgctccggtcggtccgggcggacttccccgagacggtgcgggacgtgcgcgggcgcggcctgatgctggggctcgagttccgggaccagtcggacgccacggccgacccgctgcggcaggtcgcccggagcgggttcttcggctacttcgtcgcgggccacatcctgcgggagcaccgcgtccgtgtcttcccgacgtcgagcgcggtgaacacgctgcggttcgagccgtcggtgtatgtcacggacgaggagatcgaccggctggaggccgcgctgcgcgacgtgtgcgcgatcatccgggacaccgacggagcgcggctggctccgatcggctga |
| *TA2* | atggatcacaccaacaccgcaaccccgtccagagaactggccgaaccttttctccgtgaagtactcgccggcgtgggcctgtcggtcgaatacgtccgtgcgaagggcaatacgctctaccaacgggacggcgagggtggcgagataccggtcgtcgacttcgcgggcggctacggctccgtcctgctcgggcacaaccggccggaaatcgtcgaaagggcgcgggaactgctcgacggggacaccccggtgcacgcgcagttctccagccacccctacgccaacaccctcgccgccgagctgaaccggatcatccgccgggaactgggcactgacgaaccgtatttcgccgtattcgcgaacagcggtgcggaagcggtcgaggccgcggtcaaacacgccgagctggaccgggggatgcgcatcgccgcgctcctggaggagatctccgtacagaacgacgccgcgcgcaccgcggtggagaacggcaccgccgtcgtcgccgagcacaccctcgaacggctcggcctcaccccgccccgggaccccgccgaggccttcggtctgctgacggccgagctcgcccggcgcaacgccggacagaccgcgcggccgccgctgttcctggcgctggagggcgggttccacggcaagctcgcggcgagtgtgcagctcacgcacaacgaggggtaccgcacccccttcaagtcgctggccgcgcaggcccggttcgtaccgcaggaccggcccgatgtcctgaagcaggtgcatgccgaggagcggggcgttctcctcggcacgcaggtcgtcgacgggcagctcaccctcaccgagcgggagttccccgtcttctgcgccttcctggtggaacccattcagggcgagggcggaatccgggtgctttcccgggaattcgccggggaaatccagaagttctgcgagtcgatcgactgccccgttgtcgtcgacgagatccagagcggaatgggcaggaccggaacacttctcgccagtgccccgctcggactgcgcggcgactatttcaccctcgccaagacactgggcggcggaatcgccaagacatccgtgatgctggtgcgcgagaagtattaccggaaggaattcgagatcgtccacagctcgaccttcgccaaggacagcttctcctgtcatatcgccctcaaggtcctcgaactgctggaggccgacggcggacgggcctaccgggccgcggaggagcgcggcgcggccctgcgggcggcgctggacggcgtacgggcggacttccccgacgtcgtcggagacgtccggggccggggcctgatgctgggcctcgaattcctggaccagtcggggtcgtcgtccccggtgatccaggagatcgcggtcggcgggttcttcggctacgtcctcgccgggcatctgctgcgccgccaccgggtgcgtacatttcctacggcgagcgcggtcaacaccctgcgcttcgagccttccatcgagctgaccgacgccgagatcgcccagctcgacgccgggctgcgcgatgtgtgcgagctgctcagggccggggcgggcgaagcgctgaccgcggcctga |
| *TE1 (DEBS TE)* | gactccggtacgcctgcacgtgaagctagctctgcactgcgtgacggctaccgccaagcgggtgtgagcggccgtgttcgtagctacctggatctgttggcaggtctgtcggatttccgcgaacattttgatggtagcgatggttttagcctggatctggttgatatggcagatggtccgggtgaggtgaccgttatttgctgcgcgggtacggctgcgatctctggtccgcatgagttcacccgtctggcaggtgccctgcgcggcattgcacctgttcgtgcggtgccgcagccgggttacgaagaaggcgaaccgctgccttctagcatggcggctgttgcagctgtgcaggctgatgctgtcattcgtacgcaaggcgataagccgttcgtggttgcgggtcactcggcaggcgcgctgatggcgtacgcgctggcgaccgaactgctggatcgtggtcacccgcctcgcggtgttgtcctgattgatgtgtacccgcctggtcatcaggacgcgatgaacgcgtggctggaggaactgacggcaacgttgttcgaccgtgaaactgttcgtatggacgacacccgtctgaccgcgttgggtgcttacgaccgcctgacgggtcaatggcgtcctcgcgaaaccggtttgccgaccctgttggttagcgcgggtgaaccaatgggcccgtggccggacgatagctggaaaccgacctggcctttcgagcacgacaccgttgcggtcccgggtgatcatttcacgatggttcaagaacatgctgatgcgattgcccgtcacattgacgcctggctgggtggcggcaacagctga |
| *TE2 (pik TE)* | gccgagcggaactgggccgtcgccgagccgtcggatcacgagcaggcggaggaggagaaggccgccgctccggcgggggcccgctccggggccgacaccggcgccggcgccgggatgttccgcgccctgttccggcaggccgtggaggacgaccggtacggcgagttcctcgacgtcctcgccgaagcctccgcgttccgcccgcagttcgcctcgcccgaggcctgctcggagcggctcgacccggtgctgctcgccggcggtccgacggaccgggcggaaggccgtgccgttctcgtcggctgcaccggcaccgcggcgaacggcggcccgcacgagttcctgcggctcagcacctccttccaggaggagcgggacttcctcgccgtacctctccccggctacggcacgggtacgggcaccggcacggccctcctcccggccgatctcgacaccgcgctcgacgcccaggcccgggcgatcctccgggccgccggggacgccccggtcgtcctgctcgggcactccggcggcgccctgctcgcgcacgagctggccttccgcctggagcgggcgcacggcgcgccgccggccgggatcgtcctggtcgacccctatccgccgggccatcaggagcccatcgaggtgtggagcaggcagctgggcgagggcctgttcgcgggcgagctggagccgatgtccgatgcgcggctgctggccatgggccggtacgcgcggttcctcgccggcccgcggccgggccgcagcagcgcgcccgtgcttctggtccgtgcctccgaaccgctgggcgactggcaggaggagcggggcgactggcgtgcccactgggaccttccgcacaccgtcgcggacgtgccgggcgaccacttcacgatgatgcgggaccacgcgccggccgtcgccgaggccgtcctctcctggctcgacgccatcgagggcatcgagggggcgggcaagtag |
| *TE3 (rim TE)* | gccgcacagcccggcctgggcggcggcgagggcggcctcctgctggccgggcccgaccccgactcgctggagcggatgttcctcgacgcgatggacgagggcaggttcccggagatccggctcatgctgcgggccctgaccggcctgcgcacgaccttcgagaacacggccgaactggtggagctgccgctgccgaccacgctcgccgagggccccgccgagccgcggctgatctgcatcagcacacccaccgccaacggcggtgtccacgaatacgccaggttcgccgcgtccttccggggcgaacggcacgtcagcgcactgccgctggtcggcttcgccgccggggaacggctgcccgccacacccaccagcgcggtcaggaccatcgcggagagcgctctgcgcgcaagcgacggcaaccccttcatcctggtcggacactcctccggcggtgcgtacgcctacgtggccgccgggctgctggagagcacctggggcatcaagcccgaggccgtggtgctgctggacaccgtcagcatccggcacgacgaccgggaggacatcgactacgacaacctgatgcaccgcaacttcctcgccgaccaggaatccccggtgcgggtgaccaactcccggctgtccgggatggggcggtacatgggcctgctcgggcagctcgacgtccagcacaccagcgctcccgtgctgatcgtccgcgccgcccaggagaccttcgccatggagggcaccgcggccgtctcgcccgacgacacgaagagcgacatgctcccgtcggccgatgtccgcatcgtcgacgcggaccacttctccatggtccgcgaccacgccccgcagaccgcgcagatcgtcaaggactggctggccaccctcggccaggcatga |
| *MaCAR* | atgacggagacgatctccaccgccgccgtgcccaccaccgacctggaggaacaggtcaagcgccggatcgagcaggtggtttcgaacgacccgcagctggcggcgctgctgccggaggactcggtcaccgaggccgtcaacgagccggacctgccgctggtcgaggtcatccgccggctgctggagggctacggcgatcgccccgcgcttggccagcgggccttcgagttcgtcaccggcgacgacggtgccaccgtgatcgcgctgaagcccgagtacacgaccgtctcctaccgggaactgtgggagcgggcggaggcgatcgccgcggcctggcacgagcagggcatccgtgacggggacttcgtcgcccagctcggcttcacctcgacggacttcgcctcgctcgacgtcgcgggcctgcgcctgggcacggtcagcgtgccgctgcagaccggcgccagcctccagcagcgcaacgccatactggaggagacccgcccggccgtgttcgcggcgtccatcgaatacctggacgcagccgtggacagcgtcctcgcgacgccctccgtgcgtctgctctcggtcttcgactaccacgccgaggtcgactcccagcgggaggcgctcgaagcagtccgcgcccggctcgagagcgccggccgcaccatcgtcgtcgaggccctcgccgaggcactggcccgcggccgggacctccctgccgcgcccctgccctcggcggaccccgacgccctgcgcctcctcatctacaccagcgggagcacaggtacgccgaagggcgcgatgtacccgcagtggctggtggccaacctgtggcagaagaagtggctcaccgacgacgtgatcccgtcgatcggagtgaacttcatgccgatgagccacctggccgggaggctgaccctgatgggcaccctcagcggcgggggaacggcctactacatcgcctcctccgacctctccaccttcttcgaggacatcgcgttgatccgaccgtccgaggtgctgttcgtgccccgggtggtggagatggtcttccaacgcttccaggccgagctggaccggtccctcgcccccggcgagtccaacagtgagatcgccgaacgcatcaaggtccggatccgcgagcaggacttcggcggccgcgtcctgtccgcgggttcggggtcggcgccgctgtccccggagatgaccgagttcatggagagtctgctccaggtccctctgcgcgacggctacgggagcacggaggcgggtggggtgtggcgggacggcgtcctgcagcggccgccggtcacggactacaagctggtggatgtacccgagctgggctacttcaccaccgactccccgcacccgcgcggtgagttgcggctgaagtcggaaacgatgttcccgggctactacaagcggccggagacgactgccgacgtcttcgacgacgagggctactacaagaccggcgacgtcgtcgccgaactcggccccgatcacctgaagtatctggaccgggtgaagaacgtgcttaagctggcccagggcgagttcgtcgccgtctccaagctggaggccgcctacacgggaagccccctggtgcggcagatcttcgtgtacggcaactccgagcgttcgttcctcctcgcggtggtggtgcccacccccgaagtgctggagcgctacgccgacagcccggacgcgctcaagccgctgatccaggactcgctgcagcaggtggccaaggacgccgagctccagtcctacgagatcccccgggacttcatcgtcgagacggtgcccttcaccgtggagtccggcctgctctccgacgcccgcaagctgctgaggccgaagctgaaagaccactacggggagcggctcgaggcactgtacgcggagctggcggagagccagaacgagcgactgcgccaactggcgcgcgaggccgccacccgccccgtcctggagaccgtcaccgacgccgctgcggcgctgctgggggcgtcctcgagcgacctcgcccccgacgtccgcttcatcgacttgggcggagactccctgtcggccctctcgtacagcgaactcctccgcgacatcttcgaggtcgacgtcccggtgggggtcatcaactccgtcgccaacgacctcgccgccatcgcccggcacatcgaggcgcagcgcacgggtgccgccacccagccgaccttcgcgtccgtccacggcaaggacgcaaccgtgatcaccgctggcgagctgaccctcgacaagttcctggacgaatcgctgctcaaggcggcgaaggacgtccagcccgccaccgcggacgtcaagaccgttctcgtcacgggcggcaacggctggctgggccgctggctcgtgctggactggctcgagcggctggcgccgaacggcggcaaggtgtatgcgctcatccgcggcgcggacgccgaagccgcgcgggcccgcctcgacgccgtctacgagtccggagacccgaagctgtccgcccactaccggcaactggcccagcagtcgctggaggtgatcgccggggacttcggcgaccaggacctcggcttgtcgcaggaggtctggcagaagctcgccaaggacgtcgacctgatcgtccactccggcgccctggtcaaccacgtcctcccctactcccagctgttcgggcccaacgtcgccggcaccgccgagatcatcaagctcgcgatctccgagcggctcaagcccgtcacgtacctgtcgacggtgggcatcgccgaccagatccccgtgaccgagttcgaggaggactcggatgtgcgggtcatgagcgcggaacgtcagataaacgacggatacgccaacggctacggcaactccaagtgggcgggagaggtgctgctgcgcgaggcgcacgacctggccggtctgccggtccgcgtcttccgcagcgacatgatccttgcgcacagcgactaccacggccagctgaacgtcaccgacgtgttcacccggtccatccagagcctgctcctcaccggcgtggccccggcatcgttctacgagctggacgccgacggcaaccgccagcgagcccattacgacggggtacccggcgacttcacggcagccagcatcaccgcgatcggcggggtgaacgtggtcgacggctaccgctccttcgacgtcttcaacccgcaccacgacggcgtctcgatggacaccttcgtggactggctgatcgacgcggggtacaagatcgcgcggatcgacgactacgaccagtggctggcccgcttcgaactggcgctgaaagggctacccgagcagcagagacagcagagcgtgctcccgctgctgaagatgtacgagaagccgcagccggccatcgacggctcggctctgcccacggccgagttctcccgcgccgtgcacgaggcgaaggtcggcgacagtggcgagatcccgcacgtgaccaaggagctgatcctgaagtacgcctccgacatccagctcctgggtctcgtctaa |
| *SrCAR* | atgaccgagtcccaaagctacgagacccggcaggcgcgaccggccggccagtcgctggccgaacgcgtcgcccggctcgtcgcgatcgacccccaggccgcggcggcagtgcccgacaaggccgtggcggaacgtgcgacccagcagggcctgcggctggcgcagcgcatcgaggccttcctgtcgggctacggcgaccggcccgcgctggcccagcgcgccttcgagatcacaaaggatccgatcaccggtcgggccgtggcgacgctgctgccgaagttcgagaccgtctcctaccgggagttgctggagcggtcccacgcgatcgcgtccgagctggccaaccacgccgaggcccccgtcaaggcgggggagttcatcgcaaccatcggcttcaccagcacggactacacctccctcgacatagcgggcgtactcctggggctcacctcggtgcccctccagaccggcgccaccaccgacacgctcaaggccatcgccgaggagacggccccggccgtcttcggcgcatcggtcgagcacctcgacaacgccgtcaccaccgccctcgccacgccgagcgtccgccggctgctggtgttcgactaccgccagggcgtcgacgaggaccgcgaggcggtcgaagcggcccgcagccgcctcgccgaggcggggagcgcggtgctggtggacaccctcgatgaggtcatcgctcgtggccgcgcgctcccgcgggtggccctgccccccgccaccgacgccggtgacgactccctgtctctactgatctacacgtccggcagcacggggacgcccaagggcgcgatgtacccggagcgcaacgtcgcccagttctggggcggcatctggcacaacgcgttcgacgacggcgactcggccccggacgtgccggacatcatggtcaacttcatgccgctgtcgcacgtcgccgggcggatcggactgatgggcaccctgtcctcggggggcacgacctacttcatcgccaagagcgacctgtccaccttcttcgaggactactccctcgcgcgccccaccaagctgttcttcgtgccgcgcatctgcgagatgatctaccagcactaccagtccgaactggaccggatcggtgccgccgacggctcgccgcaggccgaggccatcaagaccgagttgcgcgagaagctcctcggcggaagggtgctcacggcggggtcgggaagtgcaccgatgagccccgagctgaccgccttcatcgagtcggtcctccaggtccacctggtggacggctatggatcgaccgaggccgggccggtgtggcgggaccgcaagctcgtcaagccgcccgtcaccgagcacaagctgatcgacgtccccgaactgggttacttctccacggactcgccctacccgcgcggcgagctggccatcaagacgcagaccatcttgcccggctactacaagcgcccggagacgaccgccgaggtcttcgacgaggacggcttctacctcaccggcgacgtggtcgccgaggtcgcgcccgaggagttcgtatacgtcgaccgccgcaagaacgtcctgaagctgtcgcagggcgagttcgtcgcgctgtccaagctcgaagccgcctacggcacctccccgctggtccggcagatctccgtctacggctccagccaacgctcgtacctgctcgccgtggtcgtgcccaccccggaggcgctcgcgaagtacggggacggcgaggcggtgaagtccgcgctgggcgacagcttgcagaagatcgcccgcgaggagggccttcagtcctacgaggtgccccgcgacttcatcatcgaaaccgacccgttcaccatcgagaacggcatcctgtccgatgcgggcaagaccctgcgccccaaggtcaaggcccggtacggcgagcggctggaggccctgtacgcccagctcgccgagacccaggccggggagctgcgtagcatccgggtcggcgcgggggaacggcctgtcattgagaccgtgcagcgggcagctgccgcgctcctcggcgccagcgccgcggaggtcgaccccgaagcccacttcagcgacctcggcggcgactccctctccgcgctgacctactccaacttcctgcacgagatcttccaggtggaggtgccggtctccgtgatcgtcagcgcggccaacaacctccggtcggtggccgcacacatcgagaaggagcgcagctccggctccgaccggccgaccttcgcctccgtgcacggggcgggcgccaccacgatccgtgcctcggacctgaaactggagaagttcctcgacgcccagacgctggcggcggccccgtcgctgccgaggccggcgagcgaggtgcgcacggtgctgctgactggttctaacggctggctcggccgtttcctggcactggcctggctggagcggctggtgccgcagggcgggaaggtggtcgtgatcgtccgcggcaaggacgacaaagccgccaaggcccgcctcgactccgtgttcgagtccggcgacccggcactgctcgcgcactacgaggaccttgccgacaagggcttggaagtgctcgccggggatttctccgacgcggacctgggactgcgtaaggccgactgggaccggctggccgacgaggtggacctgatagtgcacagcggcgccctcgtcaaccacgtcctcccgtacagccagctgttcggccccaacgtggtcggcaccgccgaagtcgccaagctggcgctcacgaagcgcctgaagccggtcacctacctgtccaccgtcgcggtggcggtcggcgtcgagccgtccgcattcgaggaggacggcgacatccgcgacgtgtcggccgtccggtcgatcgacgagggctacgccaacggttacggcaactccaagtgggccggcgaggtcctgctgcgcgaggcctatgagcacgcgggcctgccagtgcgcgtcttccgctccgacatgatcctggcccaccgcaagtacaccgggcagctgaacgttcccgaccagttcacgcggctcatcctcagcctgctcgcgacgggcatcgcccccaagtcgttctaccagctggacgccaccggtggccggcagcgggctcactacgacgggatccccgtcgacttcaccgcggaggcgatcaccacgctcggcctcgccggaagcgacggctaccactccttcgacgtcttcaacccgcaccacgacggtgtaggtctagacgagttcgtcgactggctggtcgaggcgggccacccgatctcccgggtcgacgactacgcggagtggctgtcgcggttcgagaccagcctcagggggctgcccgaggcccagcgccagcacagcgtgctgccgctcctgcacgcgttcgcccaaccggcccccgccatcgacggctcgcccttccagaccaagaacttccagtcgagtgtgcaggaggcgaaggtgggggccgagcacgacatcccgcatctggacaaggcgctgatcgtcaagtacgcggaagacatcaagcagctcggcctgctc |
| *MmCAR* | atgtccccgatcacccgcgaggagcggctcgagcgccgcatccaggacctctacgccaacgacccgcagttcgccgccgccaagcccgccaccgcgatcaccgcggccatcgaacgccccggcctgccgctgccccagatcatcgagacggtgatgaccggctacgccgaccggcccgccctggcccaacgctccgtggagttcgttaccgacgccggcaccggccacaccaccctgcgtctcctgccgcacttcgagactatcagttacggcgagctctgggaccggatctcagctctggcagatgtgctgtcgaccgagcagaccgtgaagccgggagaccgcgtctgcctgctgggcttcaactccgtcgactacgcgacgatcgacatgacgctggcgagactgggtgccgtcgccgtgccgctccagacgagcgccgccatcacccagctgcagcccatcgtcgccgagacccagcctaccatgatcgcggccagcgtggacgccctcgccgacgcgaccgaactggccctctccgggcagaccgccacccgggtcctcgtgttcgaccaccaccggcaggtggacgcccaccgcgcggcggtcgagtccgcccgggagcggcttgcagggtcggccgtcgtcgagaccctcgcggaagcgatcgcccgcggcgacgtcccccggggggcctcggcggggtcggcccccggcaccgacgtgtcggacgacagcctcgcactgctcatctacacctccggctccacgggtgcgccgaagggcgcgatgtacccgcggcggaacgtcgcgacgttctggcgcaagcgcacctggttcgagggcgggtacgagccctccatcacgctcaacttcatgccgatgagccacgtcatgggccgccagatcctgtacggaacgctgtgcaacggcggcacggcgtacttcgtggccaagtccgacctgagcaccctgttcgaggacctggcgctggtgcggcccaccgagctgaccttcgtcccgcgcgtctgggacatggtcttcgacgagttccagtccgaggtcgacaggcgtctggtcgacggcgcggaccgggtggccctggaagcgcaggtcaaggccgagatccgcaacgacgtgttgggcggtcgctacacgtcggccctcacaggctcggcgccgatcagcgacgagatgaaagcgtgggtcgaggagctcctcgacatgcacctcgtcgagggctacggctccacggaggccggcatgatcctgatcgacggcgcgatccgacggcccgccgtgctcgactacaagctggtcgacgtcccggacctgggctacttcctgacggaccggccgcacccccggggagaactgctggtcaagaccgactcgctgttcccgggctactaccagcgcgccgaggtgaccgcggacgtattcgacgcggacgggttctaccggaccggggacatcatggccgaggtcggcccggaacagttcgtctatctggaccgccggaacaacgtcctcaagctctcgcagggcgagttcgtgaccgtctccaagctggaggcggtcttcggcgacagtccgctggtgcggcagatatacatctacggtaactcggcgcgggcctacctgctcgccgtcatcgtccccacgcaggaggcactcgacgcggtgcccgtcgaggagctgaaggcccggctgggcgattcgctccaggaagtggccaaggccgccggactccagtcctacgagatcccccgcgacttcatcatcgagaccacgccctggaccctggagaacggcctgctgaccggcatccgcaagctcgctcgcccgcagctcaagaagcactacggtgaactgttggagcagatctacaccgacctggcgcatggccaggccgacgagctgcgctccctcaggcagagcggcgccgacgccccggttctggtcacggtctgccgcgctgccgcggcgctcctcgggggcagcgcgtccgacgtgcaacccgacgcccacttcaccgacctgggcggggactccctgtcggccctgtccttcaccaacctgctgcacgagatcttcgacatcgaggtcccggtgggagtgatcgtgtccccggccaacgatctccaggcgctcgccgactacgtcgaggccgcccgcaagccggggagcagccgtcccaccttcgcctccgtccacggtgcctcgaacggccaggtgacggaggtgcacgccggcgacctcagcctggacaagttcatcgacgcggccacgctggccgaggcgccgcgcctgcccgccgccaacacccaggtccgcaccgtgctgctgacaggggcgaccggcttcctcggccgctacctcgcgctggagtggctggagcggatggacctcgtcgatggcaagctgatctgcctggtgcgggcgaagagcgacacggaagccagggcgcgtctggacaagaccttcgactccggggaccccgagctgctggcgcattaccgggcgctggcgggcgaccacctggaggtcctggccggtgacaagggcgaggcggacctcggactcgaccgacagacctggcagcgcctcgccgacaccgtcgacctcatcgtggacccggccgcgctggtgaaccacgtgctcccctacagccagctgttcggcccgaacgccctcggcaccgccgaactcctgcggctcgccttgacgtcgaagatcaagccgtactcctacacctccaccatcggcgtcgccgaccagatcccgcccagcgcgttcaccgaggacgcggacatccgggtgatctccgccacgagagccgtggacgactcgtacgccaacggctactccaactccaagtgggcgggggaggtgctgctccgcgaggcgcacgacctgtgcggcctgccggtggcggtcttccggtgcgacatgatcctcgccgataccacctgggccggccagctcaacgtgcccgacatgttcacccgcatgattctgtccctggcggccaccggcatcgcccccggctccttctacgagctggccgccgacggggcgcggcagcgcgcccactacgacggtctgcccgtcgagttcatcgcggaggccatctcgacgctgggcgcccagtcgcaggacggcttccacacctaccacgtcatgaacccgtacgacgacggtatcggactcgacgagttcgtcgactggctcaacgagagcggctgccccatccagcgcatcgccgactacggcgactggctgcaacggttcgagacggccctgcgcgccctgccggaccggcagcggcacagctcgctgctgcccctgctgcacaactaccgccagccggagcgcccggtccgcggctcgatcgctccgacggaccgcttccgcgccgccgtccaggaggccaagatcgggcccgacaaggacatcccgcacgtcggcgcgccgatcatcgtcaagtacgtatccgacctgcgcctgctcgggctcctc |
| *YqhD* | atgaacaacttcaacctgcacacgcccacccgcatcctgttcggcaagggcgccatcgcgggactccgcgagcagatcccgcacgacgcgcgggtgctgatcacgtacggcggcgggagcgtcaagaagacgggcgtcctcgaccaggtcctggacgccctgaagggcatggacgtgctggagttcggcggcatcgagcccaatccggcctacgagaccctgatgaacgcggtgaagctggtccgcgaacagaaggtcaccttcctgctggcggtgggaggcggcagcgtcctcgacggaacgaaattcatcgctgccgccgcgaactacccggagaacatcgacccctggcacatcctccagaccgggggcaaggaaatcaagtcggcgatcccgatgggctgcgtgctcaccctgccggccaccggtagcgagtccaacgccggggcggtcatctcccgcaagaccaccggcgacaagcaggcgttccactccgcccacgtccagcccgttttcgcggtgctggacccggtctacacctacacgctcccgccccggcaggtggccaacggtgtcgtcgacgccttcgtgcacaccgtcgagcagtacgtgaccaagccggtggacgcgaagatccaggaccggttcgccgaagggatcctcctgacgctcatcgaggacgggccgaaggcgctgaaggagccggagaactacgacgtgagggccaacgtcatgtgggccgccacccaggcccttaacggcctgatcggcgccggcgtgccgcaggactgggcgacccacatgctgggccacgagctgaccgccatgcacggcctcgaccacgcccagacactggccatcgtcctgcccgcactgtggaacgagaagcgcgacaccaagcgggccaagctgctgcagtacgccgaacgggtctggaacatcaccgagggctccgacgacgagcgaatcgacgccgccatcgcggcgacgcgcaacttcttcgagcaactgggtgtacccacgcatctgtcggactacgggctggatggctcgtccatacccgcgctcctcaagaagctcgaggagcacgggatgacccagctcggcgagaaccacgacatcaccctcgacgtctcgcgccggatctacgaggcggcccgt |
| *ChnD* | atgcactgctactgcgtgacccaccacgggcagccgctcgaggacgtcgagaaggaaatcccccaacccaagggcaccgaggtcctgctgcacgtcaaggcggccggactgtgccacaccgatctgcacctgtgggagggctactacgacctgggcggcggcaagcggctgagcctcgccgaccgggggctaaaaccgccgctgaccctctcccacgagatcaccggtcaggtcgtcgccgtcggcccggacgcggagtccgtgaaggtgggcatggtctccttggtgcacccctggatcggctgcggggagtgcaactactgcaagcgcggcgaggagaacctgtgtgcgaagccgcagcagctcggcatcgccaagcccgggggcttcgccgagtacatcatcgtcccgcacccgcgctacctggtggacatcgccgggctggacctggccgaggcagcgccgctggcctgcgccggcgtcaccacgtacagcgcgctcaagaagttcggcgacctcatccagtccgagccggtcgtgatcatcggcgcgggcggactcgggctcatggccctggagctgctcaaggccatgcaggcgaagggcgcgatcgtggtcgacatcgacgactccaagctcgaggccgcccgcgcggcgggcgccctctcggtcatcaactcgcgttcggaggacgccgcccagcagctgatccaggccaccgacggtggagctcggctgatcctcgacctcgtcgggagcaaccccaccttgtccctcgcgctggcgtccgcggcccgggggggtcacatcgtgatctgcggcctgatgggcggcgagatcaagctgtcgatccccgtgataccgatgcgccccctgacgatccagggctcctacgtcggcacggtggaggagctgcgcgaactggttgaactcgtgaaggagacgcacatgtcggccatccccgtcaagaagctgccgatcagccagatcaacagcgccttcggcgacctgaaggacggcaacgtcatcggccggatcgtgctgatgcatgagaactaa |
| *YahK* | atgaagatcaaggcggtcggcgcctactcggccaagcagcccctcgaaccgatggacatcacgcggcgggagcccggcccgaacgacgtcaagatcgagatcgcgtactgcggcgtgtgtcactccgacctgcaccaggtgcgctccgagtgggcgggcaccgtctacccgtgcgtgcccggtcacgagatcgtcggccgggtcgtcgcggtaggcgaccaggtcgagaagtacgcccccggagacctcgtgggggtgggctgcatcgtcgactcctgcaagcactgcgaagagtgcgaggacggcctggagaactactgcgaccacatgaccggcacctataactccccgacgccggacgaacccggacacaccctgggcggctacagccagcagatcgtggtgcacgagcgctacgtcctgcgcatccgccacccgcaggagcagctcgccgccgtggcgccgctgctgtgcgcgggcatcaccacgtactcgccgctgcggcactggcaggccggccccggcaagaaggtgggcgtcgtcgggatcggtggcctcggccacatgggcatcaagctcgcgcacgccatgggcgcgcatgtcgtggccttcacgaccagcgaggcgaaacgcgaggcggccaaggccctgggggcggacgaggtggtcaacagccgcaacgccgacgaaatggctgcccacctgaagtccttcgacttcatcctgaacacggtcgccgcaccccacaacctggacgacttcaccaccctcctcaagcgtgacgggaccatgacgctggtgggtgccccggccactccgcacaagtcgccggaggtcttcaacctgatcatgaagcggagggcgatcgccgggtcgatgatcgggggaatccccgagacccaggagatgctcgacttctgcgccgagcacggcatcgtggccgacatcgagatgatccgggccgatcagatcaacgaggcctacgagcggatgctgcgcggcgacgtcaagtaccgcttcgtcatcgacaaccgcaccttgaccgactaa |
| *YjgB* | atgtccatgatcaagtcgtacgccgcgaaggaggccggaggcgagctggaggtctacgagtacgaccccggggaactgcggccgcaggacgtcgaggtgcaggtcgattactgcgggatctgccactccgacctgtcgatgatcgacaacgagtggggcttctcgcagtacccgctggtcgcggggcacgaggtcattgggcgggtggtggccctcggcagcgcggcccaggacaagggcctccaggtgggccagcgggtcggcatcggctggacggcccgctcgtgtggacactgcgacgcttgcatctccggcaaccagatcaactgcgagcagggcgcggtgccgacgatcatgaaccgcggcgggttcgccgagaagctccgtgcggactggcagtgggtgatccccctgcccgagaacatcgacatcgagagcgcgggtccgctgctgtgcggcggcatcacggtcttcaagccgctcctcatgcaccacatcaccgcgacgtcccgcgtgggcgtcatcggcatcggggggctgggccacatcgcgatcaagctcctgcacgcgatgggctgcgaggtgaccgccttctcgtccaacccggccaaggagcaggaggtcctcgccatgggtgccgacaaggtggtcaactcccgggacccgcaagccctgaaggcgctggccggccagttcgacctcatcatcaacaccgtgaacgtctccctggactggcagccgtacttcgaggcactcacctacggcggcaacttccacaccgtcggcgccgtcctgacccccctgtccgtccccgcgttcaccctgatcgccggcgaccggagcgtgagcggtagtgccaccggcaccccctacgaattgcgcaagctgatgcgcttcgcggccaggtcgaaagtcgcgcccacgaccgagctgttcccgatgagcaagatcaacgacgccatccagcacgtacgcgacggcaaggcccggtaccgcgtcgtgctcaaggccgacttctaa |
| *ScADH2* | atgtcgatccccgagacccagaaggccatcattttctacgaatcgaacggcaagctggagcacaaggacatcccggtgccgaagcccaaaccgaacgaactgctgatcaacgtcaagtacagcggcgtctgccacaccgacctgcacgcctggcacggggattggcccctgccgacgaagctcccgctggtgggtgggcacgagggcgccggcgtggtggttggcatgggtgagaatgtgaagggctggaagatcggcgactacgcgggcatcaagtggctgaacgggagctgcatggcgtgtgagtactgcgagctcggcaacgagtccaactgcccgcacgcggacctgtccgggtacacccacgacggctccttccaggagtacgcgaccgccgacgccgtccaggccgcgcacatcccgcagggcacagacctcgccgaggtggcgccgatcctgtgcgccgggatcaccgtctacaaggcgctcaagtcggccaacctgcgggccggccactgggcggccatcagcggcgcggctggcggtctcggaagcctggccgtgcagtacgccaaggccatgggctaccgcgtcctgggcatcgacggggggccgggaaaggaggagctgttcacgtcgctcggcggcgaggtgttcatcgacttcaccaaggagaaggacatcgtgtcggcggtcgtgaaggccaccaacggcggcgcccatggcatcatcaacgtctccgtcagcgaagcggcaatcgaggcctccacgcgttactgccgggcgaacggcaccgtggtcctcgtcggcctccccgcgggcgcgaagtgctcctccgacgtcttcaaccacgtcgtcaagtccatctcgatcgtcggctcctatgtgggcaaccgcgccgacacccgcgaggccctggacttcttcgcccgggggctcgtgaagagtcccatcaaggtcgtaggtctgagctcgctgcccgagatctacgagaagatggagaagggccagatcgccggccgctacgtcgtcgacacgtccaagtaa |

### **Supplementary Table 4. Statistics for data collection and refinement of CpkC TR**

|  | **CpkC TR** |
| --- | --- |
| ***Data collection*** |  |
| Wavelength (Å) | 0.97741 |
| Resolution range (Å) | 44.76 - 1.84 (1.88 - 1.84) |
| Detector Distance (mm) | 220 |
| Φ (deg.) collected / ΔΦ (deg.) | 180/0.25 |
| Exposure time (seconds) | 0.3 |
| Temperature of collect (Kelvin) | 100 |
| ***Data statistics*** |  |
| Space group | P 21 21 21 |
| Unit-Cell parameters (Å) | a=49.63 b=103.55 c=148.32 |
| Total reflections | 445567 (44265) |
| Unique reflections | 67220 (4891) |
| Multiplicity | 6.6 (6.7) |
| Data completeness (%) | 100 (99.7) |
| <I/σ(I)> | 12.5 (0.9) |
| R_merge_ (%)  R_pim_ (%) | 0.095 (1.90)  0.059 (1.20) |
| CC_1/2_ | 0.999 (0.358) |
| ***Structure Refinement*** |  |
| Reflections used in refinement | 67220 (4891) |
| Reflections used for R_free_ | 1828 (151) |
| R_factor_ | 0.183 (0.338) |
| R_free_ | 0.218 (0.352) |
| RMS from ideal geometry |  |
| Bond lengths (Å) | 0.008 |
| Bond angles (º) | 0.77 |
| Average B-factor | 35.9 |
| Macromolecules | 35.2 |
| Ligands | 37.0 |
| Solvent | 38.6 |
| Ramachandran Plot |  |
| Favoured region (%) | 97.5 |
| Outliers region (%)  Rotamer outliers (%) | 0  0.1 |
| Number of TLS groups | 16 |

### **Supplementary Table 5. Thermodynamic analysis of different synthesis pathways of 2-ethyl-1,3-hexanediol and 2-ethyl-3-hydroxyhexanoic acid**

| Product | Synthesis pathway | Stoichiometric of synthesis pathway | Δ_r_G’^o^ (kcal/mol) |
| --- | --- | --- | --- |
| 2-ethyl-1,3-hexanediol | PKS KS catalysed decarboxylative Claisen condensation-based pathway | Butyryl-CoA + Ethylmalonyl-CoA + 4 H^+^ + 3 NAD(P)H → 2-ethyl-1,3-hexanediol + 2 CoA + CO_2_ + 3 NAD(P)^+^ | -2.78 |
|  | Thiolase catalysed non-decarboxylative Claisen condensation-based pathway | 2 Butyryl-CoA + 3 H^+^ + 3 NAD(P)H → 2-ethyl-1,3-hexanediol + 2 CoA + 3 NAD(P)^+^ | 0.31 |
| 2-ethyl-3-hydroxyhexanoic acid | PKS KS catalysed decarboxylative Claisen condensation-based pathway | Butyryl-CoA + Ethylmalonyl-CoA + H^+^ + NAD(P)H + H_2_O → 2-ethyl-3-hydroxyhexanoic acid + 2 CoA + CO_2_ + NAD(P)^+^ | -7.36 |
|  | Thiolase catalysed non-decarboxylative Claisen condensation-based pathway | 2 Butyryl-CoA + NAD(P)H + H_2_O → 2-ethyl-3-hydroxyhexanoic acid + 2 CoA + NAD(P)^+^ | -4.27 |

a.	Calculated through stoichiometric summations of standard formation Gibbs energies (Δ_f_G’^o^) of each substrate and products of the pathways. Δ_f_G’^o^ were estimated through the group contribution method under standard conditions (298K, pH 7, zero ionic strength)^14^.
